# Synthetic yeast chromosome XI design enables extrachromosomal circular DNA formation on demand

**DOI:** 10.1101/2022.07.15.500197

**Authors:** Benjamin A Blount, Xinyu Lu, Maureen R M Driessen, Dejana Jovicevic, Mateo I Sanchez, Klaudia Ciurkot, Yu Zhao, Stephanie Lauer, Robert M McKiernan, Glen-Oliver F Gowers, Fiachra Sweeney, Viola Fanfani, Evgenii Lobzaev, Kim Palacios-Flores, Roy Walker, Andy Hesketh, Stephen G Oliver, Yizhi Cai, Giovanni Stracquadanio, Leslie A Mitchell, Joel S Bader, Jef D Boeke, Tom Ellis

## Abstract

We describe construction of the 660 kilobase synthetic yeast chromosome *XI* (*synXI*) and reveal how synthetic redesign of non-coding DNA elements impact the cell. To aid construction from synthesized 5 to 10 kilobase DNA fragments, we implemented CRISPR-based methods for synthetic crossovers *in vivo* and used these methods in an extensive process of bug discovery, redesign and chromosome repair, including for the precise removal of 200 kilobases of unexpected repeated sequence. In *synXI*, the underlying causes of several fitness defects were identified as modifications to non-coding DNA, including defects related to centromere function and mitochondrial activity that were subsequently corrected. As part of synthetic yeast chromosome design, loxPsym sequences for Cre-mediated recombination are inserted between most genes. Using the *GAP1* locus from chromosome *XI*, we show here that targeted insertion of these sites can be used to create extrachromosomal circular DNA on demand, allowing direct study of the effects and propagation of these important molecules. Construction and characterization of *synXI* has uncovered effects of non-coding and extrachromosomal circular DNA, contributing to better understanding of these elements and informing future synthetic genome design.

## Introduction

Our rapidly improving understanding of DNA function, along with our ability to design and build large DNA constructs, has led to us being able to create synthetic genomes assembled from chemically synthesized DNA designed *in silico (Gibson et al., 2010*; Fredens *et al*., 2019). Designing and assembling synthetic genomes provides opportunities to assess current understanding of how DNA sequence and structure underpin cellular properties and behavior. Altering a DNA sequence, even when preserving encoded amino acid sequences, can affect how a gene is transcribed (Alper *et al*., 2005), how mRNA is localised (Marc *et al*., 2002), processed (Pleiss *et al*., 2007) and translated (Yu *et al*., 2015), as well as altering spatial localization (Mercy *et al*., 2017), 3D interactions of genomic DNA (Duan *et al*., 2010), and interactions with nuclear DNA-associated proteins (Garvie and Wolberger, 2001). Changes predicted to have no functional effect lead to unexpected phenotypes, an opportunity to uncover the underlying cause and refine our understanding arises. When this process occurs at a genomic scale, the scope for learning more about how DNA functions within a cell, from the base pair to genome level, is considerable.

The synthetic yeast genome project, Sc2.0, is an international collaboration to build the first eukaryotic synthetic genome, that of *Saccharomyces cerevisiae*. The Sc2.0 genome contains many design features that probe eukaryotic genome biology and will yield strains with encoded abilities not found in nature. Perhaps the most immediately useful of these features is the incorporation of loxPsym recombinase target sites in the 3’ untranslated region (UTR) of almost all nonessential genes as well as at certain “landmark sites” where elements such as repeated DNAs or tRNA genes have been deleted by design (Dymond *et al*., 2011). These loxPsym sites enable a process of on-demand combinatorial chromosome rearrangement, Synthetic Chromosome Rearrangement and Modification by LoxPsym-mediated Evolution (SCRaMbLE). The diversity of gene content and arrangements generated by SCRaMbLE in a population of cells with Sc2.0 chromosomal DNA is vast (Shen *et al*., 2016), and these synthetic diversified populations also show wide phenotypic variation. By isolating ‘SCRaMbLEd’ cells from a population with phenotypes of interest, synthetic genetic changes resulting in desirable qualities can be identified. These can include properties desirable for biotechnology, such as enhanced growth on an alternative feedstock (Blount *et a*l., 2018), improved product biosynthesis (Liu *et al*., 2018) or resistance to adverse growth conditions (Luo *et al*., 2018). The SCRaMbLE system has also been used as a driver of random gene loss in efforts to determine the content of minimal or reduced eukaryotic genomes (Luo *et al*., 2021).

In addition to the SCRaMbLE system, the Sc2.0 genome has many other design features (Richardson *et al*., 2017). These include gene recoding to remove all instances of the TAG stop codon and incorporating synonymous mutation watermarks, dubbed PCRTags, in all genes of each chromosome. Several types of genetic element are also removed or recoded. Retrotransposon sequences including Long Terminal Repeats (LTRs) have been removed, as have subtelomeric repeats and all introns not previously characterised as being essential. As hotspots for chromosomal instability, tRNA gene sequences are removed from the 16 chromosomes to ultimately be complemented by a new 17th chromosome dedicated to tRNA genes (Schindler *et al*., in preparation). Such widespread changes to the genome are likely to affect cellular processes and phenomena not necessarily captured by general phenotypic screens used to characterize synthetic chromosomes thus far (Annaluru *et al*., 2014). One such phenomenon is formation of extrachromosomal circular DNA (eccDNA).

eccDNA is formed when a section of DNA is excised from a chromosome and forms a circular DNA species in the nucleus. The exact mechanisms for this vary between eccDNAs (Carroll *et al*., 1988; Dillon *et al*., 2015; van Loon *et al*., 1994) but include recombination between repeated sequences, including LTRs (Møller *et al*., 2015). Loci found on eccDNA elements typically contain replicative elements in addition to coding (Møller *et al*., 2015). As replication and segregation of this DNA during cell division is not controlled by the standard chromosomal mechanisms, copy number of the DNA is dysregulated and inheritance is asymmetric (deCarvalho *et al*., 2018). As a result, populations of cells can display high levels of heterogeneity in eccDNA copy number. In humans, the presence of ecDNA (equivalent to eccDNA in yeast) is thought to be a driver of evolution (Ling *et a*l., 2021). Accumulation of ecDNA in the nucleus of cells is also emerging as an important factor in cancer (Paulsen *et al*., 2018; Wu *et al*., 2019), aging (Hull *et al*., 2019) and the stimulation of immune response (Wang *et al*., 2021). eccDNAs found in yeast typically form via recombination at LTRs and other sequences, and yeast eccDNAs are thought to aid in adaptation to challenging environmental conditions (Møller *et al*., 2015). The replacement of LTR sequences with loxPsym recombination sites in Sc2.0 chromosomes gives us a new opportunity to directly study the effects of eccDNAs on the cell.

Here we report the assembly and successful debugging of synthetic chromosome *XI* (*synXI*) of the Sc2.0 synthetic yeast genome. We developed and deployed a range of CRISPR-based approaches to combine sections of synthetic DNA *in vivo*, debug complex growth defects and correct large structural variations. We also demonstrate that the loxPsym formatting of Sc2.0 genetic loci allows on-demand formation of eccDNA species. These molecules show the uneven inheritance patterns of natural eccDNAs and will be a useful tool for future studies of these important molecules.

## Results

### *synXI* Assembly

#### synXI *design and synthesis*

In line with the other synthetic chromosomes generated by the Sc2.0 project, we designed *synXI* following the Sc2.0 design criteria set out in (Richardson *et al*., 2017; **Table 1**). These criteria include the removal of tRNA sequences, repeat DNA, non-essential introns and transposon-associated sequences, the replacement of telomeres with a custom-designed telomere seed sequence and the conversion of all TAG stop codons to TAA codons. We also introduced 457 pairs of PCRTag watermarks and inserted 199 loxPsym recombination sites into the 3’ untranslated region (UTR) of non-essential genes and sites of key design changes.

**Table 1:**
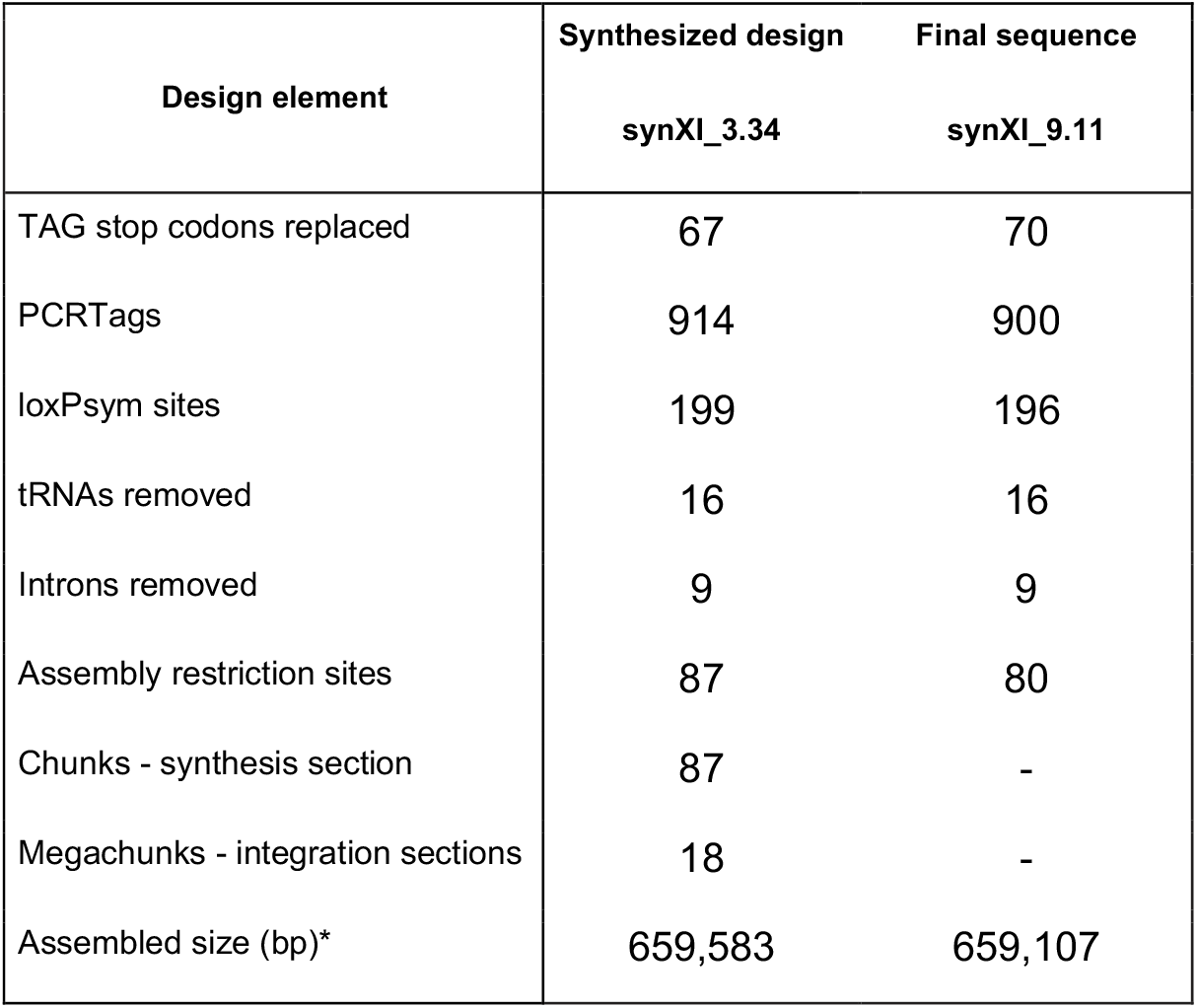
Summary of synthetic chromosome synXI design and final sequence. Asterisk denotes that this number does not take into account in vivo fluctuation in telomere sequence length.

The resulting sequence, *synXI*_3.34, was divided into “chunk” sections, ranging from 4.8 kb to 9.8 kb in size. Each chunk was flanked by recoded recognition sites for restriction enzymes which cleave to generate non-palindromic sticky ends. The chunks were grouped into 18 “megachunks” with a letter designation. Where chunks represent units of DNA synthesis, megachunks represent units of *in vivo* assembly. With the exception of the first 2 chunks, which comprised megachunk A, we assigned chunks into groups of 5 per megachunk, with the final chunk in each megachunk having an auxotrophic marker sequence added to the 3’ end. The resulting synthetic sequence consists of 18 megachunks, A-R, further subdivided into 87 chunks (**Fig 1A**). The total amount of DNA synthesized was 701,706 bp, assembled into a 659,583 bp chromosome. Chunks were synthesized and cloned into bacterial vectors by GenScript Biotech (Piscataway, NJ, USA, chunks A1-C5) and by GeneArt AG (Regensburg, Germany, chunks D1-R5).

**Figure 1:**
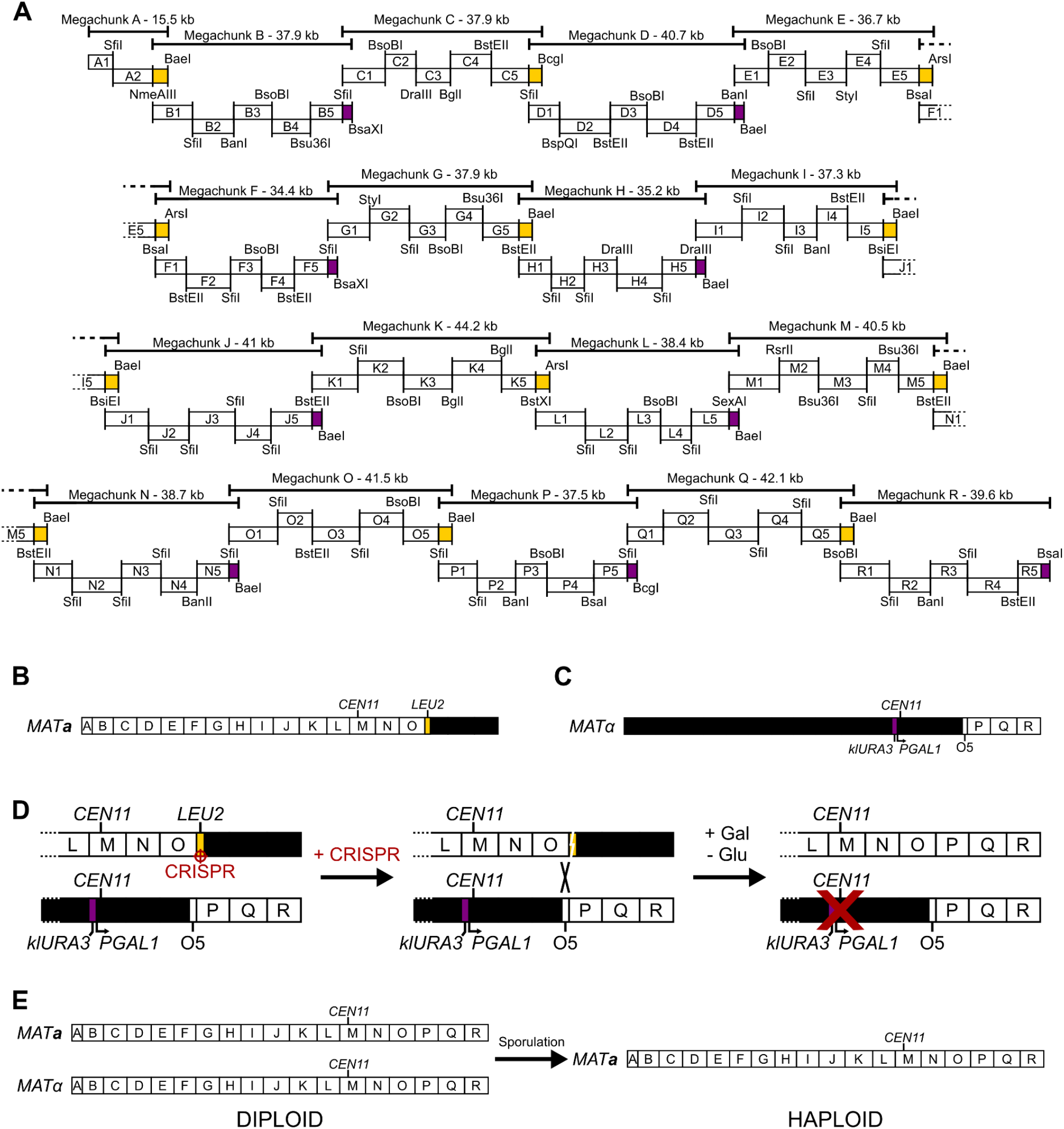
Synthetic chromosome XI design, synthesis and assembly. (A) Schematic overview of the synthetic DNA sections making up synXI_3.34 with megachunk groupings and assembly restriction sites indicated. Purple blocks indicate a URA3 marker gene and yellow blocks indicate a LEU2 marker gene. (B) Topology of the synXI assembly strain ysXIa25 with white lettered boxes representing integrated megachunk sections and black sections representing wild type sequence. (C) Topology of the synXI assembly strain ysXIa30 with white lettered boxes representing integrated chunk or megachunk sections and black sections representing wild type sequence. (D) Overview of the method to consolidate synthetic chromosomal sequences in a diploid cell in vivo, generating a complete synthetic chromosome. (E) Overview of the generation of a haploid strain containing a single copy of the complete synXI.

Due to a non-standard GFF notation of intron-encoding genes, the software platform used for design (Biostudio) did not identify TAG stop codons in these genes and, subsequently, we erroneously included 3 TAG codons in the *synXI*_3.34 design. These TAG codons were in the intron-containing genes *SFT1* (encoded in chunk M1), *ECM9* (chunk M3) and *YKR005C* (chunk M3). These TAG stop codons were subsequently recoded to TAA in design *synXI*_3.36. As DNA synthesis had already been completed at the time of this design update, the existing M1 and M3 chunk plasmids were edited to conform to *synXI*_3.36. A summary of *synXI* versions is shown in **Table 2** and more in-depth descriptions of strains and chromosome versions can be found in **Table S1** and **Table S5**.

**Table 2:**
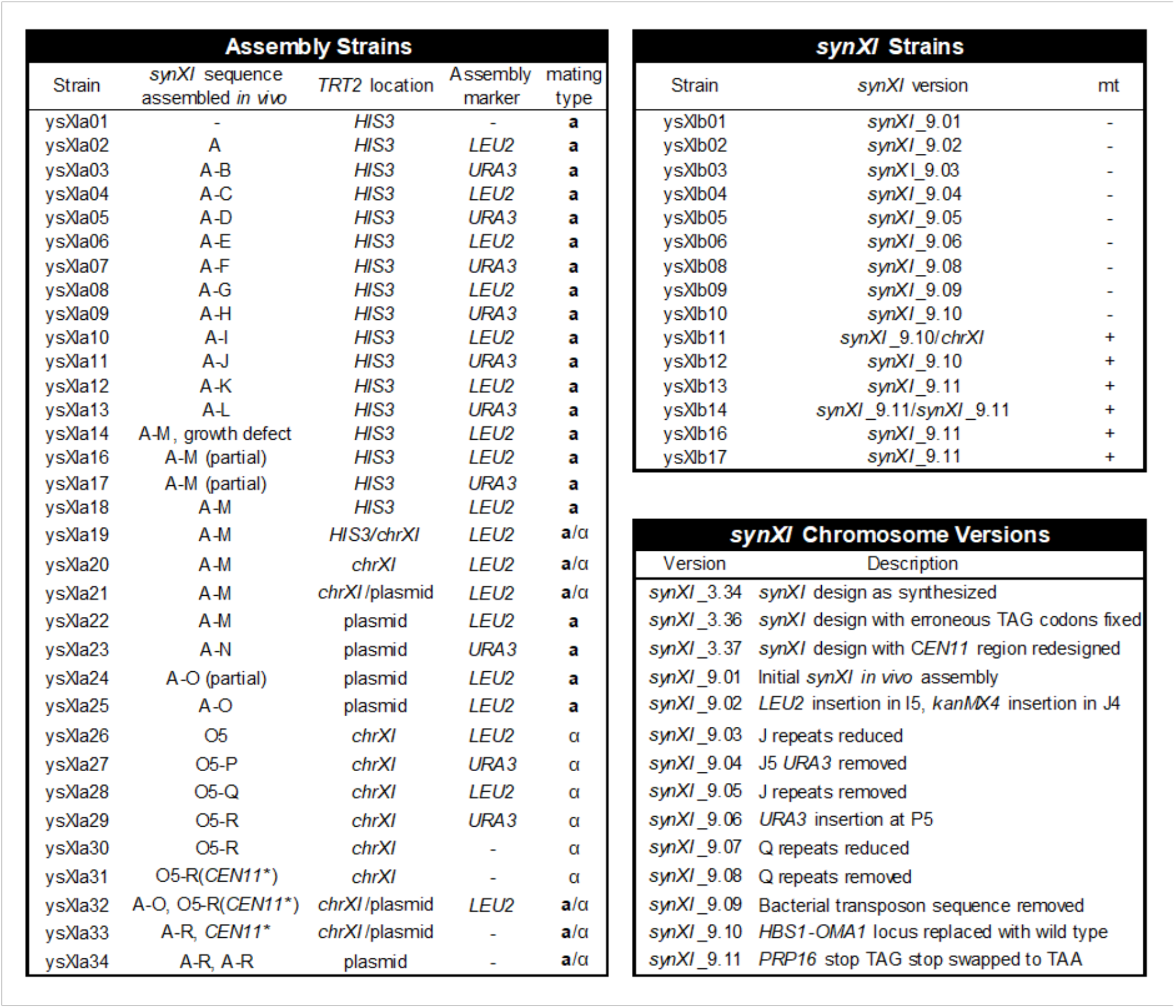
Summary of the strains used to construct and debug synXI and the iterative versions of the synXI chromosome. The “synXI sequence assembled in vivo” column indicates the amount of synthetic sequence successfully integrated to replace chrXI sequence. Letters represent whole megachunks, O5 refers to chunk O5. CEN11* indicates that CEN11 has been replaced with the klURA3-GAL1p-CEN11 construct. The presence or absence of mitochondrial function is given in the “mt” column. More details on strains used can be found in Table S1. More details on synXI versions can be found in Table S5.

#### Generating a recipient strain for construction

A major design feature of Sc2.0 is the removal of all tRNA sequences from the synthetic chromosomes and their incorporation into a tRNA neochromosome. In most cases, tRNA gene removal is compensated by additional genes encoding those tRNAs in other chromosomal loci. This is not the case with unique tRNA genes, for which removal is lethal to the cell. A unique threonine tRNA gene, *tT(CGU)K* (*TRT2*), is present in chromosome XI at a locus replaced by chunk B5. Full integration of megachunk B therefore required complementation of *TRT2* at another locus.

As well as an additional copy of *TRT2*, we required the starting strain for *synXI* construction to have a selectable marker in the chromosome XI (*chrXI*) sequence that would be replaced by the initial megachunk integration event. To address this, we used a derivative of a BY4741 strain with a *kanMX4* marker inserted in *YKL220C* (strain Y07039; Winzeler *et al*., 1999; supplied by EUROSCARF). We integrated the *TRT2* gene at the chromosome XV *HIS3* locus of this strain to give us the starting strain for *synXI* assembly, ysXIa01 (**Table 2**).

#### Assembling megachunks A-L

Prior to transformation into the recipient yeast strain, we enzymatically assembled the constituent megachunks of *synXI* from synthesized DNA *in vitro*, as previously described (Dymond *et al*., 2011; Annaluru *et al*., 2014; Mitchell *et al*., 2017; Richardson *et al*., 2017). The overall construction strategy is outlined in **Figure 1a**. We built up the synthetic chromosomal sequence *in vivo* by systematic replacement of the wild type *chrXI* sequence using the switching auxotrophies progressively for integration (SwAP-In) methodology in this manner. After each megachunk integration, we performed PCRTag analysis to confirm replacement of native sequence with synthetic DNA and performed phenotypic growth spot assays to confirm that the synthetic sequence introduced did not cause growth defects.

We successfully performed iterative megachunk integrations to generate strain ysXIa13, containing megachunks A through L, with no fitness defects observed.

#### Debugging a fitness defect caused by sequence changes around the centromere

Isolation of a megachunk M integrant proved to be challenging. No screened transformant colonies that appeared after 3 days were full megachunk M integrants. After leaving transformation plates at room temperature for a further week and screening smaller transformant colonies that became visible, we identified one colony with fully integrated megachunk M sequence, designated strain ysXIa14. This strain had a severe growth defect (i.e. a bug) on YPD at 30°C. (**Fig S1A**). Replacement of native sequence with the 40.5 kb megachunk M introduces many sequence variants that could affect strain fitness including 9 loxPsym insertions, 3 TAG stop codon recoding events, 4 intron deletions, 1 deletion of a tRNA and associated LTR sequence and 45 recoded sections in CDSs that introduce PCRTags (**Fig 2A**). We assumed that one such variant was responsible for this bug; to identify the bug-associated locus, we reintroduced the five “M” chunks individually into strain ysXIa13 by targeting the corresponding chromosomal loci with CRISPR/Cas9 and providing the synthetic chunk DNA as the repair template (**Fig2A**). Transformants from CRISPR/Cas9 reactions introducing chunk M2 grew visibly more slowly than those from reactions introducing the other chunks, leading us to focus on the M2 sequence for causes of the bug.

**Figure 2:**
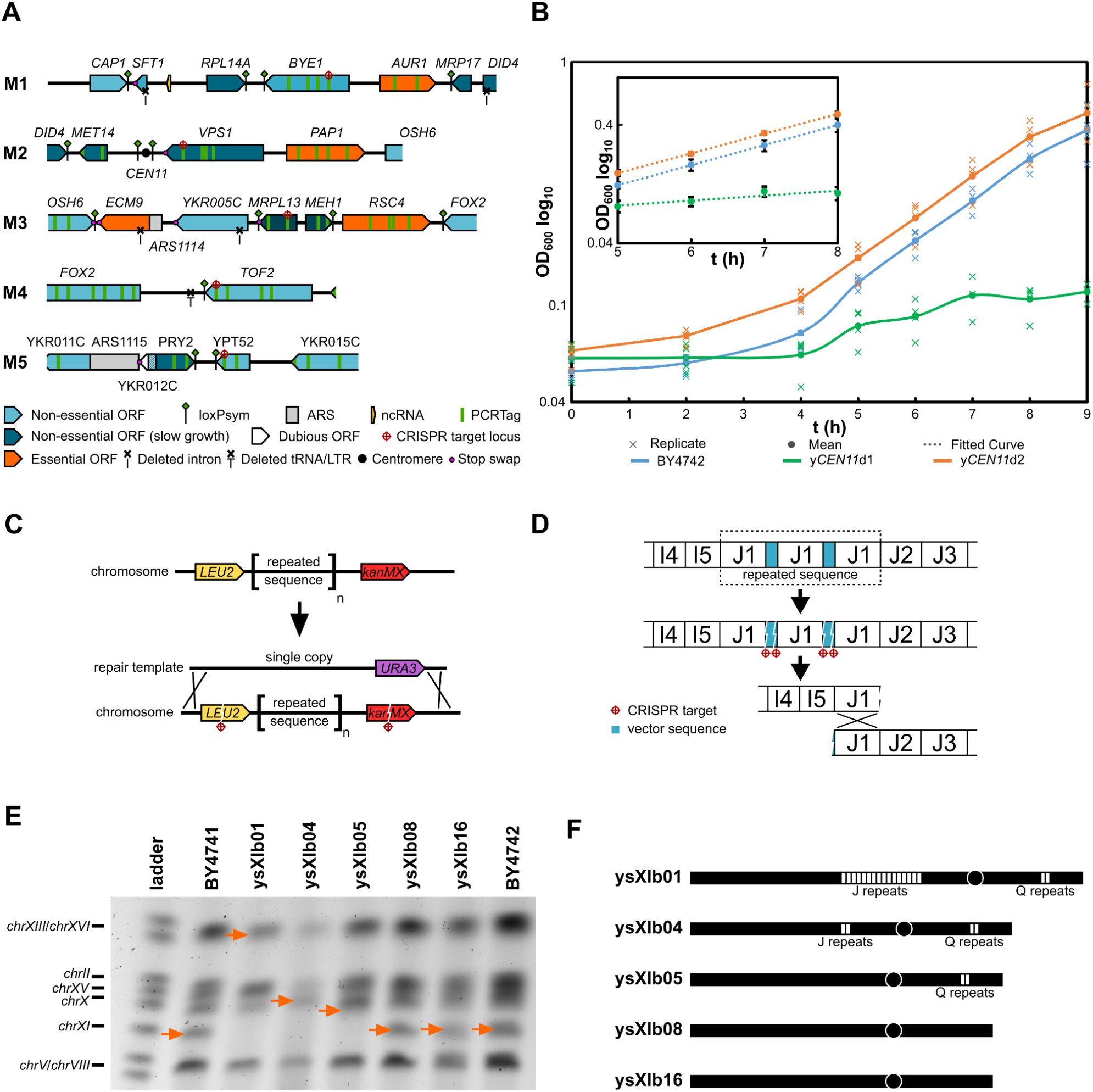
Debugging of centromere and repeated sequence regions in synXI. (A) Overview of the synthetic chromosomal locus corresponding to megachunk M, subdivided into constituent chunks. Changes to the sequence made during synthetic redesign are highlighted with symbols explained beneath the overview. (B) Growth of BY4742 and strains containing centromeric locus variants. Biological replicates are plotted as crosses, n=3, with the mean value plotted as a solid line. Inset is a selection of the same data taken from a period when all cultures were undergoing exponential growth, with mean values plotted, error bars representing standard deviation and a fitted logarithmic curve as a dotted line (BY4742 growth rate [μ] = 0.390 h^-1^, R^2^ = 1; yCEN11d1 μ = 0.096 h^-1^, R^2^ = 0.790; yCEN11d2 μ = 0.384 h^-1^, R^2^ = 1). (C) Overview of the initial strategy to condense repeats in the megachunk J region. (D) Structure of megachunk J repeat sequence as deduced from nanopore sequencing data and the revised strategy to condense these repeats in vivo by CRISPR-mediated recombination. (E) PFGE gel of genomic DNA extracted from BY4741 and various strains generating during the repeat condensation process. Orange arrows show the inferred position of chrXI or synXI. (F) Diagrammatic overview of the repeat sequences in the strains analysed by PFGE in panel E. Each white box represents a predicted copy of repeated sequence. See also Figure S2 and Figure S3.

Close inspection of chunk M2 revealed an unannotated design error in the sequence proximal to the centromere, *CEN11*. The planned design had been to insert a loxPsym site 100 bp either side of the annotated *CEN11* sequence. However, rather than inserting a loxPsym site 100 bp downstream of *CEN11*, 34 bp of native sequence had been deleted and replaced by a loxPsym site now situated 66 bp downstream of the centromere. Although the deleted bases fell outside of the annotated centromeric sequence, we decided that this anomaly in the design warranted further investigation.

To determine the effect of the synthetic *CEN11* region on fitness, we used CRISPR/Cas9 to replace the centromeric region of strain BY4742-*CEN11\** with a 2.1 kb section of chunk M2, spanning the centromere and the surrounding sequence. The resulting strain, y*CEN11*d1, displayed a clear growth defect in YPD compared to BY4742 (**Fig 2B**).

We then redesigned the centromeric region of chunk M2 to restore the inadvertently deleted sequence downstream of *CEN11* and move the loxPsym site into the 3’ UTR of the next gene, VPS1 (**Fig S1B**). Strain y*CEN11*d2, in which the centromeric region of BY4742-*CEN11\** has been replaced with the redesigned centromeric sequence (*CEN11*_3_37) and does not display a growth defect when grown in YPD (**Fig 2B**).

To further investigate whether other permutations of the centromeric locus might affect fitness, we generated 3 further designs. In *CEN11*_3_37b we inserted loxPsym sites on either side of *CEN11*. In *CEN11*_3_37c we replicated the original design error, deleting 34 bp of sequence, but this time at the site upstream of *CEN11*. Finally, in *CEN11*_3_37c we tested whether the specific sequence deleted in the design error was important by inserting the loxPsym site 100 bp downstream of *CEN11* and replacing the preceding 34 bp with a random sequence with the same GC content as the replaced sequence. The topologies of these *CEN11* variants are shown in **Figure S1B**. As the growth defect associated with the synthetic *CEN11* locus was particularly pronounced at 37 °C, we assayed the *CEN11* variant strains for growth at this temperature. None of the strains, except *CEN11*_3_35, showed a noticeable effect on growth (**Fig S1C**). We concluded that the function of the centromere was impaired by insertion of the loxPsym site too close to the right of the centromere. We thus moved forward with chromosome design version *synXI*_3.37, incorporating redesigned *CEN11* region *CEN11*_3_37 in a new version of chunk M2, which could now integrate without conferring a fitness defect in YPD. This new version of megachunk M was successfully integrated, using an approach described in the methods section, yielding strain ysXIa18.

#### Introducing a tRNA array

Whilst the integration of *TRT2* sequence at the *HIS3* locus complemented the deletion of *TRT2* as part of the integration of megachunk B, this solution does not complement the other tRNAs removed during assembly of *synXI* and is not compatible with the downstream consolidation of multiple synthetic chromosomes in one cell. A vector containing an array of the native tRNA genes of *chrXI* under the control of promoter and 3’ untranslated region (3’UTR) elements from *Ashbya gossypii* and *Eremothecium coryli*, pRS413-*chrXI*_tRNA, was generated as part of the construction of a tRNA neochromosome (Schindler, D. *et al*, in preparation). We decided that this plasmid-based construct represented a more favorable method of tRNA gene complementation that would travel with *synXI* during genetic crosses.

To remove *TRT2* from the *HIS3* locus, we mated strain ysXIa18 with strain Y15078 (BY4742 *YKR007W::kanMX4*, EUROSCARF) to generate the diploid ysXIa19. We then used CRISPR/Cas9 mediated homologous recombination to replace the *HIS3::TRT2* locus with *ΔHIS3* locus template sequence PCR-amplified from BY4741. We transformed the resultant strain with tRNA array vector pRS413-*chrXI*_tRNA, sporulated and dissected tetrads. From this we isolated strain ysXIa22, a haploid strain with full chromosomal megachunk A-M sequence, no *TRT2* sequence at the *HIS3* locus and the pRS413-*chrXI*_tRNA vector.

#### *Completing* synXI *assembly by combining two semi-synthetic chromosomes* in vivo

After successful integration of megachunk N to generate ysXIa23, we had difficulty fully integrating megachunk O. The sequence around the junction of chunks O3-O4, corresponding to genes *HFL1* and *MRS*4, consistently failed to integrate. As with the integration issues with megachunk M, we presumed this to be due to inefficient *in vitro* assembly. We isolated a strain, ysXIa24, with incomplete integration of megachunk O with residual wild-type sequence at the presumably problematic junction. We co-transformed this with a CRISPR/Cas9 system targeted to the *YKR051W*_1_WT_R PCRTag sequence and a repair template consisting of *in vitro* ligated and gel-purified O3 and O4 chunks. Using this approach, we successfully isolated integrants without the need for marker swapping and subsequent auxotrophic marker integration. The resulting strain, ysXIa25, had full integration of megachunks A to O (**Fig 1B**).

To increase the rate of synthetic chromosome construction, we iteratively integrated megachunks P, Q and R into a second construction strain, ysXIa26 - a strain we generated by integrating chunk O5 into BY4742 (**Fig 1C**). Integration of the final megachunk, megachunk R, resulted in a *URA3* auxotrophic marker proximal to the universal telomere cap of the right chromosomal arm. We removed this via CRISPR/Cas9 editing. The strain containing *synXI* sequence from chunk O5 through to the right arm telomere, ysXIa30, showed no fitness defects on YPD medium.

Prior to combining the two completed synthetic sections of chromosome XI, we modified the wild type *CEN11* centromere region of ysXIa30 to enable selective loss of this chromosome. Activating transcription from an inducible promoter upstream of a centromere has previously been shown to disrupt centromere function and cause loss of the chromosome during mitotic growth(Hill and Bloom, 1987). To implement this, we integrated a construct containing a *URA3* gene derived from *Kluyveromyces lactis (Bakota et al., 2012*) and the *GAL1* galactose inducible promoter into the *CEN11* region of ysXIa30. In this strain, ysXIa31, the *GAL1* promoter drives transcription through the centromere upon induction with galactose (**Fig S2**). We then mated strains ysXIa25 (*MAT**a***, megachunks A-O, pRS413-*chrXI*_tRNA) with ysXIa31 (*MATα*, megachunks O5-R, *CEN11*::*klURA3*-*PGAL1*) to form a diploid strain, ysXIa32, with all of the *synXI* sequence, albeit on two separate chromosomes.

To combine the two synthetic sections into one complete synthetic chromosome, we designed a CRISPR gRNA that targets the border of chunk O5 and the *LEU2* marker inserted in the 5’ end of O5 in the chromosome inherited from ysXIa25 (with megachunks A-O). We transformed this with *cas9* into the diploid ysXIa32, generating a double strand break (DSB) at the 5’ end of the synthetic section of *synXI*.A-O which could be repaired via host-mediated homologous recombination using the chromosome inherited from ysXIa31 (*synXI*.O5-R) as repair template (**Fig 1D**). This generated the fully synthetic chromosome *synXI*_9.01. We then grew the CRISPR/Cas9 transformant cells overnight in YPGal to induce the *GAL1* promoter upstream of *CEN11* and force loss of the non-synthetic copy of *chrXI*. After plating the culture, we tested 6 colonies by PCRTag analysis to verify that all synthetic - and no wild type - *chrXI* sequence was present. All colonies tested had no detected wild type sequence and contained a fully synthetic chromosome XI. We sporulated the *synXI* diploid strain and dissected tetrads to isolate strain ysXIb01, a haploid strain containing *synXI*_9.01(**Fig 1E**).

Whole genome sequencing and analysis using the Perfect Match Genomic Landscape strategy (Palacios-Flores *et al*., 2018) revealed that we had constructed *synXI*, but with several deviations from the designed sequence. A full list of these deviations is given in **Table S5**. The most notable of these was a marked increase in coverage depth at regions corresponding to chunks J1-J4 and chunks Q1-Q2 (**Fig S3A**). This indicated that these sequences are repeated several times on the chromosome. Indeed, the size of *synXI*_9.01 in ysXIb01 did visualize as around 200kb larger than expected when analysed for size by Pulsed-Field Gel Electrophoresis (PFGE, **Fig S3B**).

### Debugging the fully assembled synthetic chromosome

#### Repeat sequence condensation and transposon sequence removal

To remove the excess repeats in the regions of *synXI*_9.01 corresponding to megachunk J, our initial strategy was to use CRISPR/Cas9 to reintroduce megachunk J DNA, replacing the multi-copy locus with a new single-copy region. To prepare strain ysXIb01 for this operation, we chromosomally re-integrated chunk I5, to introduce *LEU2* into *YKL069W*, and also integrated a modified chunk J4 with a *kanMX4* selection cassette inserted into *YKL053C-A*, giving us strain ysXIb02. Megachunk J was integrated into this strain, along with *cas9* and gRNAs targeting DSBs to the *LEU2* and *kanMX4* markers (**Fig 2C**). Using PFGE, we then analysed the chromosome size of 3 transformants that had lost *LEU2* and *kanMX4* and had gained the *URA3* marker in megachunk J DNA (**Fig S3C**). Colony C had a chromosome band most consistent with significant *synXI* repeat reduction. We designated this strain as ysXIb03.

To confirm the extent of repeat DNA sequence removal at the J locus, we performed nanopore sequencing with ysXIb03 genomic DNA. This showed that some repeat sequence was still present but read lengths were now long enough to capture the entire region, showing just 3 copies of chunk J1 inserted in tandem. These reads also revealed that the J1 repeats were interspersed with plasmid backbone sequence from the J1 chunk vector, a feature likely to have been missed when aligning short reads to a scaffold sequence. We found by restriction analysis of the J1 chunk plasmid that the restriction enzyme *Sfi*I did not cut effectively at the 3’ end of the chunk to release it from the vector. We reason that this led to a situation where partially-digested J1 chunk vectors were concatenating during the *in vitro* ligation, meaning that the megachunk J molecules transformed into yeast for chromosomal integration were containing multiple J1 repeats.

To remove the remaining repeated sequence at the megachunk J locus by CRISPR-directed gap-repair, we transformed ysXIb03 with *cas9* and gRNAs targeting the J1 plasmid sequences flanking the J1 chunk DNA (**Fig 2D**). PCR screening of transformant cells to confirm loss of plasmid-derived sequence led us to isolate strain ysXIb04, in which megachunk J repeats had been condensed down to single copy.

Due to its smaller size, we could take a more straightforward approach to repeat sequence condensation in the megachunk Q region. First, we inserted a *URA3* marker upstream of the repeated Q region in ysXIb04 via re-integration of chunk P5. We then co-transformed into this strain a full copy of megachunk Q and a CRISPR/Cas9 construct targeting the *URA3* marker (**Fig S3D**). Using PCR screening we were able to isolate a transformant, ysIXb08, in which the Q condensed to single copy. We conducted PFGE analysis that supported the reduction of the repeats in ysXIb08 (**Fig 2E-F**).

Nanopore sequencing of strain ysIXb08 confirmed that repeated sequences in regions J and Q had been successfully removed. Interestingly, when analysing the long sequencing reads, we also noticed that there was a sequence discrepancy in the *TRK2* CDS that we had missed by short read sequencing analysis. The CDS showed a partial duplication and the insertion of 2 foreign sequences (**Fig S4**). One of these sequences showed partial identity to the bacterial vector on which the corresponding synthetic DNA chunk was propagated and the other had 99% identity to a gene encoding an *E. coli* transposase DDE domain protein (Genbank accession QFU33765.1). We assume that the O3 chunk vector caused a fitness defect in the *E. coli* host and a transposon insertion into the vector was selected-for during pre-assembly plasmid propagation.

To remove the bacterial transposon-associated insertion sequences, we repeated the propagation of the O3 chunk vector in the *E. coli* host, but now growing the cells slowly at 18°C to reduce burden during growth and thus reduce the chances of stress-induced transposon insertions into the vector. We transformed this chunk DNA into ysXIb08, along with a CRISPR/Cas9 construct targeting the transposase insertion sequence. From the resulting transformants we were able to isolate strain ysXIb09, in which the *TRK2*-transposon locus had been replaced by a single intact copy of *TRK2*.

#### Debugging a respiratory growth defect associated with megachunk Q

Routine spot assays performed after every round of integration during *synXI* construction revealed a persistent growth defect at 37°C on glycerol growth medium (YPG) following megachunk Q integration (**Fig 3A, Fig S5A**). Unlike previous defects during chromosome assembly, this defect was not rescued by the subsequent round of megachunk integration and so not related to marker gene insertion into the locus. As glycerol is a non-fermentable carbon source, this defect is indicative of a problem with mitochondrial function.

**Figure 3:**
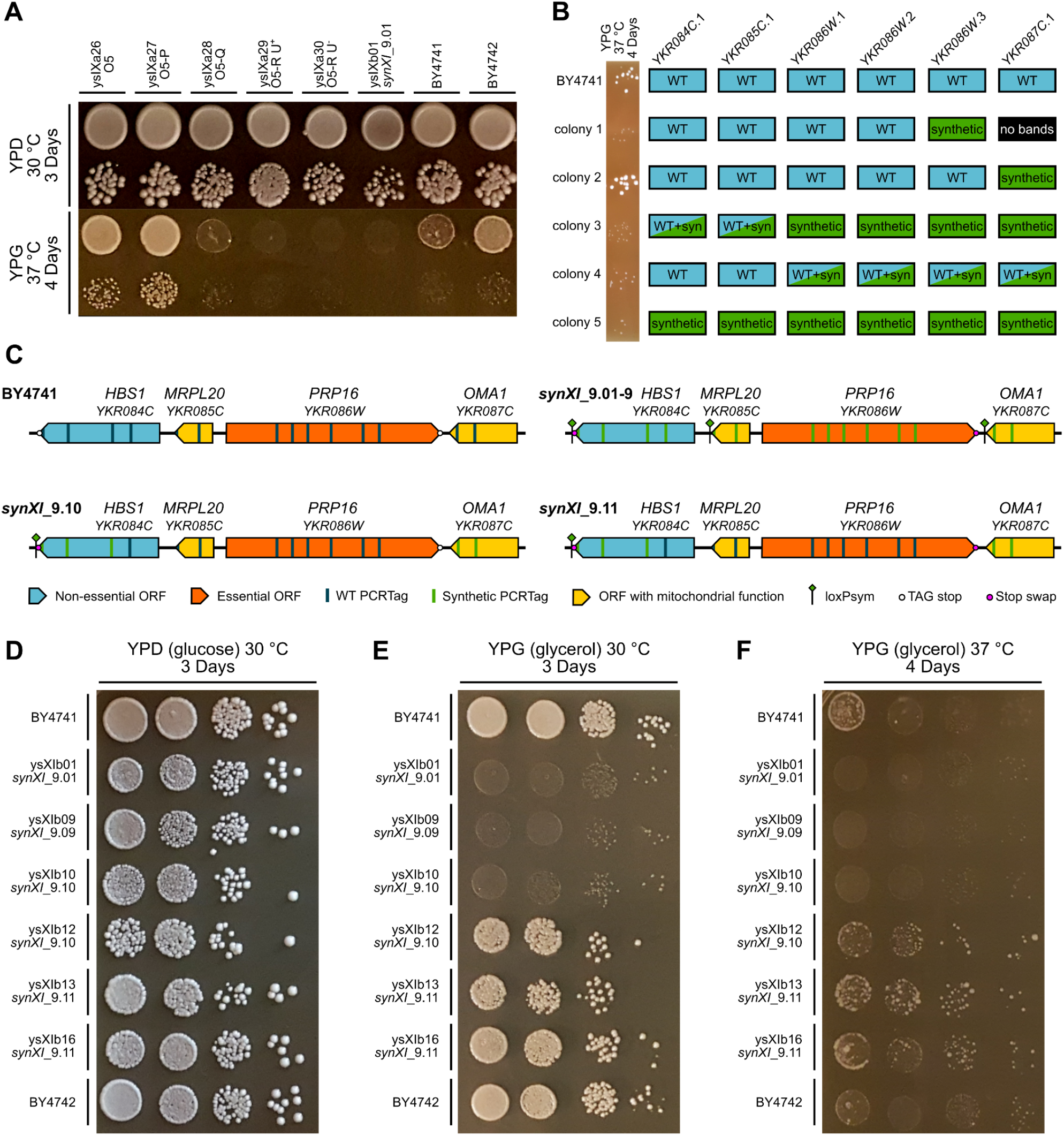
Successful debugging of a respiratory growth defect associated with megachunk Q. (A) Growth spot assays of synXI assembly intermediates following megachunk integration on YPD (glucose) and YPG (glycerol) to test respiratory function. For each strain and condition, the top spot is a ×10^−1^ dilution and the bottom spot is a ×10^−3^ dilution. (B) shows the PCRTag screening results at the YKR084C.1-YKR087C.1 locus for debugging megachunk Q transformant colonies, with corresponding ×10^−3^ dilution YPG 37 °C growth spots to the left, indicating respiratory function. (C) Schematics of the HBS1-OMA1 locus in BY4741 and the in vivo synXI iterations. (D-F) Growth spot assays with strains involved in respiratory growth defect debugging on YPD (glucose) and YPG (glycerol) to test respiratory function. Dilution spots increase in steps of ×10^−1^ from ×10^0^ on the left to ×10^−3^ on the right. See also Figure S5.

Standard debugging approaches revert synthetic DNA regions back to wild type sequence but doing this in ysXIb09 failed to yield any strains with rescued growth on YPG. This indicated that the defects might be in genes encoding mitochondrial proteins and that these defects could cause permanent damage to, or loss of, the mitochondrial DNA. We also found that in contrast to BY4741, ysXIb01 colony growth was not affected by treatment with ethidium bromide and we were unable to obtain any amplification product from PCR screens using mitochondrial genome-specific screening primers using ysXIb01 genomic DNA as template (**Fig S5B-C**). These observations all pointed toward an absence of the mitochondrial genome - meaning that even if the underlying genetic cause of the defect was fixed, full respiratory capacity would remain absent as functional mitochondria couldn’t be restored.

To make debugging possible, we attempted to integrate into strain ysXIa27 (O5-O integrant) using as much synthetic megachunk Q DNA as possible without introducing the defect. We co-transformed megachunk Q DNA with CRISPR/Cas9 constructs targeting DSBs to chromosomal sites in three locations, the chunk P5 *URA3* marker, *MSA2W* and *HBS1*. From this transformation, we were only able to isolate 5 colonies that phenotypically screened as having lost *URA3* and gained *LEU2*. However, PCRTag screening of these colonies revealed all 5 had almost complete integration of megachunk Q, apart from at a locus between PCRTags *YKR084C*.1 and Y*KR087C*.1. For each of these colonies, we compared PCRTag composition to YPG 37°C phenotype (**Fig 3B**). Colony 2 had no associated YPG 37°C defect and had a fully synthetic megachunk Q region, except between PCRTags *YKR084C*.1 and *YKR086W*.3. We confirmed with further PCR screens that the loxPsym site introduced into the 3’UTR of *OMA1* was also absent in this strain.

The synthetic reformatting of the identified region in megachunk Q notably contains loxPsym insertions into the 3’UTRs of two genes encoding mitochondrial proteins, *MRPL20* and *OMA1* (**Fig 3C**). Deletion of *MRPL20* has been previously shown to result in mitochondrial genome loss (Kitakawa *et al*., 1990; Zhang and Singh, 2014) whereas *OMA1* is involved in maintaining respiratory supercomplexes, with null mutants showing deterioration in respiratory function (Bohovych *et al*., 2015). These phenotypes are consistent with the fitness defect observed in the *synXI* strains. We hypothesized that the loxPsym insertions may interfere with 3’UTR encoded targeting of mRNA to the mitochondria (Marc *et al*., 2002).

To fix the respiratory growth defect, we used CRISPR/Cas9 to replace the synthetic region in ysXIb09, that spans from *HBS1* to *OMA1*, with a PCR amplicon of the equivalent region from colony 2 of the debugging process (**Fig 3C**). The resulting strain, ysXIb10, still had the YPG 37 °C growth defect (**Fig 3D-F**) due to the prior mitochondrial damage. To replenish this strain with healthy mitochondria, we backcrossed it with BY4742-*CEN11*\*, then enriched for *synXI*_9.10 by galactose-induced loss of *chrXI*, before sporulating and isolating the *MAT****a*** strain ysXIb12. This strain, with *HBS1-OMA1* locus replacement and mitochondrial replenishment showed reversion to the parental respiratory growth phenotype (**Fig 3D-F**).

When repairing the *HBS1-OMA1* region, along with removing the loxPsym sites associated with *MRPL20* and *OMA1*, we also reintroduced a TAG stop codon into *PRP16*. To conform to the design criteria of the Sc2.0 genome, we then swapped this TAG stop codon to TAA by CRISPR/Cas9-mediated recombination with a template amplified from chunk Q2. This generated chromosome *synXI*_9.11 in strain ysXIb13. We backcrossed this strain a further time with BY4742-*CEN11\** and isolated *HIS3+/URA3-MAT****a*** strain ysXIb16, which displayed good respiratory fitness (**Fig 3D-F**). This strain underwent PCRTag analysis targeted to loci across *synXI* (**Fig S6A-B**) and genome sequencing to confirm the debugged sequence of *synXI*_9.11.

#### Identifying and resolving ploidy issues

To ensure that there were no discrepancies in *synXI*_9.11 copy number in ysXIb16, we confirmed that full genome sequencing of ysXIb16 showed consistent levels of read coverage across all regions of the genome (**Fig 4A**). However, upon mating ysXIb16 with BY4742 we found that dissected spores had low viability frequencies (**Fig 4B**). When we tested ysXIb16 on L-canavanine plates we failed to observe any surviving colonies, indicating that ysXIb16 is not a haploid strain (**Fig 4C**). As sequencing of this strain showed no discrepancies in read coverage or heterogeneity, we concluded that ysXI16b is a homozygous diploid.

**Figure 4:**
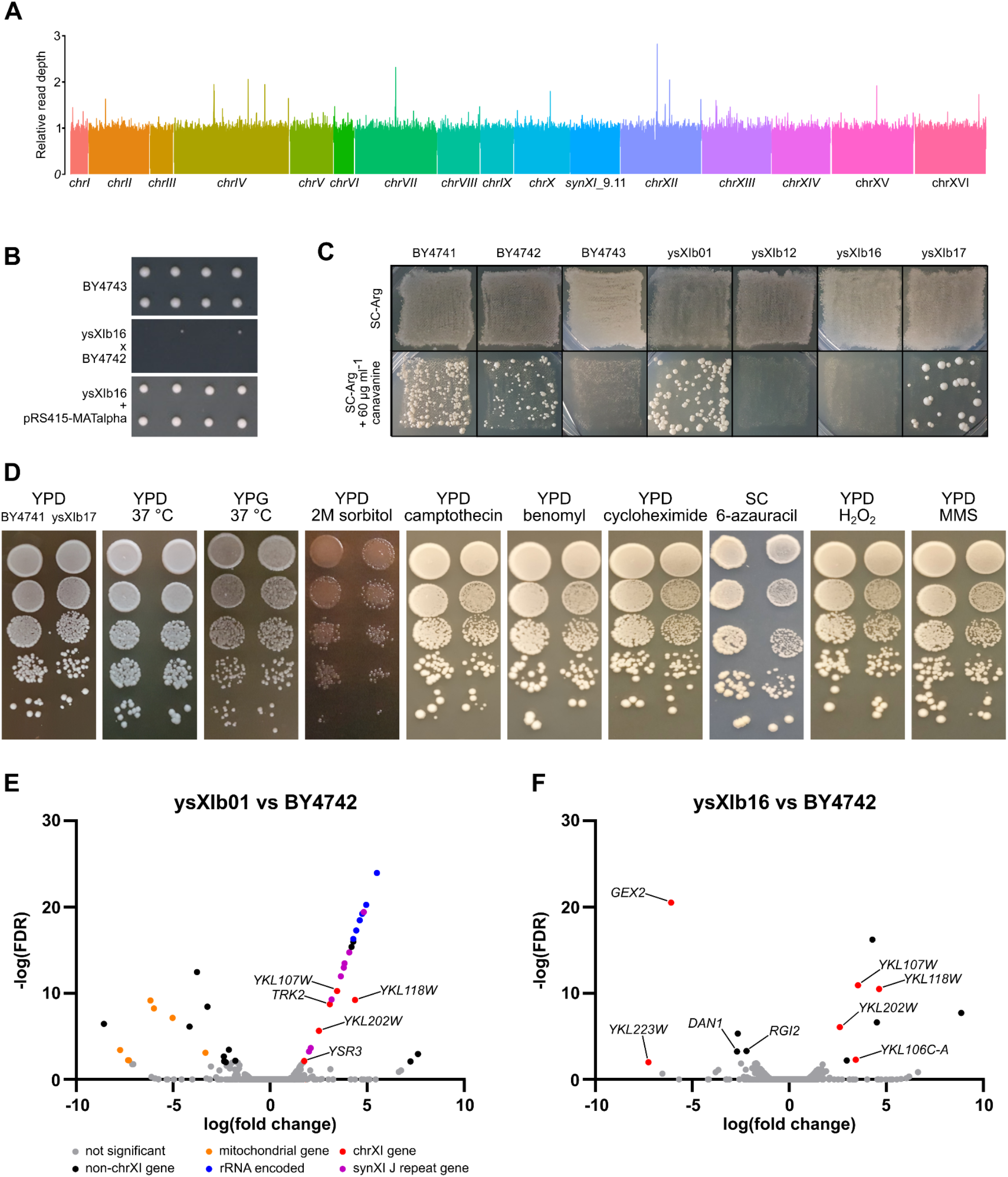
Assessing ploidy, fitness and transcriptional profile of the synXI strain. (A) shows Illumina sequencing read coverage over the whole genome of ysXIb16. (B) Spores from dissected tetrads derived from sporulated strains BY4743, ysXIb16 x BY4742 and ysXIb16 pRS415-MATalpha. Spores from 2 tetrads were dissected for each strain, arrayed horizontally and grown on YPD plates for 2 days at 30 °C. (C) Canavanine ploidy assay patches. Strains were grown on YPD and then replica plated onto SC-Arg with and without canavanine and grown at 30 °C. Growth on canavanine is indicative of haploidy. (D) Growth spot assays of ysXIb17 and a BY4741 parental control in various conditions to assess cellular fitness. Cultures were serially diluted and spotted from top to bottom, with dilutions increasing from ×10^0^ in steps of ×10^−1^. BY4741 was spotted on the left, ysXIb17 was spotted on the right. Sorbitol was added to 2M and camptothecin was added to 1 μgml^-1^. Other additives were added as indicated in the methods section. Unless otherwise indicated, plates were incubated at 30 °C. (E) Volcano plots showing transcript abundance in ysXIb01 compared to BY4742, as determined by RNAseq. (F) Volcano plots of transcript abundance in ysXIb16 compared to BY4742 as determined by RNAseq. For panels E and F, the y axis represents statistical significance in the form of -log_10_ of the false discovery rate, the x axis represents the log_10_ fold change in transcript abundance compared to BY4742 levels. Significant points are those with a false discovery rate <0.01. See also Figure S6 and Figure S7.

To generate a haploid version of ysXIb16 we introduced pYZ412, expressing the *MATα* mating locus, and sporulated the strain. We dissected tetrads and found much-improved spore viability (**Fig 4B**). We performed L-canavanine assays on a colony isolated from tetrad dissection, ysXIb17, which confirmed that this strain is indeed haploid. We performed further canavanine assays that showed the initial *synXI*_9.01 strain, ysXIb01, is haploid but ysXIb12, the strain that underwent backcrossing after editing the *HBS1-OMA1* locus, is not (**Fig 4C**). It is therefore likely that the homozygous diploidy was introduced during this backcrossing process.

### Assessing the fitness and transcriptional profile of *synXI* strain

We performed growth spot assays with ysXIb17 on a wide range of media types and conditions designed to test the robustness of various cellular processes (**Fig 4D, Fig S7**). Under all conditions tested ysXIb17 performed well and showed no notable defects compared to the parental strain, including in YPG media.

We next used RNAseq analysis to compare the transcriptional profiles of the parental strain to the initial *synXI* assembly strain ysXIb01 and to ysXIb16, which contains the debugged *synXI*_9.11. We selected BY4742 as our parental control as its auxotrophic profile most closely matched the *synXI* strains. For the purposes of our analysis, we eliminated mating-type specific genes to compensate for mating type differences between the parental and *synXI* strains. This included transcripts of the dubious ORF *YKL177W* as it almost entirely overlaps *STE3*, which encodes the receptor for a factor pheromone in *MATα* cells. The CDS of *FLO10* was entirely recoded by an early version of REPEATSMASHER due to the presence of highly repetitive sequence (Richardson *et al*., 2017). As a result, the *synXI FLO10* has <70% identity to the wild type version. This lack of sequence similarity and the repetitive nature of the mRNA in BY4742, coupled with the extremely low expression levels typically observed in S288C-derived strains, led us to also omit *FLO10* from our analysis.

The genes showing significantly different expression (FDR<0.01) compared to BY4742, from the initial strain before debugging (ysXIb01) largely fall in a few categories (**Fig 4E**). 8 genes with higher expression are located in the repeat sequences corresponding to megachunk J. Presumably higher transcript levels of these genes are a result of their expanded copy number. 6 genes with lower expression are encoded on the mitochondrial genome and their differential expression is likely related to the respiratory growth defect we see in the cells. Another interesting group with higher transcription levels maps to ORFs encoded within the 35s rRNA. The function of these ORFs, *YLR154W*-A/B/C/E/F and *YLR154C*-G, is not clear but their increased transcription may indicate an increase in the copy number of the DNA encoding the 35S rRNA. Of the other differentially expressed genes, 5 are located on *synXI*. These include *TRK2*, the site of the bacterial transposon insertion.

When comparing the transcriptional profile of the final strain, ysXIb16 to BY4742 (**Fig 4F**), we observe that the debugging process has reverted the transcription of the megachunk J repeat genes, the mitochondrial genes, the transcripts embedded in the 35s rRNA and *TRK2* to showing no significant differences to the parental strain. There are 6 significantly differentially expressed genes that are located on *synXI. GEX2* and *YKL223W* neighbor the telomeres, so we assume that their downregulation in ysXIb16 is due to a proximal telomeric repression effect. Increased transcription in 3 other genes, *YKL107W, YKL118*W and *YKL202W*, is also seen in ysXIb01. The remaining differentially expressed *synXI* gene YKL106C-A is found 77 bp downstream of the *YKL107W* CDS and we assume its increased transcript level in the analysis is due to the two genes having overlapping transcripts (Xu *et al*., 2009). *YKL118W* and *YKL202W* are both small dubious ORFS overlapping sequences changed in the synthetic redesign process. In the case of *YKL118W*, this is a Ty1 LTR and *YKL202W* overlaps repetitive sequence 3’ of *MNN4*. For both of these ORFs, we do not expect the altered levels of transcripts identified in the RNAseq analysis to have notable biological effects. Of the *synXI* genes, only the increase in *YKL107W* transcription is likely to have biological relevance. This gene encodes an aldehyde reductase involved in the detoxification of toxic aldehydes (Wang *et al*., 2019). An explanation for this increase in transcription is not immediately clear to us, although recoding of the CDS to incorporate PCRTags and a BstEII restriction site may be the underlying cause.

There are a further 7 genes showing significantly different expression between ysXIb16 and BY4742 that are not located on *synXI*. These consist of 5 ORFs that are of dubious or unknown function, and 2 further genes with lower transcription in ysXIb16: *RGI2* and *DAN1*. Neither *RGI2* or *DAN1* were differentially expressed in ysXIb01. *DAN1* and *RGI2* have previously been shown to be repressed by aerobic growth (Sertil *et al*., 1997) and high glucose conditions (Domitrovic *et al*., 2010) respectively. As the transcriptional changes are modest and limited to these 2 genes, slight differences in oxygen and glucose availability to the cells whilst culturing the strains may explain these differences.

### The SC2.0 format *GAP1* locus is an effective tool for studying extrachromosomal circular DNA behavior

Our characterization of strains in which *synX*I_9.11 completely replaces *chrXI* revealed very close transcriptional and phenotypic similarity to the parental strains. However, it is possible that the design principles we implemented have more subtle or situation-specific effects that we have not observed. One behaviour that could be expected to be substantially affected is the formation of certain species of extrachromosomal circular DNA (eccDNA). The canonical example of a functional eccDNA in yeast is the *GAP1* eccDNA, thought to be formed via a recombination event between LTR sequences flanking *GAP1* and ARS1116 in *chrXI* (Gresham *et al*., 2010; Møller *et al*., 2013; Møller *et al*., 2015) (**Fig 5A**). The circularisation of *GAP1* and ARS1116 into an eccDNA allows cells to vary the copy number of *GAP1* within a population. *GAP1* encodes a general amino acid permease (Jauniaux and Grenson, 1990) and *GAP1* copy number expansion through eccDNA formation is thought to be enriched-for in nitrogen limited conditions by improving a cell’s ability to import amino acids under nitrogen starvation (Gresham *et al*., 2010; *Møller et al*., *2015*). In the *synXI* synthetic reformatting process the LTR (d) regions involved in *GAP1* locus circularisation, *YKRCd11* and *YKRCd12*, have been removed and replaced with loxPsym sites (**Fig 5B**). The synthetic *GAP1* locus is therefore an ideal testbed to study the potential effects of synthetic chromosome redesign,including LTR removal, on the formation of eccDNA through recombination-based mechanisms.

**Figure 5:**
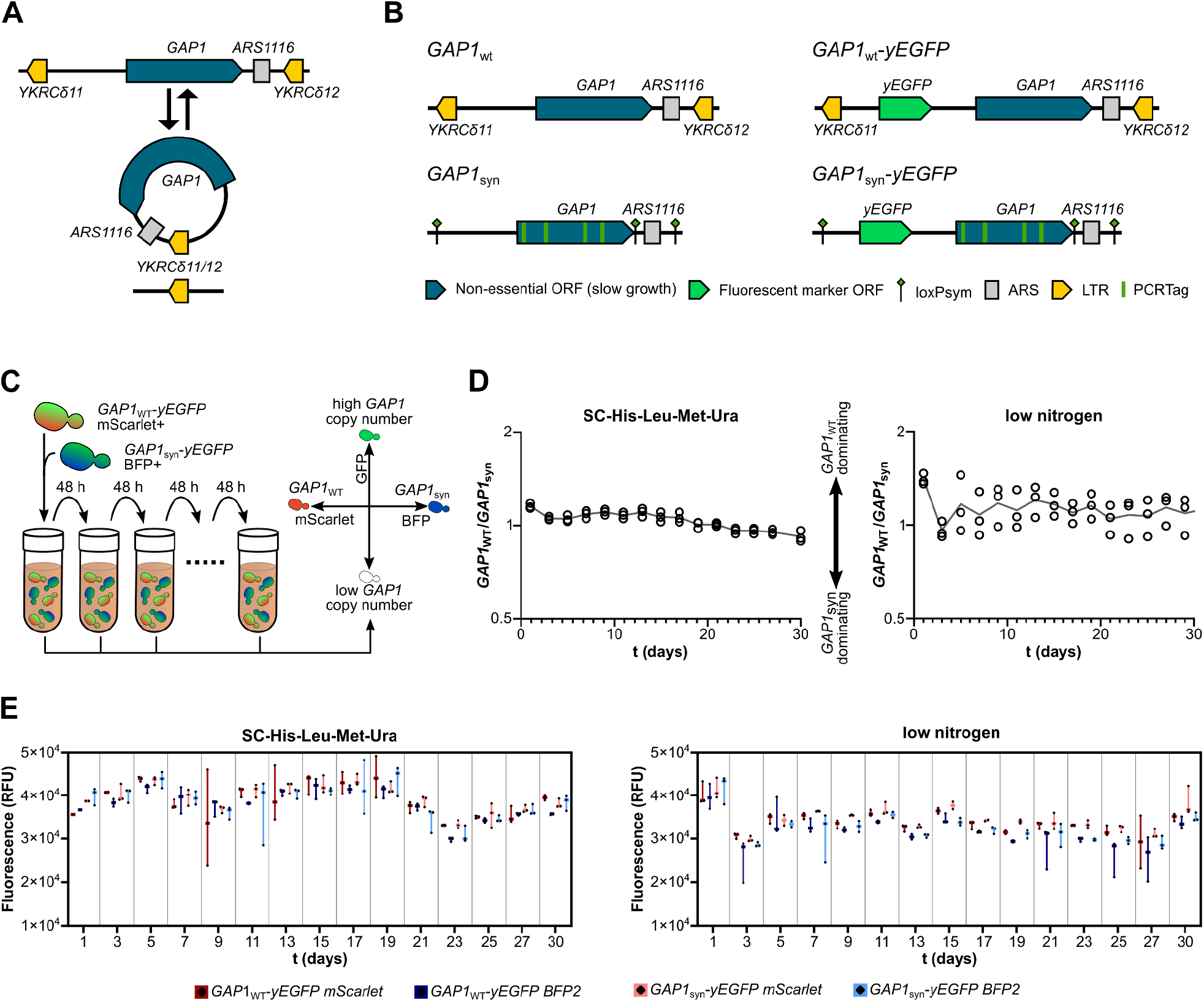
The non-eccDNA forming synthetic GAP1 locus is not detrimental to growth under nitrogen limitation. (A) Overview of eccDNA formation at the GAP1 locus. (B) Structure of GAP1 loci built and integrated into BY4741 to study eccDNA dynamics. (C) Overview of the competition assays and the determination of strain background and eccDNA copy number through fluorescence. Competition assays are sub-cultured every 48 hours with a sample analysed by flow cytometry to assess the ratio of cell types (red fluorescence vs blue fluorescence) and GAP1 copy number (green fluorescence). (D) The ratio of BY4741-GAP1_WT_-yEGFP-mScarlet to GAP1_syn_-yEGFP-BFP cells under competitive growth sampled over the duration of the competition assays in synthetic and low-nitrogen media. Biological replicates (n=3) are plotted as circles and mean values are plotted as a line. Y axis has a log_2_ scale. Data for competition assays between BY4741-GAP1_WT_-yEGFP-BFP and GAP1_syn_-yEGFP-mScarlet cells is shown in Figure S9. (E) Geometric mean GFP fluorescence values of each cell type in each of the competition assays as determined by flow cytometry. Each biological replicate (n=3) is plotted as a diamond with the range represented by a bar. See also Figure S8 and Figure S9.

First, we set out to investigate synthetic redesign effects on *GAP1* eccDNA formation within populations over time. To do this, we generated a strain of BY4741 in which the native *GAP1* locus (*GAP1*_*WT*_) was replaced with the synthetic redesigned *GAP1* locus (*GAP1*_*syn*_) found in *synXI*_9.11. The *GAP1*_*syn*_ locus had no discernible effect on strain growth in YPD, SC or low nitrogen growth media (**Fig S8**). We then inserted a *yEGFP* fluorescent marker gene into the region upstream of *GAP1*, in both the BY4741 and BY4741-*GAP1*_*syn*_ strains, enabling use of GFP fluorescence as a proxy for *GAP1* locus copy number (Lauer *et al*., 2018; **Fig 5B**). Again, no effects were seen on strain growth due to this gene insertion (**Fig S8**). Next, we generated versions of BY4741-*GAP1*_*WT*_-*yEGFP* and BY4741-*GAP1*_*syn*_-*yEGFP* tagged to have externally inducible red and blue fluorescence, respectively. This required integration of *mScarlet* or *BFP2* fluorescent marker genes into chromosome III at the *LEU2* locus. Finally, we transformed all strains with the plasmid pHLUM (Mülleder *et al*., 2012) to give all strains full prototrophy.

Using these strains, we set up long term competition assays in both low nitrogen media and synthetic media, initially inoculating with a precise 1:1 ratio of BY4741-*GAP1*_WT_-*yEGFP*-*mScarlet* and BY4741-*GAP1*_syn_-*yEGFP*-*BFP* or with a 1:1 ratio of BY4741-*GAP1*_WT_-*yEGFP*-*BFP* and BY4741-*GAP1*_syn_-*yEGFP*-*mScarlet* (**Fig 5C**). The competition assays ran for 30 days, with cells sub-cultured every 2 days and a sample of these analysed by flow cytometry for BFP, mScarlet and GFP fluorescence. We found that cells with *GAP1*_*WT*_ were unable to outcompete cells with *GAP1*_*syn*_, even in low nitrogen conditions (**Fig 5D, Fig S9**). We also saw that *GAP1* copy number, as inferred from GFP fluorescence, did not increase in any of the strains in either media condition (**Fig 5E**). We conclude from this that in batch culture, *GAP1* eccDNA formation in BY4741-derived yeast is either not selectively enriched in low-nitrogen conditions or occurs at such a low rate that it is not detected by a 30 day competition assay.

Isolation and study of cells with a specific rare eccDNA event can be extremely challenging. Not only are cells with the circularization event potentially difficult to isolate, but the inherent instability of eccDNA and its asymmetric inheritance pattern mean that presence of the eccDNA over the course of an experiment can be difficult to maintain (Arrey *et al*., 2022). The *GAP1*_*syn*_ locus offers a unique way of bypassing these issues as the elements responsible for the circularisation mechanism have been replaced by loxPsym sites, which can be targeted for recombination by Cre recombinase. We introduced the SCRaMbLE plasmid p*SCW11*-*cre*-EBD to BY4741-*GAP1*_syn_-*yEGFP*, induced SCRaMbLE with β-estradiol and cured the strains of the plasmid. As Cre was no longer present in the cells and loxPsym sites are not large enough to be targeted by the native homologous recombination machinery, the SCRaMbLE recombination events are not reversible in the way that eccDNA formation through LTRs is. Using this process, we were able to isolate two strains, *GAP1*_SCRaMbLE-9_ and *GAP1*_SCRaMbLE-19_, which had expanded the chromosomal *GAP1* locus, and a third strain, *GAP1*_*SPecc*_, containing a *GAP1* <u>S</u>CRaMbLE <u>P</u>roduced <u>e</u>xtra<u>c</u>hromosomal <u>c</u>ircle (SPecc, **Fig 6A**).

**Figure 6:**
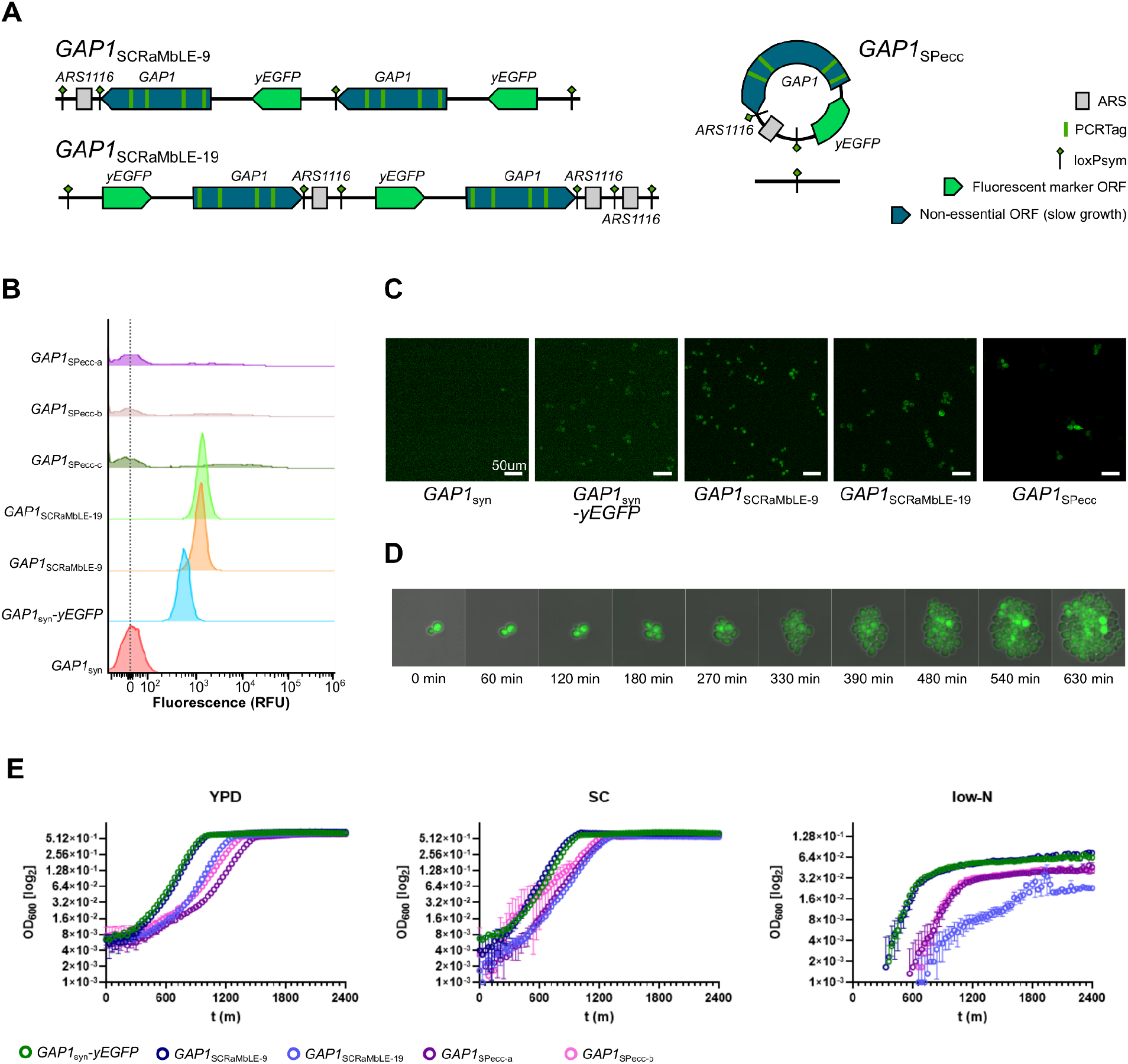
Cre-mediated recombination can circularise an Sc2.0-formatted locus in vivo to form eccDNA. (A) Structure of the GAP1 loci generated through SCRaMbLE. (B) Population GFP fluorescence, as determined by flow cytometry, of strains with GAP1 locus arrangements shown in panel A. The 3 GAP1_SPecc_ samples (a-c) are derived from 3 different GAP1_SPecc_ colonies. Fluorescence values are normalised to the geometric mean fluorescence value of strain GAP1_syn_ (denoted by dashed line). (C) GFP fluorescence microscopy images of GAP1_syn_ cells with various GAP1 locus arrangements. Cell types are given below the images. Images were taken at 20x magnification. (D) Time course GFP fluorescence microscopy images of GAP1_SPecc_ cells taken at 20x magnification. (E) Growth of GAP1 derivative strains in 96 well plates under rich (YPD), defined (SC) and nitrogen limited (low-N) media conditions at 30 °C. Mean OD_600_ values from 3 biological replicates are plotted as circles, error bars represent standard deviation.

We used flow cytometry to determine that the *yEGFP* copy number was increased in both strains with an expanded chromosomal *GAP1* locus (**Fig 6B**). In contrast, the fluorescence profile of GAP1_SPecc_ cultures showed that many cells lost *yEGFP* expression, and thus likely lost the GAP1 SPecc. However, many highly fluorescent cells were still present in the *GAP1*_*SPecc*_ population and showed a spread of fluorescence with a geometric mean around 13 times higher than in the single-copy *yEGFP* parental strain. This is fully consistent with a highly variable gene copy number that would be expected from an eccDNA (**Fig 6B**). We analysed these strains by fluorescence microscopy and the images supported the flow cytometry findings, particularly showing the variable fluorescence seen in *GAP1*_*SPecc*_ cells (**Fig 6C**). Time lapse microscopy showing a single budding *GAP1*_*SPecc*_ cell growing into a population of cells illustrates the uneven inheritance and variable copy number of the *GAP1*_*SPecc*_ within the population (**Fig 6D**). This behavior is consistent with how eccDNA is expected to behave within a population following its formation (Arrey *et al*., 2022).

We used our SCRaMbLE-derived strains to determine the effects of *GAP1* copy number on growth in rich, defined and low nitrogen media (**Fig 6E**). In all conditions, we saw no noticeable difference in growth between the *GAP1*_*SCRaMbLE-9*_ strain with 2 chromosomal copies of *GAP1* and the *GAP1*_*syn*_ strain with a single *GAP1* copy. The *GAP1*_*SCRaMbLE-19*_ strain, which also has 2 *GAP1* copies but also has 2 additional copies of ARS1116, had a clear growth defect in low nitrogen medium. We found that 2 independent isolates of the *GAP1*_*SPecc*_ strain each showed moderate growth defects in rich and defined media. Surprisingly, in low nitrogen medium these cultures with *GAP1*_*SPecc*_ also showed a slight growth defect, delayed entry into logarithmic growth and a lower final density. We have shown that SCRaMbLE-based methods can be used to generate strains with *GAP1* copy number expansion that is either stable, in the case of the chromosomally expanded *GAP1* locus strains, or follows an eccDNA inheritance pattern, in the case of the strains with *GAP1* SPeccs. In either case, there is no positive effect on growth in nitrogen-limited batch culture after *GAP1* copy number expansion.

## Discussion

We have assembled and debugged synthetic chromosome XI of the Sc2.0 synthetic yeast genome. Strain ysXIb17 has full replacement of its native *chrXI* with *synXI*_9.11 and we have found its fitness and transcriptional profiles to be very similar to those of the parental strain.

Complications and difficulties in the assembly process led us to develop a range of effective CRISPR/Cas9 approaches to debugging and editing the designed synthetic DNA *in vivo*. Targeted integration of synthetic DNA chunks, followed by phenotypic screening, allowed us to effectively narrow down the potential sequence causing fitness defects. By focusing on narrower regions of synthetic DNA sequence, we were able to identify underlying causes and precisely edit the problematic sequences to restore fitness. Through the process of debugging by CRISPR/Cas9-mediated integration of megachunk Q, we also had a strong indication that the efficiency of megachunk integration and subsequent colony screening could be markedly improved. Co-transformation of CRISPR/Cas9 constructs, targeting loci across the chromosomal region being replaced, reduced the number of transformant colonies whilst enriching for a high proportion of successfully integrated sequence. Combined with our CRISPR/Cas9 approach to combine chromosome sections *in vivo* via targeted mitotic crossover, we believe that our refined methodologies will allow much faster and more efficient parallelised assembly of future synthetic chromosomes and genomes.

Over the course of assembling and debugging *synXI*, we uncovered cases of redesigned sequences in non-coding DNA leading to phenotypic defects in the host cell. We identified the insertion of loxPsym sites to be a common underlying cause of these fitness defects. Insertion of a loxPsym site 66 bp downstream of the annotated centromere *CEN11* caused a pronounced slowing of growth. Although various permutations made to the sequence around *CEN11* did not produce a similar effect, sequence surrounding annotated centromeres should be edited with caution in future projects. We also found that loxPsym insertions into the 3’ UTRs of *MRPL20* and *OMA1*, both encoding mitochondrial proteins, led to strains defective in mitochondrial function. Additionally, we were unable to isolate a strain with a successful integration of a loxPsym site into *MRS4*, another gene encoding mitochondrial function (Kreike *et al*., 1986). Previous studies have shown that in genes encoding proteins with mitochondrial function, the 3’UTR is important in localizing mRNA to the mitochondrial outer membrane (Marc *et al*., 2002). This occurs through mechanisms that are either dependent or independent of binding to the protein Puf3p via a consensus binding domain within the 3’ UTR (Saint-Georges *et al*., 2008). Previous studies have identified Puf3p binding consensus sequences in the 3’ UTRs of both *MRPL20* and *MRS4* (Foat *et al*., 2005; Gerber *et al*., 2004). We speculate that the designed insertion of loxPsym sites into a small subset of these mitochondrial genes interferes with the correct localization of mRNA to the mitochondria, resulting in defects in respiratory growth. Strains in which these loxPsym sites were removed reverted to a healthy respiratory growth phenotype following mitochondrial DNA restoration. In future synthetic construct design, we would recommend caution in altering the 3’UTR sequences of genes with mitochondrially targeted mRNA or any other regulatory motifs in the region.

As well as fitness defects, another negative feature of the *synXI* construction process was the introduction of structural variations in the form of large DNA sequence repeats, as has been seen in other synthetic yeast chromosome studies (Shen *et al*., 2017; Wu *et al*., 2017; Xie *et al*., 2017). Some of these repeats are artefacts of the construction process. In the case of megachunk J an inefficiently cut *Sfi*I site in the J1 chunk plasmid appears to be directly associated with the generation of the repeats. It is also notable that the repeats in the megachunk Q region contain the majority of the *MRPL20* CDS. As the respiratory growth defect was fixed after we reverted *MRPL20* to its native sequence, it’s possible that the repeats in megachunk Q subtly alleviated the effects of a defective synthetic *MRPL20* and were thus selected for.

Short read sequencing was able to determine that repeats were present, via increased read coverage depth over repeated sequence, but long read sequencing was much better at capturing the structure of repeated sequences and helped us identify the best strategy for their condensation. As long read sequencing data allowed us to construct contigs de novo, without a predicted sequence scaffold, it was also better at identifying unexpected insertions like the bacterial transposase sequence discovered in *TRK2*. We conclude that the use of long read sequencing to help determine the structure of chromosome-scale synthetic constructs is highly beneficial. Where extensive repeats occur, the CRISPR/Cas9 repeat condensation methods we developed here are effective at restructuring a chromosome to remove higher-order deviations from the designed sequence.

The removal of native repeat sequences in the Sc2.0 chromosome redesign process is intended to improve stability and reduce the incidence of genomic rearrangements. However, in some cases dynamic events involving interaction between repeated elements have biological function, such as in some eccDNA formation events. *GAP1* eccDNA formation to adapt to nitrogen-limited conditions is often cited as a classic example of eccDNA function in the literature (Gresham *et al*., 2010; Møller *et a*l., 2013, Møller *et al*., 2015; Arrey *et al*., 2022;). Its location on *chrXI* gave us a good test case to investigate the unintended effects on eccDNA function that could be caused by the synthetic chromosome redesign. The native LTR repeats that were previously shown to recombine to form the *GAP1* eccDNA in wild type strains have been replaced by loxPsym sites in *synXI*, effectively removing the proposed natural circularisation mechanism. We were surprised to find that in our study, we were not able to demonstrate any competitive advantage of cells with a wild-type *GAP1* locus over those with the synthetic locus in prolonged nitrogen-limited batch culture conditions. The formation of *GAP1* eccDNA is thought to be a rare event selectively enriched in conditions in which the eccDNA confers an advantage such as chemostat culture (Gresham *et al*., 2010; Møller *et al*., 2013). But we did not observe any evidence that *GAP1* eccDNA was enriched in our passage experiments, as expression of a fluorescent marker integrated into the *GAP1* locus remained stable throughout.

One explanation is that the eccDNA formation event did not occur during our assay. An alternative explanation is that, in these conditions, *GAP1* eccDNA did not confer any advantage to cells. Whilst *GAP1* eccDNA has been shown to be enriched for in cells showing adaptation to nitrogen limitation in chemostat conditions (Gresham and Hong, 2015), to our knowledge, no previous studies have specifically induced the formation of *GAP1* eccDNA and thus decoupled the direct effects of *GAP1* copy number from other potential cellular adaptations to low-nitrogen media.

The synthetic reformatting of the *GAP1* locus gave us a unique opportunity to directly study *GAP1* eccDNA by using a SCRaMbLE-analogous process to induce irreversible *GAP1* eccDNA formation in cells with a synthetic *GAP1* locus. We were able to show that these SPeccs were inherited asymmetrically down cell lineages and that they resulted in widely heterogenous expression levels within a population. This behaviour follows our expectations of an eccDNA species (Arrey *et al*., 2022), such as that of the native *GAP1*.

Interestingly we found that increased *GAP1* copy number, either through chromosomal tandem repeats or through the formation of SPeccs, did not correlate with improved growth in nitrogen limited conditions. This would suggest that *GAP1* copy number expansion through eccDNA formation alone is not sufficient, in batch culture, to confer an improved growth phenotype in low-nitrogen conditions for BY4741-derived yeast.

The generation of SPeccs allowed us to directly study the effects of the yeast *GAP1* eccDNA on the cell. Whilst the synthetic *GAP1* locus was designed and generated as part of the *synXI* construction process, synthetic loci allowing generation of SPeccs analogous to other eccDNAs would not require full synthetic chromosomes to be exploited. Given the emerging importance of eccDNAs in the fields of evolution, immunology (Wang *et al*., 2021), cancer (Ling *et a*l., 2021; Noer *et al*., 2022) and aging (Hull *et al*., 2019), we believe that the SPecc methodology will be a valuable tool for a wide range of future studies, both in yeast and in other organisms. SPeccs may also be promising new tools for fields such as synthetic biology and biotechnology. The ability to confer a population with widely heterogenous expression of a foreign gene or pathway of interest could be a novel way to determine optimal gene expression levels or to generate strains with self-sacrificing individuals, for example where a minority sequester toxic metals or metabolites to the benefit of the wider population.

The Sc2.0 synthetic chromosome XI has been assembled and will make up part of the complete synthetic yeast genome. Many of the lessons learnt in this project, as well as our approaches to chromosome design, assembly, debugging and structural manipulation of the genome will contribute to future synthetic genome projects, both in yeast and in an expanded range of organisms. Additionally, our study and manipulation of a synthetic eccDNA locus has delivered a new tool enabling us to generate eccDNAs and specifically examine their functions *in vivo*. This will open up new methodologies for synthetic biology and biotechnology as well as providing new tools for those studying the roles played by these DNA species in many aspects of biology.

## Methods

### Strains, media and growth conditions

The *Saccharomyces cerevisiae* strains used and generated in this study are listed in **Table S1**. Unless otherwise stated, liquid yeast cultures were grown shaking at 30 °C in YPD medium (10 g l^-1^ yeast extract, 20 g l^-1^ peptone, 20 g l^-1^ glucose). YPGal medium (10 g l^-1^ yeast extract, 20 g l^-1^ peptone, 20 g l^-1^ galactose) was used to induce galactose-inducible promoters. YPG medium (10 g l^-1^ yeast extract, 20 g l^-1^ peptone, 20 g l^-1^ glycerol) was used to assess respiratory growth. Synthetic complete medium (SC; 6.7 g l^-1^ yeast nitrogen base, 1.4 g l^-1^ yeast synthetic dropout medium supplemented with appropriate amino acids absent, 20 g l^-1^ glucose) was used for auxotrophic selection, or with all amino acids supplemented as a defined complete medium. Low-nitrogen medium (MG; 1.6 g l^-1^ yeast nitrogen base without ammonium sulphate, 20 g l^-1^ glucose, 0.35mM L-glutamine) was used to provide nitrogen limited conditions. For growth on plates, media were supplemented with 20 g l^-1^ agar. Where required for *kanMX4* selection, media were supplemented with G-418 disulfate solution (Formedium) to 250 μg ml^-1^. For testing strain fitness under various perturbations, media were supplemented with sorbitol (osmotic stress; 1M, 1.5M or 2M), camptothecin (topoisomerase inhibitor; 0.1, 0.5 or 1 μg ml^-1^), benomyl (microtubule inhibitor; 15 μg ml^-1^), 6-azauracil (transcription elongation inhibitor; 100 μg ml^-1^), or methyl methanesulfonate (MMS; DNA alkylating agent; 0.05%). When testing cells with cycloheximide (protein synthesis inhibitor; 10 μg ml^-1^) or hydrogen peroxide (H_2_O_2_; oxidative stress; μg ml^-1^) liquid YPD cultures were supplemented with the additive and incubated for 2 hours prior to being washing with water, diluted and plated.

*Escherichia coli* DH10B (Grant *et al*., 1990) (Thermo Scientific) was used for vector cloning and propagation. Luria Bertani medium was used for bacterial growth with ampicillin (100 μg ml^-1^), kanamycin (50 μg ml^-1^) or spectinomycin (50 μg ml^-1^) added for selection as required.

### Strain generation

ysXIa01 assembly recipient strain: We linearised the pRS403::*TRT2* plasmid at the *HIS3* locus by digestion with NdeI and transformed the fragment into Y07039. The successful integrant was strain ysXIa01.

BY4742-*CEN11\** strain with galactose inducible *chrXI* loss: The 2985 bp EcoRI/HindIII restriction fragment of p*CEN11\** was integrated into the *CEN11* locus of BY4742, replacing the native sequence and giving strain BY4742-*CEN11\**. To generate the strains used to test *CEN11* variant configurations, *CEN11* variant regions were PCR amplified from plasmids p*CEN11*_3_37_M2, p*CEN11*_3_37_M2b, pCEN11_3_37_M2c and pCEN11_3_37_M2f using primers BB877/BB878 and integrated into the BY4742-*CEN11\** locus to replace the *CEN11\** sequence using CRISPR/Cas9.

*GAP1*_syn_ Synthetic *GAP1* locus strain: Plasmids pSXI_3_34_O1 and pSXI_3_34_O2 were digested with SfiI and the gel purified chunk DNA sections were ligated together. This ligated DNA was used as template for PCR amplification of the synthetic *GAP1* locus with primers BB582/BB585. The 4668 bp product was integrated into BY4741, replacing the wild type *GAP1* locus, using CRISPR/Cas9.

*GAP1* GFP reporter strains: Strains *GAP1*_WT_-*yEGFP* and *GAP1*_syn_-*yEGFP* were generated by insertion of the *PPFY1*-*yEGFP*-*TCYC1* cassette from pSV-PFY1p (Blount *et al*., 2012) 1118 bp upstream of the *GAP1* CDS of strains BY4741 and *GAP1*_syn_ respectively. This was done using CRISPR/Cas9 with a repair template encoding the insertion sequence assembled from a PCR fragment encoding the *yEGFP* expression cassette amplified from pSV-PFY1p with primers XL494/XL495 and PCR fragments encoding homology arms amplified from BY4741 genomic DNA using primers XL492/XL493 and XL496/XL497. The 3 PCR products were pooled and co-transformed with the CRISPR/Cas9 DNA into the recipient strain, with the repair template being formed through in vivo homologous recombination combining the 3 overlapping fragments.

Competition assay strains with inducible mScarlet reporters: Strains *GAP1*_WT_-*yEGFP*-*mScarlet* and *GAP1*_syn_-*yEGFP*-*mScarlet* were generated by integrating plasmid pXL007, containing a tetracycline-inducible mScarlet expression cassette, into the *LEU2* locus of strains *GAP1*_WT_-*yEGFP* and *GAP1*_syn_-*yEGFP* respectively.

Competition assay strains with inducible BFP reporters: Strains *GAP1*_WT_-*yEGFP*-*BFP* and *GAP1*_syn_-*yEGFP*-*BFP* were generated by integrating plasmid pXL008, containing a tetracycline-inducible BFP2 expression cassette, into the *LEU2* locus of strains *GAP1*_WT_-*yEGFP* and *GAP1*_syn_-*yEGFP* respectively.

### Mating, sporulation and tetrad isolation

Diploid yeast strains were generated by streaking strains onto YPD agar, incubating at 30 °C for 2 days and then mixing patches of colonies from cells of opposite mating type together on a fresh YPD agar plate. Cells were incubated for 4 hours at 30 °C before restreaking onto a fresh media plate with appropriate selection.

To set up sporulation cultures of diploid strains, cells were grown for 24 hours in pre-sporulation medium (10 g l^-1^ yeast extract, 20 g l^-1^ peptone, 10 g l^-1^ potassium acetate) before being washed twice in water and then resuspended in sporulation medium (10 g l^-1^ potassium acetate, 0.35 g l^-1^ yeast synthetic dropout medium and required amino acids supplemented to 0.25 x the amount added to synthetic complete medium). Sporulation cultures were incubated at 30 °C for 1-5 days until spore formation was visible under a microscope.

To isolate haploid strains, 200 µl sporulated cells were washed and resuspended in 200 µl water with LongLife Zymolyase (G-Biosciences). Cells were incubated at 37 °C for 10-20 minutes before 800 µl water was added to the cells. Digested cells were plated onto a YPD plate and tetrads were dissected to individual spores arrayed on the plate using a SporePlay+ tetrad dissection microscope (Singer Instruments). Mating type of isolated haploids was determined by PCR as previously described (Huxley *et al*., 1990).

### Plasmids

The plasmids used and generated in this study are listed in **Table S6**.

pRS403::*TRT2* tRNA complementing integrative vector - We PCR amplified the *TRT2* CDS, along with 380p upstream and 267 bp downstream sequence, using BY4741 genomic DNA as template with primers BB210/BB211. The PCR product was cloned into the multiple cloning site of pRS403 as an EcoRI/XmaI restriction fragment.

pSXI_3_36_M1 edited M1 chunk vector - We performed two PCR amplifications with pSXI_3_34_M1 template DNA, using primer pairs BB286/BB287 and BB288/BB289. The products were gel purified, pooled and used as template for a PCR reaction with primers BB286 and BB289. The 831 bp product containing the desired TAG>TAA change in the *SFT1* CDS was purified and assembled into the 12870 bp NdeI digest fragment of pSXI_3_34_M1 by Gibson isothermal assembly (Gibson *et al*., 2009).

pSXI_3_36_M3 edited M3 chunk vector - We PCR amplified four fragments from pSXI_3_34_M3 using primer pairs BB290/BB291, BB292/BB293, BB294/BB295 and BB296/297. The BB290/BB291and BB292/BB293 products were combined and used as template for PCR amplification using primers BB290 and BB293, yielding a 2496 bp product. The BB294/BB295 and BB296/297 products were combined and used as template for PCR amplification using primers BB294 and BB297, yielding a 1812 bp product. Both fragments, containing the TAG>TAA recoded *ECM9* and *YKR005C* regions, were purified and assembled into the 10639 bp SalI digest fragment of pSXI_3_34_M3 by Gibson isothermal assembly.

pSXI_3_37_M2 edited M2 chunk vector - We PCR amplified 4 fragments from a pSXI_3_34_M2 template. Fragment 1 was amplified with primers BB369/BB370, fragment 2 was amplified with primers BB371/BB372, fragment 3 was amplified with primers BB367/BB373 and fragment 4 was amplified with primers BB374/BB368. PCR with primers BB367/BB368 using a mixture of fragments 3 and 4 generated fragment 5, a 530 bp sequence covering *CEN11* with the downstream loxPsym site removed. Fragments 1 and 2 were ligated together using T4 DNA ligase to generate fragment 6, a 2050 bp sequence including the region 3’ of *YKR001C* CDS with a loxPsym sequence insertion. The 514 bp NsiI/SpeI restriction product of fragment 5, the 1998 bp SpeI/SexAI restriction product of fragment 6 and the 7535 bp restriction product of pSXI_3_34_M2 were ligated together with T4 DNA ligase to create pSXI_3_37_M2.

p*CEN11*_3_37b *CEN11* variant plasmid - We PCR amplified pSXI_3_34_M2 template DNA with primers BB468/BB472 to yield a 511 bp fragment, and with primers BB469/BB473 to yield a 286 bp fragment. These PCR products were ligated together with T4 DNA ligase and the ligation product was PCR amplified with primers BB468/BB469 to yield a 798 bp fragment, which was cloned into the 9285 bp NsiI/SalI pSXI_3_34_M2 restriction digest product as an NsiI/SalI restriction fragment.

p*CEN11*_3_37c *CEN11* variant plasmid - We PCR amplified p*CEN11*_3_37b template with primers BB468/BB475 to yield a 193 bp product, and with primers BB469/BB476 to yield a 571 bp product. These PCR products were ligated together with T4 DNA ligase and the ligation product was PCR amplified with primers BB468/BB469 to yield a 765 bp fragment, which was cloned into the 9285 bp NsiI/SalI pSXI_3_34_M2 restriction digest product as an NsiI/SalI restriction fragment.

p*CEN11*_3_37f *CEN11* variant plasmid - We PCR amplified pSXI_3_34_M2 template DNA with primers BB468/BB481 to yield a 511 bp fragment, and with primers BB469/BB479 to yield a 253 bp fragment. These PCR products were ligated together with T4 DNA ligase and the ligation product was PCR amplified with primers BB468/BB469 to yield a 764 bp fragment, which was cloned into the 9285 bp NsiI/SalI pSXI_3_34_M2 restriction digest product as an NsiI/SalI restriction fragment.

p*CEN11\**, containing *CEN11* region with *PGAL1* and a *K. lactis URA3* expression cassette - The region upstream of *CEN11* was PCR amplified from BY4741 genomic DNA with primers BB433/BB434; the *K. lactis URA3* cassette was amplified from pJJH1304 with primers BB435/BB436; *PGAL1* was amplified from BY4741 genomic DNA with primers BB437/BB438; and *CEN11* and its downstream region were amplified with primers BB439/BB440. The *CEN11* upstream and *K. lactis URA3* PCR products were pooled and used as template for PCR with primers BB433/BB436, yielding a 2035 bp product. The *PGAL1* and *CEN11* downstream PCR products were pooled and used as template for PCR with primers BB437/BB440, yielding a 1010 bp product. In a final PCR step, the 2035 bp and 1010 bp products were pooled and amplified with primers BB433/BB440 to give a 2999 bp product encoding the modified *CEN11* region. The assembled PCR product was ligated into the multiple cloning site of pUC19 as an EcoRI/HindIII restriction fragment.

pRS405-*LEU2*::*URA3* marker swapper construct plasmid - *URA3* was PCR amplified from pRS406 with primers oLM396/oLM397 and cloned into pRS405 as an AflII fragment.

pSXI_3_34_J4::*kanMX*, J4 chunk plasmid with *kanMX4* insertion - We modified pSXI_3_34_J4 to incorporate a *kanMX4* marker within the J4 chunk sequence by PCR amplifying the *kanMX4* marker from the genomic DNA of strain Y07039 using primers BB570 and BB571. The 1460bp product was cloned into pSXI_3_34_J4 as an XbaI/NarI restriction fragment, replacing 15 bp of *YKL053C-A*, generating plasmid pSXI_3_34_J4::*kanMX*.

pYZ412 *MATα* expression vector - the 3.4 kb *MATα* locus from pXZX353 (Xie *et al*., 2018) was subcloned into pRS415(Sikorski and Hieter, 1989).

p*SCW11*-*cre*-EBD-*kanMX4* SCRaMbLE plasmid with *kanMX4* marker - The vector was assembled using MoClo-YTK assembly (Lee *et al*., 2015) with plasmids pYKT083 (*AmpR*-*ColE1*), pYTK003 (ConL1), pYTK051 (*TENO1*), pYTK067 (ConR1), pYTK077 (*kanMX4*), pYTK081 (*CEN6*/*ARS4*) and plasmids with parts cloned from p*SCW11*-*cre*-EBD (Cai *et a*l., 2015) encoding the *SCW11* promoter (pJCH021) and the *cre*-EBD CDS (pJCH022).

pXL007, with a tetracycline inducible mScarlet reporter in a *LEU2* integraton cassette - The vector was assembled using MoClo-YTK assembly (Lee *et al*., 2015) and had a *LEU2* integration cassette containing *PRAD27*-[*TetA*-Nuclear Localisation Signal-*GAL4* activation domain]-*TADH1* Tet-On cassette, a tetO_7_-*PPHO5*-*mScarlet*-*TTDH1* fluorescent reporter cassette and *LEU2* marker on ColE1-*kanR* backbone.

pXL008, with a tetracycline inducible BFP reporter in a *LEU2* integraton cassette - The vector was assembled using MoClo-YTK assembly (Lee *et al*., 2015) and had a *LEU2* integration cassette containing *PRAD27*-[*TetA*-Nuclear Localisation Signal-*GAL4* activation domain]-*TADH1* Tet-On cassette, a tetO_7_-*PPHO5*-*BFP2*-*TTDH1* fluorescent reporter cassette and a *LEU2* marker cassette on a *ColE1*-*kanR* backbone.

### Online databases and resources

Unless otherwise stated, native *S. cerevisiae* CDSs and other DNA sequence elements were defined according to their annotations in the *Saccharomyces* Genome Database (Cherry *et al*., 2012). The basic local alignment search tool (BLAST; Altschul *et al*., 1990) was used to align DNA sequences against sequence databases (Karsch-Mizrachi *et al*., 2018).

### DNA extraction

Yeast genomic DNA for PCR screening was extracted using the GC prep method (Blount *et al*., 2016). Yeast genomic DNA for genome sequencing was extracted using Genomic-tip kits (Qiagen). Plasmid DNA was isolated from bacterial hosts using QIAprep Spin Miniprep kits (Qiagen).

### DNA Transformations

Linear DNA for chromosomal integration and plasmid DNA was transformed into yeast recipient cells using the lithium acetate method, as previously described (Annaluru *et al*., 2014). Cells underwent heat shock at 42 °C for 14 minutes and a 10 minute recovery step in 5 mM CaCl_2_ prior to plating on appropriate media. Plasmid DNA was introduced to *E. coli* recipient cells via electroporation with a MicroPulser (Bio-Rad).

### Synthetic chromosome assembly

Chunk DNA was released from the plasmid backbone through restriction digest at the designed nonpalindromic cutting sites (fig 1A), separated through 1% agarose gel electrophoresis, selectively excised from the gel and purified using a QIAquick Gel Extraction kit (Qiagen). Chunks constituting each megachunk were ligated together *in vitro* with T4 DNA ligase (New England Biolabs) overnight at 16 °C and then concentrated in a Concentrator Plus (Eppendorf) prior to transformation into the recipient strain. Transformant colonies were phenotypically selected for gain of the new auxotrophic marker and loss of the previous marker prior to PCRTag analysis. In this way, native chromosomal DNA was sequentially replaced with synthetic DNA following the SwAP-In approach (Dymond *et al*., 2011; Richardson *et al*., 2017).

In completing megachunk M, we had difficulty isolating a transformant with successful integration of the locus at the chunk M4-M5 junction (at gene *TOF2*), presumably due to inefficient *in vitro* restriction and ligation between DNA fragments. To integrate the missing synthetic DNA of this locus into a strain with a partial M integration, ysXIa16, we inserted the marker swapper construct *LEU2*::*URA3* into the chromosomal *LEU2* marker that was introduced by the prior megachunk M integration. This generated strain ysXIa17 with a functional *URA3* gene and a disrupted *LEU2*. We then ligated chunks M4 and M5 together *in vitro* and the full-length ligation product was purified and transformed into ysXIa17. We screened for auxotrophies, selecting colonies for gain of *LEU2* and loss of *URA3*.

### PCRTag analysis

Genomic DNA from megachunk transformant colonies with the correct auxotrophic profile was PCR screened for gain of synthetic DNA and loss of the corresponding wild type DNA. This was done using PCRTag primers, which target the synthetic PCRTag watermarks and their wild type equivalents (**Table S3**). Colonies confirmed to have gained all PCRTag sequences and lost all equivalent wild type sequences were considered to be successful megachunk integrants and progressed to the next round of megachunk integration. Following megachunk integration, successful megachunk integrants underwent spot assays on YPG media at 30 °C and YPD media at 30 °C and 37 °C.

### Growth spot assays

Saturated overnight yeast cultures were used to inoculate 5 ml YPD cultures. Cultures were grown to mid-exponential phase, normalised to an OD_600_ of 1, pelleted by centrifugation, washed in water, pelleted again and resuspended in water. Washed normalised cells were serially diluted in water in one-in-ten steps. Diluted cells were plated in 10 μl spots onto media plates and incubated at the appropriate temperature for the assay.

### CRISPR/Cas9 genome editing and debugging

Chromosomal editing using CRISPR/Cas9 was carried out using a previously described gap-repair vector system (Shaw *et al*., 2019). Target sequences were identified using the Benchling guide RNA design tool (http://www.benchling.com). To edit pWS082 to encode a retargeted gRNA, we amplified the vector using the phosphorylated primer BB353 and the desired retargeting primer, consisting of a 3’ sequence to bind pWS082 and a 5’ sequence encoding the retargeted gRNA region (**Table S2**). The exception is for the gRNAs targeted to *YKRCδ11, YKRCδ12* and the *yEGFP* insertion site upstream of *GAP1*, which were targeted using annealed oligonucleotides as previously described (Shaw *et al*., 2019). The PCR product was treated with DpnI at 37 °C for 1 hour to remove any template material, isolated by agarose gel electrophoresis, excised and then purified with a QIAquick Gel Extraction kit. The purified retargeted linear vector was self-ligated using T4 DNA ligase, which was then heat inactivated. The gRNA vector piece for transformation into the recipient cell was generated by PCR amplifying the circularised retargeted vector with primers BB421 and BB422. The product underwent agarose gel electrophoresis and purification and was co-transformed into the recipient cell along with the BsmBI restriction fragment of the CRISPR/Cas9 plasmid and the repair template. For the specific CRISPR/Cas9 target sites, repair templates and primers see **Table S4**.

### Pulsed-field gel electrophoresis

Samples for pulsed-field gel electrophoresis were prepared using a CHEF Yeast Genomic DNA Plug Kit (Bio-Rad) with lyticase from *Arthrobacter luteus* (Sigma-Aldrich) and recombinant proteinase K (Roche). Samples were run on a 1% certified megabase agarose (Bio-Rad) in TAE gel using a CHEF-DR III Pulsed Field Electrophoresis System (Bio-Rad) for 24 hours, at 6V cm^-1^ with a 60-120 second switch time ramp at an included angle of 120°. DNA was visualised under UV light following staining for 30 minutes with 0.5 µg ml^-1^ GelRed Nucleic Acid Stain (Millipore) and destaining in deionised water for 1 hour.

### Mitochondria assays

To determine whether loss of mitochondrial function affects cell growth and viability in a strain, cells were grown overnight and then incubated for 24 hours at 30 °C with or without the addition of ethidium bromide, a mitochondrial DNA depletion agent (Slonimski *et al*., 1968), to a final concentration of 10 µg ml^-1^. cells were then diluted to an OD600 of 0.001 and then plated onto YPD agar.

To determine whether cells were ρ^0^, we performed PCRs on genomic DNA templates with primer pairs targeting the mitochondrial 15S ribosomal RNA encoding *15S_RRNA*/*YNCQ0002W* (oLM394/oLM395) and the mitochondrial gene *COX2* (oLM398/oLM399). We visualised the PCR products by agarose gel electrophoresis. Product bands (634 bp for the 15S rRNA and 602 bp for *COX2*) were indicative of the mitochondrial genome being present in cells.

### Ploidy determination

To assess whether strains were haploid, they were first patched onto YPD plates and incubated at 30 °C for 2 days. The plates were then replica plated onto SC-Arg plates with and without L-canavanine sulphate (Sigma-Aldrich) added to a final concentration of 60 µg ml^-1^. As survival on L-canavanine is reliant on mutation of the CAN1 gene, a spontaneously acquired recessive trait (Whelan *et al*., 1979), strains with colony growth L-canavanine were assumed to be haploid.

### Genome sequencing

For Illumina MiSeq genome sequencing, yeast genomic DNA was quantified by Qubit fluorometry using a dsDNA HS Assay Kit (Thermo). For strain ysXIb01, library prep and sequencing was performed by BaseClear BV. For other strains, whole genome sequencing libraries were generated using the NEBNext Ultra II FS DNA Library Prep Kit (NEB) and sequenced using an Illumina NextSeq 500/550 High Output Kit v2.5 (75 Cycles). The Illumina MiSeq sequencing data for ysXIb01 was analysed using the Perfect Match Genomic Landscape strategy, as previously described (Palacios-Flores *et al*., 2018). Sequencing data for other strains was analysed using the Synthetic Yeast sequencing pipeline (Stracquadanio, G. *et al*, in preparation). Read coverage over genomic loci was determined and plotted as previously described (Zhao *et al*.).

Nanopore sequencing and analysis was performed as previously described (Blount *et al*., 2018).

### Transcript analysis

We grew cells in YPD or YPG medium at 30 °C until mid-exponential growth phase (OD_600_ ∼2). Cell culture corresponding to ∼3 × 10^8^ cells was harvested by centrifugation, washed in 0.8% physiological salt solution and resuspended in 500 µl solution of 1M sorbitol and 100 mM ethylenediaminetetraacetic acid (EDTA). We generated spheroplasts by digesting cells with 50U zymolyase (Zymo Research) at 30 °C for 30 minutes. We then collected spheroplasts by centrifugation and isolated RNA using the NucleoSpin RNA Plus kit (Macherey-Nagel). We evaluated RNA quality and integrity by Qubit fluorometry using an RNA BR Assay Kit (Thermo), spectrophotometry with a NanoDrop (Thermo) and on a 2100 Bioanalyser using an RNA 2000 Nano Kit (Agilent). RNA sequencing was performed by Novogene Co. Briefly, mRNA was purified from total RNA with poly-T oligo-attached magnetic beads prior to cDNA synthesis, adaptor ligation and sequencing on an Illumina platform.

RNA sequencing data was analysed using a custom pipeline. First, we pre-processed the Illumina unstranded paired-end reads by trimming adapters and remove low quality bases. Then, we built a reference *synXI* genome by replacing the wild type BY4742 *chrXI* with the *synXI*_9.11 and created a reference transcriptome by considering only protein-coding genes. Importantly, when a gene in the synthetic chromosome was deleted, we replaced it with the corresponding wild type one; this allows us to readily cross check for sample mislabelling, since no expression is expected from a deleted locus.

We then used transcriptomes and reads to quantify gene expression using kallisto quant with sequence based bias correction (Bray *et al*., 2016). Successively, differentially expressed genes were identified with edgeR (Robinson *et al*., 2010), using the exactTest method with dispersion parameter set to 0.22 to account for the lack of replicates. Finally, we reported as significantly differentially expressed all the genes with False Discovery Rate (FDR) less or equal to 0.01.

### Growth curves

Overnight YPD cultures were harvested, washed and used to inoculate 100 µl cultures in a 96-well plate with a starting OD600 normalised to 0.02. Plates were incubated and measured in a Synergy HT Microplate Reader (Biotek) shaking at 30 °C. Mean absorbance values of equivalent blank media wells were subtracted from data points.

### Competitive growth assays

Strains were grown overnight in SC--His-Leu-Ura-Trp medium, normalised for OD_600_, washed and resuspended in water. Each competing strain was inoculated to an initial OD_600_ of 0.2 in 500 µl of media in a 96-deep well plate. Each co-culture competition assay was performed in 3 independently inoculated wells to give biological triplicate data. Co-cultures were sub-cultured 5 µl into 500 µ fresh media in a new 96-deep well plate every 48 hours. At sub-culturing points, 100 µl of each culture was also transferred into a well in a 96-well plate containing fresh media and induced with 1 µM anhydrotetracycline (aTc) for 6 hours. The ratio of each fluorescent population in the co-culture was determined using flow cytometry.

### Flow cytometry

The fluorescence of the cells was measured by an Attune NxT Flow Cytometer (Thermo Scientific) with the following settings for measuring the size of the cell, complexity of the cell, yeGFP, BFP and m-Scarlet: FSC 100 V, SSC 355 V, BL1 450 V, VL1 345 V, YL2 510 V. 10,000 events were collected for each experiment and analysed by FlowJo.

### SCRaMbLE at the *GAP1* locus

Strain *GAP1*_syn_-*yEGFP*, transformed with plasmid p*SCW11*-*cre*EBD-*kanMX*4 was grown overnight in YPD media supplemented with 200 µg ml^-1^ G418S. Culture was diluted to an OD600 of 0.2 in 5 ml YPD media supplemented with 200 µg/mL G418S and grown for 4 hours. SCRaMbLE was induced by addition of β-estradiol to a final concentration of 1 µM. Cultures were grown for a further 2 hours before being washed twice in water and resuspended in 5 ml YPD. Cells were diluted x 10^−3^ in YPD and plated onto YPD agar plates. Plates were incubated at 30 °C for 3 days. Colonies were analysed by eye under blue light and those with increased GFP expression underwent screening to detect rearrangements at the *GAP1* locus. PCR analysis of colonies using exhaustive combinations of primers BB582, BB585, XL217, XL788, XL789, XL790, XL808 and XL809 was used to determine the structure of the *GAP1* locus for each strain.

### Fluorescence microscopy

Agarose pads were prepared as previously described (Skinner *et al*., 2013). For each sample, 2 μl of cell suspension was pipetted onto a coverslip of an imaging dish (idiTreat, µ-Dish 35 mm) and an agarose pad was placed on top. Cells in the imaging dish were visualised using a Nikon ECLIPSE Ti microscope by time-lapse imaging, with the following settings: Nosepiece, 20x, PFS on, interval 30 min, optical conf. BF and GFP, gain 1552.

## Supporting information

synXI_3.37 sequence

synXI_9.11 sequence

## Acknowledgements

We would like to thank Paul Freemont, Alistair Elfick, Ian Roberts and Anil Wipat for their support and advice, William Shaw for his help in establishing CRISPR gap-repair methods and Markus Ralser for his gift of plasmid pHLUM. This work was funded by BBSRC awards BB/K019791/1 and BB/R002614/1. B.A.B. was supported by the University of Nottingham through a Nottingham Research Fellowship.

## References

Alper, H., Fischer, C., Nevoigt, E., and Stephanopoulos, G. (2005). Tuning genetic control through promoter engineering. Proceedings of the National Academy of Sciences 102, 12678–12683. https://doi.org/10.1073/pnas.0504604102.

Altschul, S.F., Gish, W., Miller, W., Myers, E.W., and Lipman, D.J. (1990). Basic local alignment search tool. J. Mol. Biol. 215, 403–410.

Annaluru, N., Muller, H., Mitchell, L.A., Ramalingam, S., Stracquadanio, G., Richardson, S.M., Dymond, J.S., Kuang, Z., Scheifele, L.Z., Cooper, E.M., et al. (2014). Total synthesis of a functional designer eukaryotic chromosome. Science 344, 55–58.

Arrey, G., Keating, S.T., and Regenberg, B. (2022). A unifying model for extrachromosomal circular DNA load in eukaryotic cells. Semin. Cell Dev. Biol. https://doi.org/10.1016/j.semcdb.2022.03.002.

Bakota, L., Brandt, R., and Heinisch, J.J. (2012). Triple mammalian/yeast/bacterial shuttle vectors for single and combined Lentivirus- and Sindbis virus-mediated infections of neurons. Mol. Genet. Genomics 287, 313–324.

Blount, B.A., Weenink, T., Vasylechko, S., and Ellis, T. (2012). Rational Diversification of a Promoter Providing Fine-Tuned Expression and Orthogonal Regulation for Synthetic Biology. PLoS ONE 7, e33279. https://doi.org/10.1371/journal.pone.0033279.

Blount, B.A., Driessen, M.R.M., and Ellis, T. (2016). GC Preps: Fast and Easy Extraction of Stable Yeast Genomic DNA. Sci. Rep. 6, 26863.

Blount, B.A., Gowers, G.-O.F., Ho, J.C.H., Ledesma-Amaro, R., Jovicevic, D., McKiernan, R.M., Xie, Z.X., Li, B.Z., Yuan, Y.J., and Ellis, T. (2018). Rapid host strain improvement by in vivo rearrangement of a synthetic yeast chromosome. Nat. Commun. 9, 1932.

Bohovych, I., Fernandez, M.R., Rahn, J.J., Stackley, K.D., Bestman, J.E., Anandhan, A., Franco, R., Claypool, S.M., Lewis, R.E., Chan, S.S.L., et al. (2015). Metalloprotease OMA1 Fine-tunes Mitochondrial Bioenergetic Function and Respiratory Supercomplex Stability. Sci. Rep. 5, 1–14.

Bray, N.L., Pimentel, H., Melsted, P., and Pachter, L. (2016). Near-optimal probabilistic RNA-seq quantification. Nat. Biotechnol. 34, 525–527.

Cai, Y., Agmon, N., Choi, W.J., Ubide, A., Stracquadanio, G., Caravelli, K., Hao, H., Bader, J.S., and Boeke, J.D. (2015). Intrinsic biocontainment: multiplex genome safeguards combine transcriptional and recombinational control of essential yeast genes. Proc. Natl. Acad. Sci. U. S. A. 112, 1803–1808.

Carroll, S.M., DeRose, M.L., Gaudray, P., Moore, C.M., Needham-Vandevanter, D.R., Von Hoff, D.D., and Wahl, G.M. (1988). Double minute chromosomes can be produced from precursors derived from a chromosomal deletion. Mol. Cell. Biol. 8, 1525–1533.

Cherry, J.M., Hong, E.L., Amundsen, C., Balakrishnan, R., Binkley, G., Chan, E.T., Christie, K.R., Costanzo, M.C., Dwight, S.S., Engel, S.R., et al. (2012). Saccharomyces Genome Database: the genomics resource of budding yeast. Nucleic Acids Res. 40, D700–D705.

deCarvalho, A.C., Kim, H., Poisson, L.M., Winn, M.E., Mueller, C., Cherba, D., Koeman, J., Seth, S., Protopopov, A., Felicella, M., et al. (2018). Discordant inheritance of chromosomal and extrachromosomal DNA elements contributes to dynamic disease evolution in glioblastoma. Nat. Genet. 50, 708–717.

Dillon, L.W., Kumar, P., Shibata, Y., Wang, Y.-H., Willcox, S., Griffith, J.D., Pommier, Y., Takeda, S., and Dutta, A. (2015). Production of Extrachromosomal MicroDNAs Is Linked to Mismatch Repair Pathways and Transcriptional Activity. Cell Rep. 11, 1749–1759.

Domitrovic, T., Kozlov, G., Freire, J.C.G., Masuda, C.A., da Silva Almeida, M., Montero-Lomeli, M., Atella, G.C., Matta-Camacho, E., Gehring, K., and Kurtenbach, E. (2010). Structural and Functional Study of Yer067w, a New Protein Involved in Yeast Metabolism Control and Drug Resistance. PLoS One 5, e11163.

Duan, Z., Andronescu, M., Schutz, K., McIlwain, S., Kim, Y.J., Lee, C., Shendure, J., Fields, S., Anthony Blau, C., and Noble, W.S. (2010). A three-dimensional model of the yeast genome. Nature 465, 363–367. https://doi.org/10.1038/nature08973.

Dymond, J.S., Richardson, S.M., Coombes, C.E., Babatz, T., Muller, H., Annaluru, N., Blake, W.J., Schwerzmann, J.W., Dai, J., Lindstrom, D.L., et al. (2011). Synthetic chromosome arms function in yeast and generate phenotypic diversity by design. Nature 477, 471–476.

Foat, B.C., Houshmandi, S.S., Olivas, W.M., and Bussemaker, H.J. (2005). Profiling condition-specific, genome-wide regulation of mRNA stability in yeast. Proc. Natl. Acad. Sci. U. S. A. 102, 17675–17680.

Fredens, J., Wang, K., de la Torre, D., Funke, L.F.H., Robertson, W.E., Christova, Y., Chia, T., Schmied, W.H., Dunkelmann, D.L., Beránek, V., et al. (2019). Total synthesis of Escherichia coli with a recoded genome. Nature 569, 514–518.

Garvie, C.W., and Wolberger, C. (2001). Recognition of Specific DNA Sequences. Molecular Cell 8, 937–946. https://doi.org/10.1016/s1097-2765(01)00392-6.

Gerber, A.P., Herschlag, D., and Brown, P.O. (2004). Extensive Association of Functionally and Cytotopically Related mRNAs with Puf Family RNA-Binding Proteins in Yeast. PLoS Biol. 2, e79.

Gibson, D.G., Young, L., Chuang, R.-Y., Venter, J.C., Hutchison, C.A., 3rd, and Smith, H.O. (2009). Enzymatic assembly of DNA molecules up to several hundred kilobases. Nat. Methods 6, 343–345.

Gibson, D.G., Glass, J.I., Lartigue, C., Noskov, V.N., Chuang, R.-Y., Algire, M.A., Benders, G.A., Montague, M.G., Ma, L., Moodie, M.M., et al. (2010). Creation of a bacterial cell controlled by a chemically synthesized genome. Science 329, 52–56.

Grant, S.G., Jessee, J., Bloom, F.R., and Hanahan, D. (1990). Differential plasmid rescue from transgenic mouse DNAs into Escherichia coli methylation-restriction mutants. Proc. Natl. Acad. Sci. U. S. A. 87, 4645–4649.

Gresham, D., and Hong, J. (2015). The functional basis of adaptive evolution in chemostats. FEMS Microbiol. Rev. 39, 2–16.

Gresham, D., Usaite, R., Germann, S.M., Lisby, M., Botstein, D., and Regenberg, B. (2010). Adaptation to diverse nitrogen-limited environments by deletion or extrachromosomal element formation of the GAP1 locus. Proc. Natl. Acad. Sci. U. S. A. 107, 18551–18556.

Hill, A., and Bloom, K. (1987). Genetic manipulation of centromere function. Mol. Cell. Biol. 7, 2397–2405.

Hull, R.M., King, M., Pizza, G., Krueger, F., Vergara, X., and Houseley, J. (2019). Transcription-induced formation of extrachromosomal DNA during yeast ageing. PLoS Biol. 17, e3000471.

Huxley, C., Green, E.D., and Dunham, I. (1990). Rapid assessment of S. cerevisiae mating type by PCR. Trends Genet. 6, 236.

Jauniaux, J.-C., and Grenson, M. (1990). GAP1, the general amino acid permease gene of Saccharomyces cerevisiae. Nucleotide sequence, protein similarity with the other bakers yeast amino acid permeases, and nitrogen catabolite repression. European Journal of Biochemistry 190, 39–44. https://doi.org/10.1111/j.1432-1033.1990.tb15542.x.

Karsch-Mizrachi, I., Takagi, T., Cochrane, G., and International Nucleotide Sequence Database Collaboration (2018). The international nucleotide sequence database collaboration. Nucleic Acids Res. 46, D48–D51.

Kitakawa, M., Grohmann, L., Graack, H.R., and Isono, K. (1990). Cloning and characterization of nuclear genes for two mitochondrial ribosomal proteins in Saccharomyces cerevisiae. Nucleic Acids Res. 18, 1521–1529.

Kreike, J., Schulze, M., Pillar, T., Körte, A., and Rödel, G. (1986). Cloning of a nuclear gene MRS1 involved in the excision of a single group I intron (bI3) from the mitochondrial COB transcript in S. cerevisiae. Current Genetics 11, 185–191. https://doi.org/10.1007/bf00420605.

Lauer, S., Avecilla, G., Spealman, P., Sethia, G., Brandt, N., Levy, S.F., and Gresham, D. (2018). Single-cell copy number variant detection reveals the dynamics and diversity of adaptation. PLoS Biol. 16, e3000069.

Lee, M.E., DeLoache, W.C., Cervantes, B., and Dueber, J.E. (2015). A Highly Characterized Yeast Toolkit for Modular, Multipart Assembly. ACS Synth. Biol. 4, 975–986.

Ling, X., Han, Y., Meng, J., Zhong, B., Chen, J., Zhang, H., Qin, J., Pang, J., and Liu, L. (2021). Small extrachromosomal circular DNA (eccDNA): major functions in evolution and cancer. Mol. Cancer 20, 113.

Liu, W., Luo, Z., Wang, Y., Pham, N.T., Tuck, L., Pérez-Pi, I., Liu, L., Shen, Y., French, C., Auer, M., et al. (2018). Rapid pathway prototyping and engineering using in vitro and in vivo synthetic genome SCRaMbLE-in methods. Nat. Commun. 9, 1936.

van Loon, N., Miller, D., and Murnane, J.P. (1994). Formation of extrachromosomal circular DNA in HeLa cells by nonhomologous recombination. Nucleic Acids Res. 22, 2447–2452.

Luo, Z., Wang, L., Wang, Y., Zhang, W., Guo, Y., Shen, Y., Jiang, L., Wu, Q., Zhang, C., Cai, Y., et al. (2018). Identifying and characterizing SCRaMbLEd synthetic yeast using ReSCuES. Nat. Commun. 9, 1930.

Luo, Z., Yu, K., Xie, S., Monti, M., Schindler, D., Fang, Y., Zhao, S., Liang, Z., Jiang, S., Luan, M., et al. (2021). Compacting a synthetic yeast chromosome arm. Genome Biol. 22, 5.

Marc, P., Margeot, A., Devaux, F., Blugeon, C., Corral-Debrinski, M., and Jacq, C. (2002). Genome-wide analysis of mRNAs targeted to yeast mitochondria. EMBO Rep. 3, 159–164.

Mercy, G., Mozziconacci, J., Scolari, V.F., Yang, K., Zhao, G., Thierry, A., Luo, Y., Mitchell, L.A., Shen, M., Shen, Y., et al. (2017). 3D organization of synthetic and scrambled chromosomes. Science 355. https://doi.org/10.1126/science.aaf4597.

Mitchell, L.A., Wang, A., Stracquadanio, G., Kuang, Z., Wang, X., Yang, K., Richardson, S., Martin, J.A., Zhao, Y., Walker, R., et al. (2017). Synthesis, debugging, and effects of synthetic chromosome consolidation: synVI and beyond. Science 355. https://doi.org/10.1126/science.aaf4831.

Møller, H.D., Andersen, K.S., and Regenberg, B. (2013). A model for generating several adaptive phenotypes from a single genetic event. Communicative & Integrative Biology 6, e23933. https://doi.org/10.4161/cib.23933.

Møller, H.D., Parsons, L., Jørgensen, T. S., Botstein, D., and Regenberg, B. (2015) Extrachromosomal circular DNA is common in yeast. Proc. Natl. Acad. Sci. U. S. A. 112 (24) E3114–E3122.

Mülleder, M., Capuano, F., Pir, P., Christen, S., Sauer, U., Oliver, S.G., and Ralser, M. (2012). A prototrophic deletion mutant collection for yeast metabolomics and systems biology. Nat. Biotechnol. 30, 1176–1178.

Noer, J.B., Hørsdal, O.K., Xiang, X., Luo, Y., and Regenberg, B. (2022). Extrachromosomal circular DNA in cancer: history, current knowledge, and methods. Trends Genet. https://doi.org/10.1016/j.tig.2022.02.007.

Palacios-Flores, K., García-Sotelo, J., Castillo, A., Uribe, C., Aguilar, L., Morales, L., Gómez-Romero, L., Reyes, J., Garciarubio, A., Boege, M., et al. (2018). A Perfect Match Genomic Landscape Provides a Unified Framework for the Precise Detection of Variation in Natural and Synthetic Haploid Genomes. Genetics 208, 1631–1641.

Paulsen, T., Kumar, P., Koseoglu, M.M., and Dutta, A. (2018). Discoveries of Extrachromosomal Circles of DNA in Normal and Tumor Cells. Trends Genet. 34, 270–278.

Pleiss, J.A., Whitworth, G.B., Bergkessel, M., and Guthrie, C. (2007). Transcript specificity in yeast pre-mRNA splicing revealed by mutations in core spliceosomal components. PLoS Biol. 5, e90.

Richardson, S.M., Mitchell, L.A., Stracquadanio, G., Yang, K., Dymond, J.S., DiCarlo, J.E., Lee, D., Huang, C.L.V., Chandrasegaran, S., Cai, Y., et al. (2017). Design of a synthetic yeast genome. Science 355, 1040–1044.

Robinson, M.D., McCarthy, D.J., and Smyth, G.K. (2010). edgeR: a Bioconductor package for differential expression analysis of digital gene expression data. Bioinformatics 26, 139–140.

Saint-Georges, Y., Garcia, M., Delaveau, T., Jourdren, L., Le Crom, S., Lemoine, S., Tanty, V., Devaux, F., and Jacq, C. (2008). Yeast Mitochondrial Biogenesis: A Role for the PUF RNA-Binding Protein Puf3p in mRNA Localization. PLoS One 3, e2293.

Sertil, O., Cohen, B.D., Davies, K.J., and Lowry, C.V. (1997). The DAN1 gene of S. cerevisiae is regulated in parallel with the hypoxic genes, but by a different mechanism. Gene 192, 199–205.

Shaw, W.M., Yamauchi, H., Mead, J., Gowers, G.-O.F., Bell, D.J., Öling, D., Larsson, N., Wigglesworth, M., Ladds, G., and Ellis, T. (2019). Engineering a Model Cell for Rational Tuning of GPCR Signaling. Cell 177, 782–796.e27.

Shen, Y., Stracquadanio, G., Wang, Y., Yang, K., Mitchell, L.A., Xue, Y., Cai, Y., Chen, T., Dymond, J.S., Kang, K., et al. (2016). SCRaMbLE generates designed combinatorial stochastic diversity in synthetic chromosomes. Genome Res. 26, 36–49.

Shen, Y., Wang, Y., Chen, T., Gao, F., Gong, J., Abramczyk, D., Walker, R., Zhao, H., Chen, S., Liu, W., et al. (2017). Deep functional analysis of synII, a 770-kilobase synthetic yeast chromosome. Science 355. https://doi.org/10.1126/science.aaf4791.

Sikorski, R.S., and Hieter, P. (1989). A system of shuttle vectors and yeast host strains designed for efficient manipulation of DNA in Saccharomyces cerevisiae. Genetics 122, 19–27.

Skinner, S.O., Sepúlveda, L.A., Xu, H., and Golding, I. (2013). Measuring mRNA copy number in individual Escherichia coli cells using single-molecule fluorescent in situ hybridization. Nat. Protoc. 8, 1100–1113.

Slonimski, P.P., Perrodin, G., and Croft, J.H. (1968). Ethidium bromide induced mutation of yeast mitochondria: Complete transformation of cells into respiratory deficient non-chromosomal “petites.” Biochemical and Biophysical Research Communications 30, 232–239. https://doi.org/10.1016/0006-291x(68)90440-3.

Wang, H., Li, Q., Zhang, Z., Zhou, C., Ayepa, E., Abrha, G.T., Han, X., Hu, X., Yu, X., Xiang, Q., et al. (2019). YKL107W from Saccharomyces cerevisiae encodes a novel aldehyde reductase for detoxification of acetaldehyde, glycolaldehyde, and furfural. Appl. Microbiol. Biotechnol. 103, 5699–5713.

Wang, Y., Wang, M., Djekidel, M.N., Chen, H., Liu, D., Alt, F.W., and Zhang, Y. (2021). eccDNAs are apoptotic products with high innate immunostimulatory activity. Nature 599, 308–314.

Whelan, W.L., Gocke, E., and Manney, T.R. (1979). The CAN1 locus of Saccharomyces cerevisiae: fine-structure analysis and forward mutation rates. Genetics 91, 35–51.

Winzeler, E.A., Shoemaker, D.D., Astromoff, A., Liang, H., Anderson, K., Andre, B., Bangham, R., Benito, R., Boeke, J.D., Bussey, H., et al. (1999). Functional characterization of the S. cerevisiae genome by gene deletion and parallel analysis. Science 285, 901–906.

Wu, S., Turner, K.M., Nguyen, N., Raviram, R., Erb, M., Santini, J., Luebeck, J., Rajkumar, U., Diao, Y., Li, B., et al. (2019). Circular ecDNA promotes accessible chromatin and high oncogene expression. Nature 575, 699–703.

Wu, Y., Li, B.-Z., Zhao, M., Mitchell, L.A., Xie, Z.-X., Lin, Q.-H., Wang, X., Xiao, W.-H., Wang, Y., Zhou, X., et al. (2017). Bug mapping and fitness testing of chemically synthesized chromosome X. Science 355. https://doi.org/10.1126/science.aaf4706.

Xie, Z.-X., Li, B.-Z., Mitchell, L.A., Wu, Y., Qi, X., Jin, Z., Jia, B., Wang, X., Zeng, B.-X., Liu, H.-M., et al. (2017). “Perfect” designer chromosome V and behavior of a ring derivative. Science 355. https://doi.org/10.1126/science.aaf4704.

Xie, Z.-X., Mitchell, L.A., Liu, H.-M., Li, B.-Z., Liu, D., Agmon, N., Wu, Y., Li, X., Zhou, X., Li, B., et al. (2018). Rapid and Efficient CRISPR/Cas9-Based Mating-Type Switching of. G3 8, 173–183.

Xu, Z., Wei, W., Gagneur, J., Perocchi, F., Clauder-Münster, S., Camblong, J., Guffanti, E., Stutz, F., Huber, W., and Steinmetz, L.M. (2009). Bidirectional promoters generate pervasive transcription in yeast. Nature 457, 1033–1037.

Yu, C.-H., Dang, Y., Zhou, Z., Wu, C., Zhao, F., Sachs, M.S., and Liu, Y. (2015). Codon Usage Influences the Local Rate of Translation Elongation to Regulate Co-translational Protein Folding. Mol. Cell 59, 744–754.

Zhang, H., and Singh, K.K. (2014). Global genetic determinants of mitochondrial DNA copy number. PLoS One 9, e105242.

Zhao, Y., Coelho, C., Hughes, A.L., Lazar-Stefanita, L., Yang, S., Brooks, A.N., Walker, R.S.K., Zhang, W., Lauer, S., Hernandez, C., et al. Debugging and consolidating multiple synthetic chromosomes reveals combinatorial genetic interactions. https://doi.org/10.1101/2022.04.11.486913.

