## Supplementary Information for "Synthetic yeast chromosome XI design enables extrachromosomal circular DNA formation on demand"

### Supplementary figures

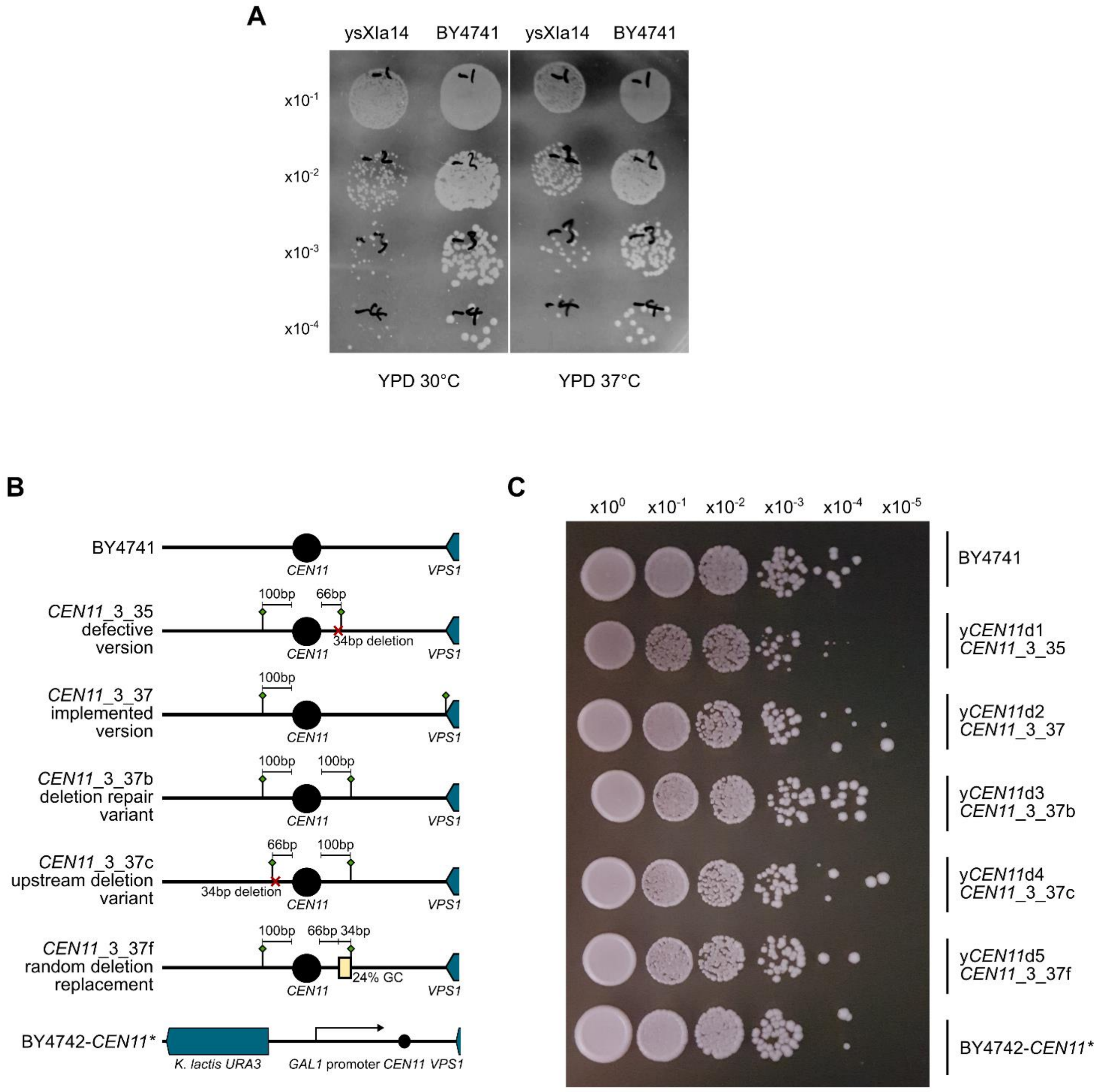


***Figure S1 - Supplementary to Figure 2.*** *(A) Growth spot assays of ysXIa14 and BY4741 parental control on YPD agar after 2 days. Incubation temperatures and spot dilution factors are indicated on the image. (B) Overview of the* CEN11 *locus variants generated in this study. Green diamonds denote loxPsym sites. (C) Growth spot assays of BY4742 strains with different* CEN11 *locus variants on YPD after 3 days growth at 37 °C. Strain names and* CEN11 *variant numbers (where appropriate) are shown to the right. Spot dilution factors are shown above the image.*

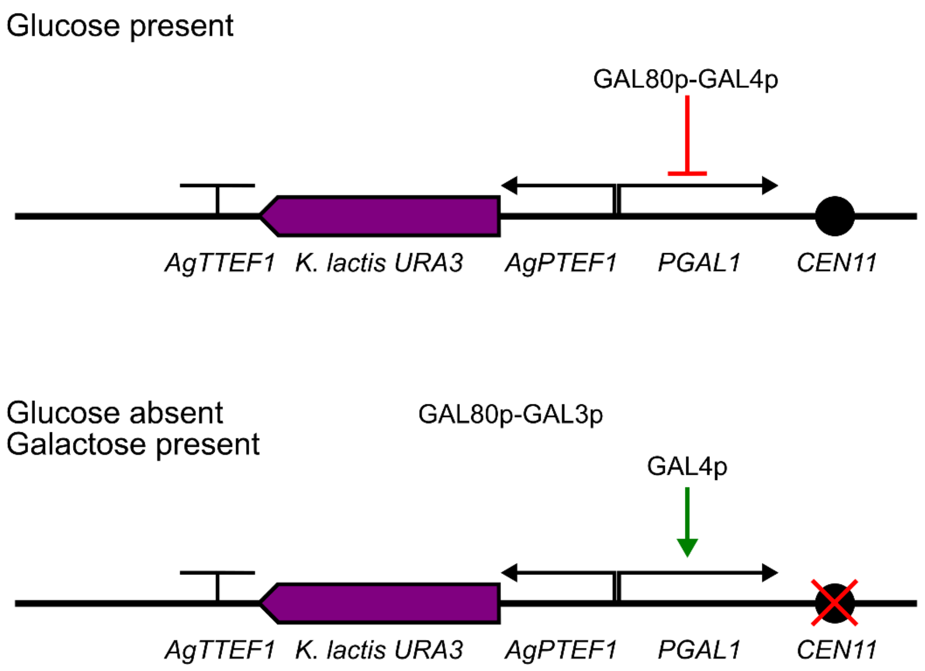


***Figure S2 - Overview of the*** **CEN11*** ***locus under glucose repression and galactose induction.*** *In the absence of glucose and the presence of galactose, transcription from the* GAL1 *promoter (*PGAL1*) disrupts the function of* CEN11*, leading to chromosome loss through miss-segregation in mitosis.* AgTTEF1 *is the* TEF1 *terminator from* Ashbya gossypii, K. lactis URA3 *is the* URA3 *coding sequence from* Kluyveromyces lactis *and* AgPTEF1 *is the* TEF1 *promoter from* A. gossypii*. The red bar indicates transcriptional repression, the green arrow represents transcriptional activation and the red cross represents disruption of centromere function.*

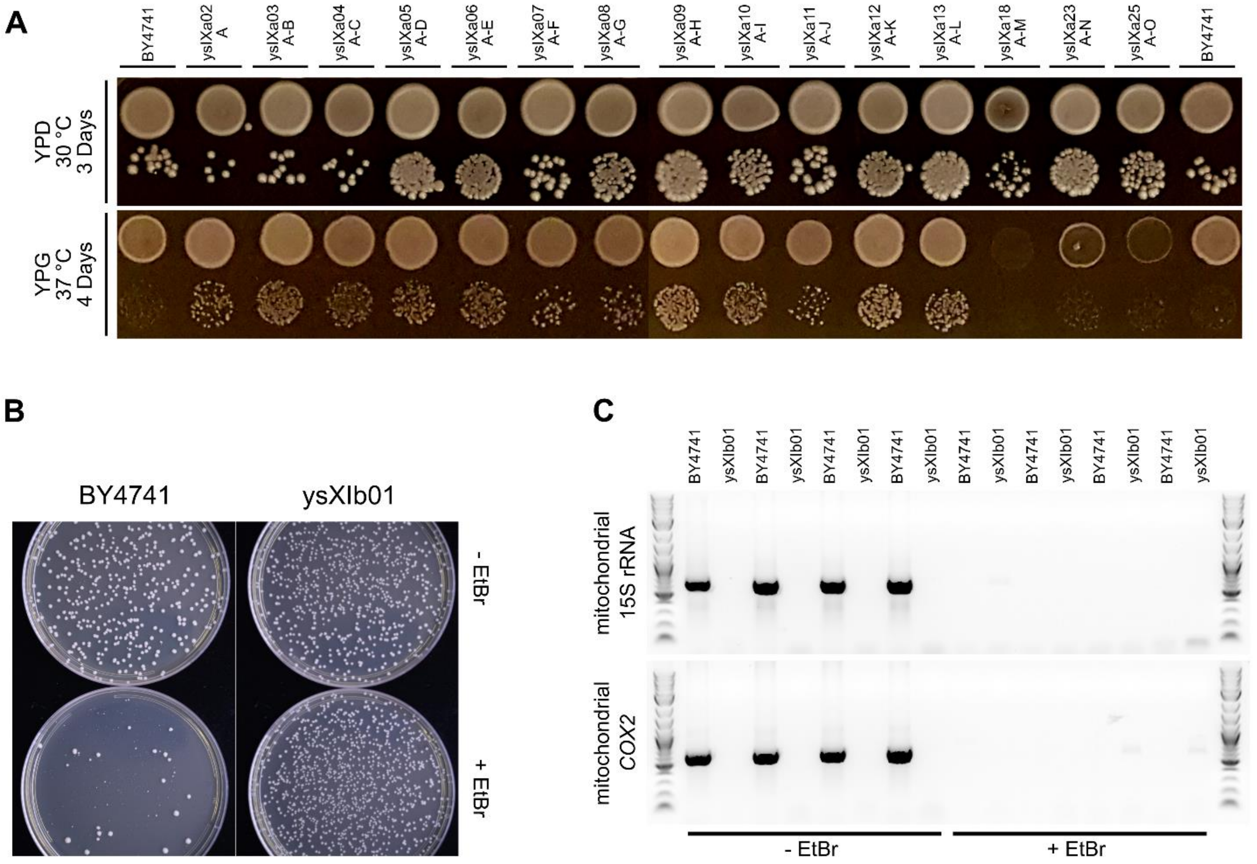


***Figure S5 – Supplementary to Figure 3.*** *(A) Growth spot assays of synXI assembly intermediates following rounds of megachunk integration on YPD (glucose) and YPG (glycerol) to test respiratory function. For each strain and condition, the top spot is a x10*$^{-1}$ *dilution and the bottom spot is a x10*$^{-3}$ *dilution. (B) Diluted cells plated onto YPD with or without 24 hours prior exposure 10 µg ethidium bromide (EtBr). EtBr causes loss of the mitochondrial genome. (C) PCR assays targeting mitochondrial genomic DNA for amplification performed on genomic DNA harvested from colonies grown on YPD or on YPD + 10 µg EtBr. Product bands indicate the presence of mitochondrial genomic DNA. n=4 biological replicates per strain and per condition. PCR products are visualised on 1% agarose gel, with the gel image colour inverted. Ladder is 1 kb Plus DNA Ladder (New England Biolabs).*

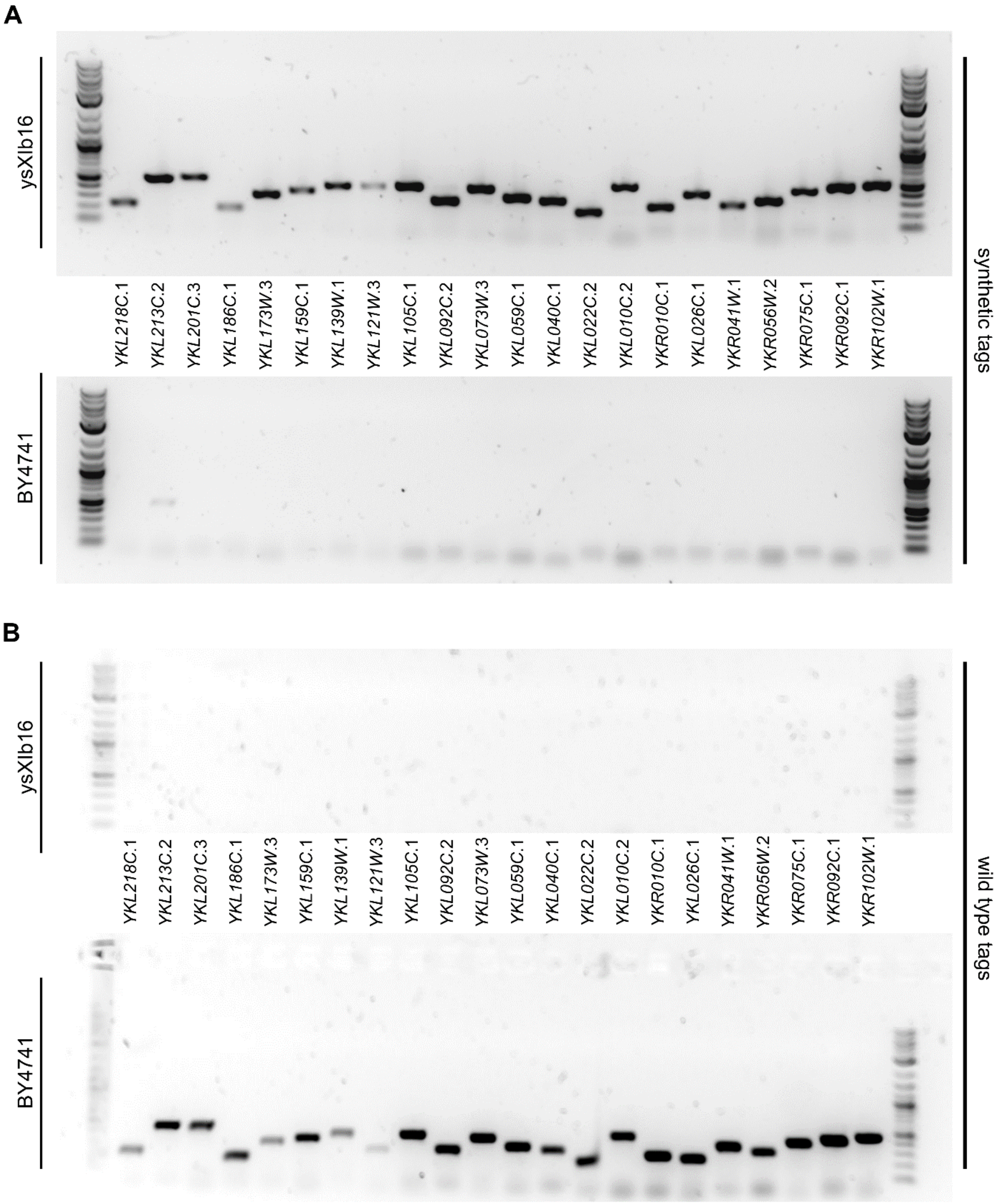


***Figure S6 – Supplementary to Figure 4.*** *(A) Genomic DNA from ysXIb16 or a BY4741 control was amplified with synthetic PCRTag primer pairs. Product bands indicate the presence of synthetic DNA. (B) Genomic DNA from ysXIb16 or a BY4741 control was amplified with wildtype PCRTag primer pairs. Product bands indicate the presence of wildtype DNA. For panels A and B, PCR products were visualised on 1% agarose gel, with the colour inverted. PCRTag primer pair targets are labelled between the gel images, with target sequences listed in Table S3. Ladder is 1 kb Plus DNA Ladder (New England Biolabs).*

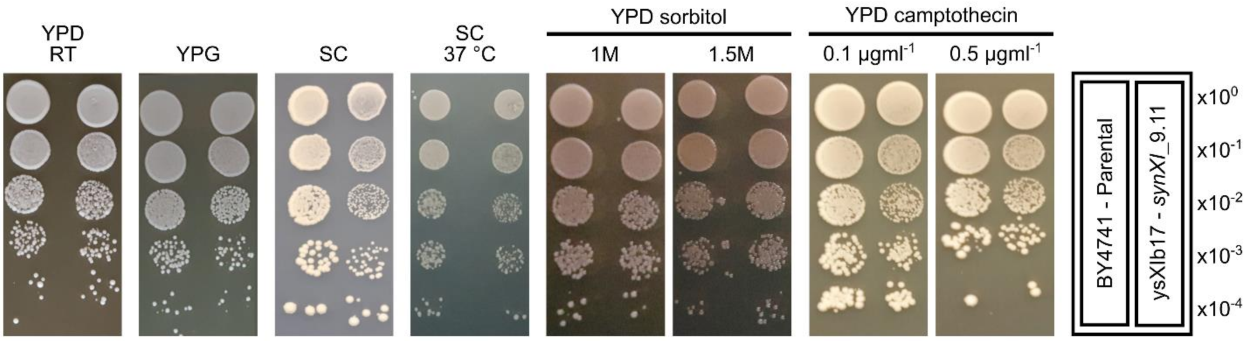


***Figure S7 – Supplementary to Figure 4.*** *Additional growth spot assays of ysXIb17 and a BY4741 parental control to assess cellular fitness. Serial dilutions of cultures were spotted as illustrated on the right. Unless otherwise indicated, plates were incubated at 30 °C.*

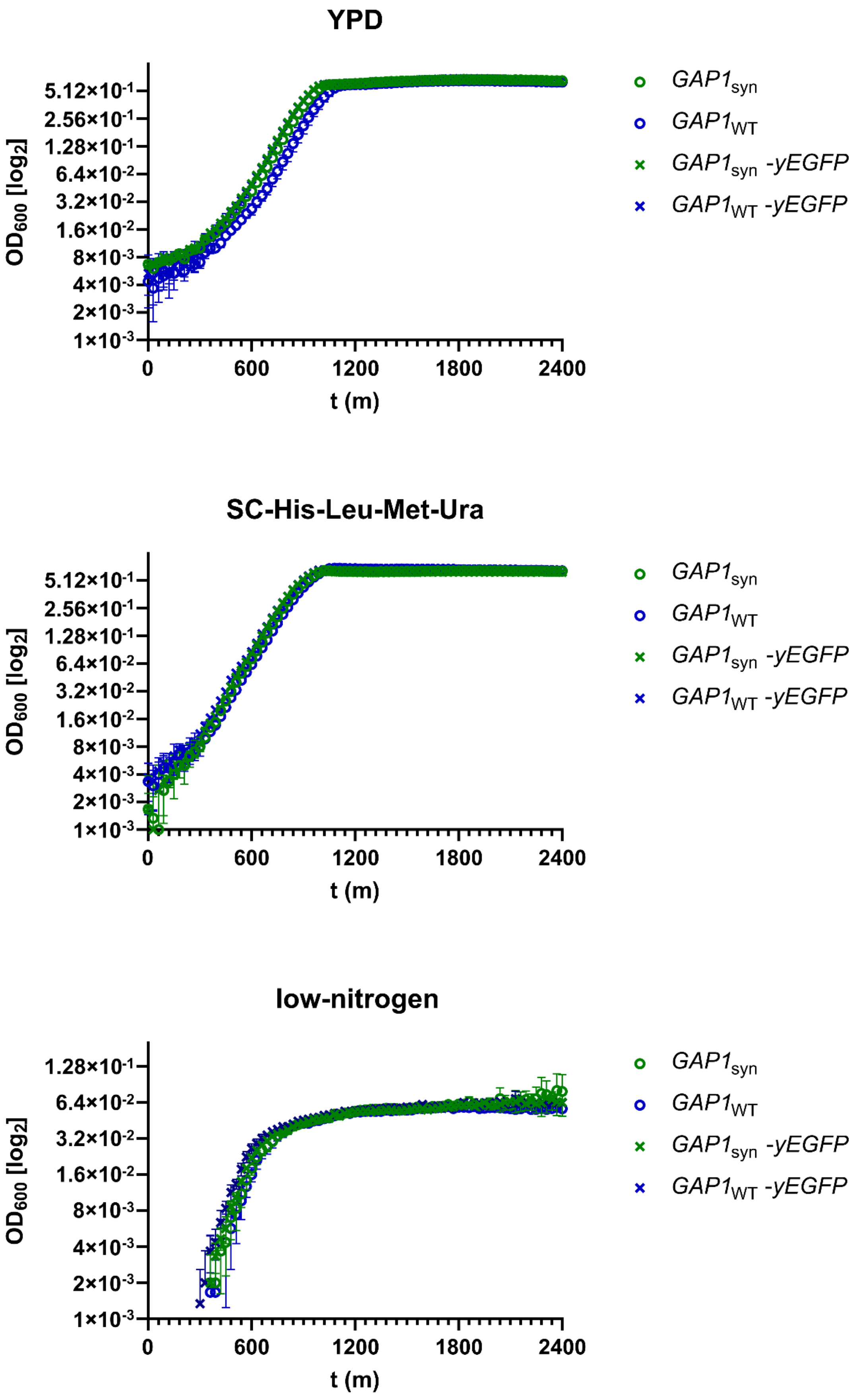


***Figure S8 – Supplementary to Figure 5.*** *Graphs show growth of BY4741 strains with different variations of the* GAP1 *locus in 96 well plates under rich (YPD), defined (SC-His-Leu-Met-Ura) and low-nitrogen media conditions at 30 °C. All strains had plasmid pHLUM, making them prototrophic. Mean* $OD_{600}$ *values from 3 biological replicates are plotted as circles, error bars represent standard deviation.*

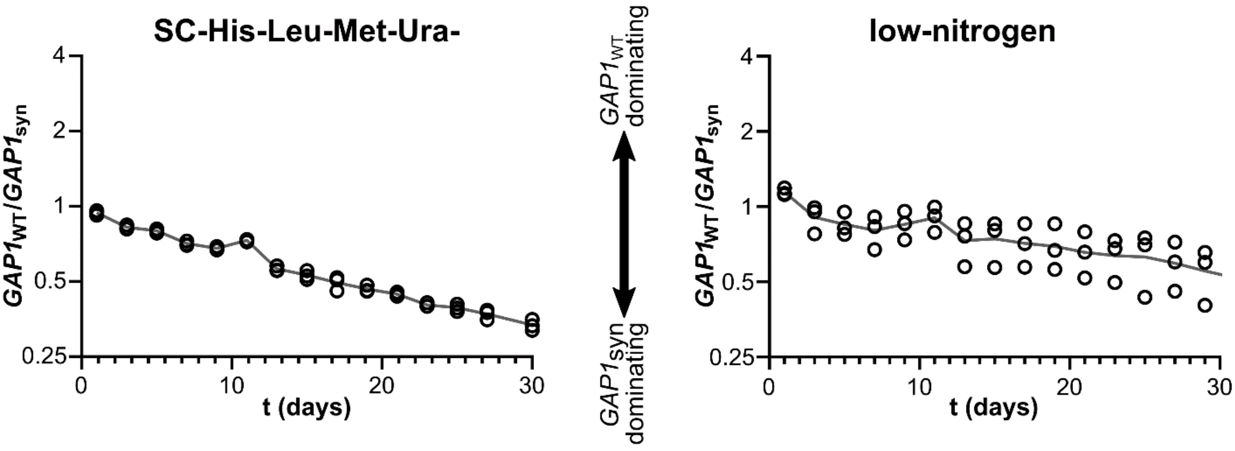


***Figure S9 – Supplementary to Figure 5.*** *Graphs show the ratio of BY4741*-GAP1$_{WT}$-yEGFP-BFP *to* GAP1$_{syn}$-yEGFP-mScarlet *cells under competitive growth sampled over the duration of the competition assays in synthetic and low-nitrogen media. Biological replicates (n=3) are plotted as circles and mean values are plotted as a line. Y axis has a log$_2$ scale.*

### Supplementary tables

| **Strain** | **Genotype** | **Description** | **Source** |
|---|---|---|---|
| BY4741 | *MAT**a** his3*Δ1 *leu2*Δ0 *met15*Δ0 *ura3*Δ0 | Referred to as parental | Brachmann *et al.*, 1998 |
| Y07039 | *MAT**a** his3*Δ1 *leu2*Δ0 *met15*Δ0 *ura3*Δ0 *YKL220C*Δ::*kanMX4* | BY4741 with YKL220C replaced by kanMX4 | EUROSCARF; Winzeler *et al., 1999* |
| ysXIa01 | *MAT**a** his3*Δ1::*HIS3*-pRS413-*TRT2 leu2*Δ0 *met15*Δ0 *ura3*Δ0 *YKL220C*Δ::*kanMX4* | Starting strain for *synXI* assembly | This study |
| ysXIa02 | *MAT**a** his3*Δ1::*HIS3*-pRS413-*TRT2 leu2*Δ0 *met15*Δ0 *ura3*Δ0 *YKL219W*::*LEU2* | *synXI* intermediate strain with megachunk A | This study |
| ysXIa03 | *MAT**a** his3*Δ1::*HIS3*-pRS413-*TRT2 leu2*Δ0 *met15*Δ0 *ura3*Δ0 *YKL205W*::*URA3* | *synXI* intermediate strain with megachunks A-B | This study |
| ysXIa04 | *MAT**a** his3*Δ1::*HIS3*-pRS413-*TRT2 leu2*Δ0 *met15*Δ0 *ura3*Δ0 *YKL187C*::*LEU2* | *synXI* intermediate strain with megachunks A-C | This study |
| ysXIa05 | *MAT**a** his3*Δ1::*HIS3*-pRS413-*TRT2 leu2*Δ0 *met15*Δ0 *ura3*Δ0 *YKL171W*::*URA3* | *synXI* intermediate strain with megachunks A-D | This study |
| ysXIa06 | *MAT**a** his3*Δ1::*HIS3*-pRS413-*TRT2 leu2*Δ0 *met15*Δ0 *ura3*Δ0 *YKL151C*::*LEU2* | *synXI* intermediate strain with megachunks A-E | This study |
| ysXIa07 | *MAT**a** his3*Δ1::*HIS3*-pRS413-*TRT2 leu2*Δ0 *met15*Δ0 *ura3*Δ0 *YKL129C*::*URA3* | *synXI* intermediate strain with megachunks A-F | This study |
| ysXIa08 | *MATa his3Δ1::HIS3-pRS413-TRT2 leu2Δ0 met15Δ0 ura3Δ0 YKL107W::LEU2* | *synXI* intermediate strain with megachunks A-G | This study |
| ysXIa09 | *MAT**a** his3*Δ1::*HIS3*-pRS413-*TRT2 leu2*Δ0 *met15*Δ0 *ura3*Δ0 *YKL091C*::*URA3* | *synXI* intermediate strain with megachunks A-H | This study |

| Strain | Genotype | Description | Source |
|---|---|---|---|
| ysXIa10 | *MAT**a** his3*Δ1::*HIS3*-pRS413-*TRT2 leu2*Δ0 *met15*Δ0 *ura3*Δ0 *YKL069W*::*LEU2* | *synXI* intermediate strain with megachunks A-I | This study |
| ysXIa11 | *MAT**a** his3*Δ1::*HIS3*-pRS413-*TRT2 leu2*Δ0 *met15*Δ0 *ura3*Δ0 *YKL048C*::*URA3* | *synXI* intermediate strain with megachunks A-J | This study |
| ysXIa12 | *MAT**a** his3*Δ1::*HIS3*-pRS413-*TRT2 leu2*Δ0 *met15*Δ0 *ura3*Δ0 *YKL025C*::*LEU2* | *synXI* intermediate strain with megachunks A-K | This study |
| ysXIa13 | *MAT**a** his3*Δ1::*HIS3*-pRS413-*TRT2 leu2*Δ0 *met15*Δ0 *ura3*Δ0 *YKL008C*::*URA3* | *synXI* intermediate strain with megachunks A-L | This study |
| ysXIa14 | *MAT**a** his3*Δ1::*HIS3*-pRS413-*TRT2 leu2*Δ0 *met15*Δ0 *ura3*Δ0 *YKR015C*::*LEU2* | *synXI* intermediate strain with megachunks A-M (3.36) | This study |
| BY4742 | *MATα his3*Δ1 *leu2*Δ0 *lys2*Δ0 *ura3*Δ0 | Referred to as parental | Brachmann *et al.*, 1998 |
| Y14868 | *MATα his3*Δ1 *leu2*Δ0 *lys2*Δ0 *ura3*Δ0 *YKL007W*Δ::*kanMX4* | BY4742 with *YKL007C* replaced by *kanMX4* | EUROSCARF; Winzeler *et al., 1999* |
| ysXIa15 | *MATα his3*Δ1 *leu2*Δ0 *lys2*Δ0 *ura3*Δ0 *YKR015C*::*LEU2* | *synXI* intermediate strain with megachunk M (3.36) | This study |
| ysXIa16 | *MAT**a** his3*Δ1::*HIS3*-pRS413-*TRT2 leu2*Δ0 *met15*Δ0 *ura3*Δ0 *YKR015C*::*LEU2* | *synXI* intermediate strain with megachunks A-M (partial, 3.37) | This study |
| ysXIa17 | *MAT**a** his3*Δ1::*HIS3*-pRS413-*TRT2 leu2*Δ0 *met15*Δ0 *ura3*Δ0 *YKR015C*::*LEU2*::*URA3* | *synXI* intermediate strain with megachunks A-M (partial, 3.37, marker swapped) | This study |
| ysXIa18 | *MAT**a** his3*Δ1::*HIS3*-pRS413-*TRT2 leu2*Δ0 *met15*Δ0 *ura3*Δ0 *YKR015C*::*LEU2* | *synXI* intermediate strain with megachunks A-M (3.37) | This study |
| ysXIa19 | *MAT**a**/α his3*Δ1::*HIS3*-pRS413-*TRT2*/*his3*Δ1 *leu2*Δ0/*leu2*Δ0 *LYS2*/*lys2*Δ0 *met15*Δ0/*MET15* *ura3*Δ0/*ura3*Δ0 *YKR015C*::*LEU2*/*YKR015C* *YKL007W*/*YKL007W*::*kanMX4* | Diploid of strains ysXIa18 and Y14868. | This study |
| ysXIa20 | *MAT**a**/α his3*Δ1/*his3*Δ1 *leu2*Δ0/*leu2*Δ0 *LYS2*/*lys2*Δ0 *met15*Δ0/*MET15 ura3*Δ0/*ura3*Δ0 *YKR015C*::*LEU2*/*YKR015C* *YKL007W*/*YKL007W*::*kanMX4* | ysXIa19 with *HIS3*-pRS413-*TRT2* integrated construct removed | This study |
| ysXIa21 | *MAT**a**/α his3*Δ1/*his3*Δ1 *leu2*Δ0/*leu2*Δ0 *LYS2*/*lys2*Δ0 *met15*Δ0/*MET15 ura3*Δ0/*ura3*Δ0 *YKR015C*::*LEU2*/*YKR015C* *YKL007W*/*YKL007W*::*kanMX4* pRS413-*chrXI_tRNA* (*HIS3*) | ysXIa20 with *chrXI* synthetic tRNA array plasmid | This study |
| ysXIa22 | *MAT**a** his3*Δ1 *leu2*Δ0 *met15*Δ0 *ura3*Δ0 *YKR015C*::*LEU2* pRS413-*chrXI*_tRNA (*HIS3*) | *synXI* intermediate strain with megachunks A-M (3.37) | This study |
| ysXIa23 | *MAT**a** his3*Δ1 *leu2*Δ0 *met15*Δ0 *ura3*Δ0 *YKR034W*::*URA3* pRS413-*chrXI*_tRNA (*HIS3*) | *synXI* intermediate strain with megachunks A-N | This study |
| ysXIa24 | *MAT**a** his3*Δ1 *leu2*Δ0 *met15*Δ0 *ura3*Δ0 *YKR055W*:: *LEU2* pRS413-*chrXI*_tRNA (*HIS3*) | *synXI* intermediate strain with megachunks A-O (partial) | This study |
| ysXIa25 | *MAT**a** his3*Δ1 *leu2*Δ0 *met15*Δ0 *ura3*Δ0 *YKR055W*:: *LEU2* pRS413-*chrXI*_tRNA (*HIS3*) | *synXI* intermediate strain with megachunks A-O | This study |
| ysXIa26 | *MATα his3*Δ1 *leu2*Δ0 *lys2*Δ0 *ura3*Δ0 *YKR055W*::*LEU2* | *synXI* intermediate strain with chunk O5 | This study |

| Strain | Genotype | Description | Source |
|---|---|---|---|
| ysXIa27 | *MATα his3*Δ1 *leu2*Δ0 *lys2*Δ0 *ura3*Δ0 *YKR077W*::*URA3* | *synXI* intermediate strain with chunks O5-P5 | This study |
| ysXIa28 | *MATα his3*Δ1 *leu2*Δ0 *lys2*Δ0 *ura3*Δ0 $\rho^0$ *YKR095W*::*LEU2* | *synXI* intermediate strain with chunks O5-Q5 | This study |
| ysXIa29 | *MATα his3*Δ1 *leu2*Δ0 *lys2*Δ0 *ura3*Δ0 $\rho^0$ *URA3* | *synXI* intermediate strain with chunks O5-R5 | This study |
| ysXIa30 | *MATα his3*Δ1 *leu2*Δ0 *lys2*Δ0 *ura3*Δ0 $\rho^0$ | *synXI* intermediate strain with chunks O5-R5. *URA3* marker removed | This study |
| ysXIa31 | *MATα his3*Δ1 *leu2*Δ0 *lys2*Δ0 *ura3*Δ0 $\rho^0$ *klURA3-PGAL1-CEN11* | *synXI* intermediate strain with chunks O5-R5. *CEN11* replaced with *CEN11** | This study |
| ysXIa32 | *MAT**a**/α his3*Δ1/*his3*Δ1 *leu2*Δ0/*leu2*Δ0 *LYS2*/*lys2*Δ0 *met15*Δ0/*MET15 ura3*Δ0/*ura3*Δ0 *YKR055W*/*YKR055W*::*LEU2 klURA3-PGAL1-CEN11*/*CEN11* pRS413-*chrXI*_tRNA (*HIS3*) | Diploid of strains ysXIa25 and ysXIa31 | This study |
| ysXIa33 | *MAT**a**/α his3*Δ1/*his3*Δ1 *leu2*Δ0/*leu2*Δ0 *LYS2*/*lys2*Δ0 *met15*Δ0/*MET15 ura3*Δ0/*ura3*Δ0 *synXI*_9.01 *klURA3-PGAL1-CEN11*/*CEN11* pRS413-*chrXI*_tRNA (*HIS3*) | ysXIa33 after induced crossover | This study |
| ysXIa34 | *MAT**a**/α his3*Δ1/*his3*Δ1 *leu2*Δ0/*leu2*Δ0 *LYS2*/*lys2*Δ0 *met15*Δ0/*MET15 ura3*Δ0/*ura3*Δ0 *synXI*_9.01 pRS413-*chrXI*_tRNA (*HIS3*) | ysXIa34 after disruption of *CEN11** | This study |
| ysXIb01 | *MAT**a** his3*Δ1 *leu2*Δ0 *lys2*Δ0 *ura3*Δ0 $\rho^0$ *synXI*_9.01 pRS413-*chrXI*_tRNA (*HIS3*) | Haploid strain with *synXI*_9.01 | This study |
| ysXIb02 | *MAT**a** his3*Δ1 *leu2*Δ0 *lys2*Δ0 *ura3*Δ0 $\rho^0$ *synXI*_9.02 *YKL069W*::*LEU2 YKL053C*::*kanMX* pRS413-*chrXI*_tRNA (*HIS3*) | Markers inserted flanking megachunk J repeats | This study |
| ysXIb03 | *MAT**a** his3*Δ1 *leu2*Δ0 *lys2*Δ0 *ura3*Δ0 $\rho^0$ *synXI*_9.03 *YKL048C*::*URA3* pRS413-*chrXI*_tRNA (*HIS3*) | Megachunk J integrated to reduce megachunk J copy number | This study |
| ysXIb04 | *MAT**a** his3*Δ1 *leu2*Δ0 *lys2*Δ0 *ura3*Δ0 $\rho^0$ *synXI*_9.04 pRS413-*chrXI*_tRNA (*HIS3*) | *URA3* marker removed | This study |
| ysXIb05 | *MAT**a** his3*Δ1 *leu2*Δ0 *lys2*Δ0 *ura3*Δ0 $\rho^0$ *synXI*_9.05 pRS413-*chrXI*_tRNA (*HIS3*) | Megachunk J repeats condensed to single copy | This study |
| ysXIb06 | *MAT**a** his3*Δ1 *leu2*Δ0 *lys2*Δ0 *ura3*Δ0 $\rho^0$ *synXI*_9.06 *YKR077W*::*URA3* pRS413-*chrXI*_tRNA (*HIS3*) | *URA3* inserted upstream of megachunk Q repeats | This study |
| ysXIb07 | *MAT**a** his3*Δ1 *leu2*Δ0 *lys2*Δ0 *ura3*Δ0 $\rho^0$ *synXI*_9.07 pRS413-*chrXI*_tRNA (*HIS3*) | Megachunk Q integrated to reduce megachunk Q copy number | This study |
| ysXIb08 | *MAT**a** his3*Δ1 *leu2*Δ0 *lys2*Δ0 *ura3*Δ0 $\rho^0$ *synXI*_9.08 pRS413-*chrXI*_tRNA (*HIS3*) | Megachunk Q repeats condensed to single copy | This study |
| ysXIb09 | *MAT**a** his3*Δ1 *leu2*Δ0 *lys2*Δ0 *ura3*Δ0 $\rho^0$ *synXI* _9.09 pRS413-*chrXI*_tRNA (*HIS3*) | Sequence insertions at *YKR050W* removed | This study |
| ysXIb10 | *MAT**a** his3*Δ1 *leu2*Δ0 *lys2*Δ0 *ura3*Δ0 $\rho^0$ *synXI* _9.10 pRS413-*chrXI*_tRNA (*HIS3*) | *YKR084C-YKR087C* locus replaced | This study |
| BY4742-*CEN11** | *MATα his3*Δ1 *leu2*Δ0 *lys2*Δ0 *ura3*Δ0 *klURA3-PGAL1-CEN11* | BY4742 with *CEN11* replaced with *CEN11** | This study |
| ysXIb11 | *MAT**a**/α his3*Δ1/*his3*Δ1 *leu2*Δ0/*leu2*Δ0 | Diploid of ysXIb10 and BY4742-*CEN11** | This study |

| Strain | Genotype | Description | Source |
|---|---|---|---|
| | *lys2*Δ0/*lys2*Δ0 *ura3*Δ0/*ura3*Δ0 *synXI*_9.10 pRS413-*chrXI*_tRNA (*HIS3*) | after disruption of *CEN11** | |
| ysXIb12 | *MAT***a**/*MAT***a** *his3*Δ1/*his3*Δ1 *leu2*Δ0/*leu2*Δ0 *lys2*Δ0/*lys2*Δ0 *ura3*Δ0/*ura3*Δ0 *synXI*_9.10 pRS413-*chrXI*_tRNA (*HIS3*) | Presumed haploid of ysXIb11 - later found to be homozygous diploid | This study |
| ysXIb13 | *MAT***a**/*MAT***a** *his3*Δ1/*his3*Δ1 *leu2*Δ0/*leu2*Δ0 *lys2*Δ0/*lys2*Δ0 *ura3*Δ0/*ura3*Δ0 *synXI*_9.11 pRS413-*chrXI*_tRNA (*HIS3*) | *YKR086W* TAG stop codon converted to TAA | This study |
| ysXIb14 | *MAT***a**/*MAT*α *his3*Δ1/*his3*Δ1 *leu2*Δ0/*leu2*Δ0 *lys2*Δ0/*lys2*Δ0 *ura3*Δ0/*ura3*Δ0 *synXI*_9.11 pRS413-*chrXI*_tRNA (*HIS3*) | Presumed diploid of ysXIb13 and BY4742-*CEN11** after disruption of *CEN11** - later deduced to be polyploid | This study |
| ysXIb16 | *MAT***a**/*MAT***a** *his3*Δ1/*his3*Δ1 *leu2*Δ0/*leu2*Δ0 *lys2*Δ0/*lys2*Δ0 *ura3*Δ0/*ura3*Δ0 *synXI*_9.11 pRS413-*chrXI*_tRNA (*HIS3*) | Presumed haploid of ysXIb16 - later found to be homzygous diploid | This study |
| ysXIb17 | *MAT***a** *his3*Δ1 *leu2*Δ0 *lys2*Δ0 *ura3*Δ0 *synXI*_9.11 pRS413-*chrXI*_tRNA (*HIS3*) | Haploid of ysXIb16. Final synXI strain | This study |
| y*CEN11*d1 | *MAT*α *his3*Δ1 *leu2*Δ0 *lys2*Δ0 *ura3*Δ0 *CEN11*_3_35 | BY4742 with synthetic *CEN11*_3_35 | This study |
| y*CEN11*d2 | *MAT*α *his3*Δ1 *leu2*Δ0 *lys2*Δ0 *ura3*Δ0 *CEN11*_3_37 | BY4742 with synthetic *CEN11*_3_37 | This study |
| y*CEN11*d3 | *MAT*α *his3*Δ1 *leu2*Δ0 *lys2*Δ0 *ura3*Δ0 *CEN11*_3_37b | BY4742 with synthetic *CEN11*_3_37b | This study |
| y*CEN11*d4 | *MAT*α *his3*Δ1 *leu2*Δ0 *lys2*Δ0 *ura3*Δ0 *CEN11*_3_37c | BY4742 with synthetic *CEN11*_3_37c | This study |
| y*CEN11*d5 | *MAT*α *his3*Δ1 *leu2*Δ0 *lys2*Δ0 *ura3*Δ0 *CEN11*_3_37f | BY4742 with synthetic *CEN11*_3_35f | This study |
| $GAP1_{syn}$ | *MAT***a** *his3*Δ1 *leu2*Δ0 *met15*Δ0 *ura3*Δ0 $GAP1_{syn}$ | BY4741 with synthetic *GAP1* locus | This study |
| $GAP1_{WT}$-*yEGFP* | *MAT***a** *his3*Δ1 *leu2*Δ0 *met15*Δ0 *ura3*Δ0 *yEGFP* | BY4741 with *yEGFP* integration upstream of *GAP1* | This study |
| $GAP1_{syn}$-*yEGFP* | *MAT***a** *his3*Δ1 *leu2*Δ0 *met15*Δ0 *ura3*Δ0 $GAP1_{syn}$ *yEGFP* | $GAP1_{syn}$ with *yEGFP* integration upstream of *GAP1* | This study |
| $GAP1_{WT}$-*yEGFP*-*BFP2* | *MAT***a** *his3*Δ1 *leu2*Δ0 *met15*Δ0 *ura3*Δ0 *yEGFP* *LEU2*::*BFP2* | $GAP1_{WT}$-*yEGFP* with aTc inducible *BFP2* integrated at the *LEU2* locus | This study |
| $GAP1_{WT}$-*yEGFP*-*mScarlet* | *MAT***a** *his3*Δ1 *leu2*Δ0 *met15*Δ0 *ura3*Δ0 *yEGFP* *LEU2*::*mScarle*t | $GAP1_{WT}$-*yEGFP* with aTc inducible *mScarlet* integrated at the *LEU2* locus | This study |
| $GAP1_{syn}$-*yEGFP*-*BFP2* | *MAT***a** *his3*Δ1 *leu2*Δ0 *met15*Δ0 *ura3*Δ0 $GAP1_{syn}$ *yEGFP* *LEU2*::*BFP2* | $GAP1_{syn}$-*yEGFP* with aTc inducible *BFP2* integrated at the *LEU2* locus | This study |
| $GAP1_{syn}$-*yEGFP*-*mScarle*t | *MAT***a** *his3*Δ1 *leu2*Δ0 *met15*Δ0 *ura3*Δ0 $GAP1_{syn}$ *yEGFP* *LEU2*::*mScarle*t | $GAP1_{syn}$-*yEGFP* with aTc inducible *mScarlet* integrated at the *LEU2* locus | This study |

***Table S1 - List of yeast strains used in this study.***

| Name | Sequence (5’ - 3') | Target | Details |
|---|---|---|---|
| BB_mat _a | ACTCCACTTCAAGTAAGAGTTTG | *MATA1* | Mating type determination. Sequence taken from Huxley *et al*., 1990 |
| BB_mat _alpha | GCACGGAATATGGGACTACTTCG | *MATALPHA1* | Mating type determination. Sequence taken from Huxley *et al*., 1990 |
| BB_mat _uni | AGTCACATCAAGATCGTTTATGG | *YCR041W* | Mating type determination. Sequence taken from Huxley *et al*., 1990 |
| BB210 | GTACGAATTCCTCACTACTATGACTTCCATAGCTCACC | BY4741, *TRT2* 5' | Amplification of *TRT2* for pRS403::*TRT2* generation |
| BB211 | GTACCCCGGGGCGAAGCAGCGTGTAAATTTCCCTATTAGG | BY4741, *TRT2* 3' | Amplification of *TRT2* for pRS403::*TRT2* generation |
| BB286 | TGACCGGAAAAGACAGTTGT | Chunk M1 | Editing M1 to conform to *synXI*_3.36 |
| BB287 | CATTCTATATACCCTGTTTAAGTTATTTTAATGAGCCAATCA AGCCAAAAATAAATAAT | Chunk M1 | Editing M1 to conform to *synXI*_3.36 |
| BB288 | ATTATTTATTTTTGGCTTGATTGGCTCATTAAAATAACTTAA ACAGGGTATATAGAATG | Chunk M1 | Editing M1 to conform to *synXI*_3.36 |
| BB289 | GGGAGAACATCTTGAGGCAATG | Chunk M1 | Editing M1 to conform to *synXI*_3.36 |
| BB290 | CAAGTTGCAGATCGTTGTTGA | Chunk M3 | Editing M3 to conform to *synXI*_3.36 |
| BB291 | GAGCGCAATGGTTTCATGATTATTCTACACCAAGTTGTTGC GCCATTTTATCTAAGCTGA | Chunk M3 | Editing M3 to conform to *synX*I_3.36 |
| BB292 | TCAGCTTAGATAAAATGGCGCAACAACTTGGTGTAGAATAA TCATGAAACCATTGCGCTC | Chunk M3 | Editing M3 to conform to *synXI*_3.36 |
| BB293 | ACGGAACATTAAAAGAGCTTGTT | Chunk M3 | Editing M3 to conform to *synXI*_3.36 |
| BB294 | TCTGATCAAAGCGGCATTTCT | Chunk M3 | Editing M3 to conform to *synXI*_3.36 |
| BB295 | TTATATGTTTTTCTTTTTTTTGTTTTAACTATCTGATTGTTCT TTCTTATTTTTATCC | Chunk M3 | Editing M3 to conform to *synXI*_3.36 |
| BB296 | GGATAAAAATAAGAAAGAACAATCAGATAGTTAAAACAAAA AAAAGAAAAACATATAA | Chunk M3 | Editing M3 to conform to *synXI*_3.36 |
| BB297 | GGTAATTCAAGTGCGGCCAT | Chunk M3 | Editing M3 to conform to *synXI*_3.36 |
| BB353 | /5PHOS/GTTTTAGAGCTAGAAATAGCAAGTTAAAATAAGGC TAG | pWS082 | Combined with target specific primer for CRISPR/Cas9 gRNA retargeting |
| BB367 | GGTGATGAGCCAAATGTCGTAGTGG | Chunk M2 | Editing M2 to conform to *synXI*_3.37 |
| BB368 | ACCGGTCGACTCAAGCTTGGTT | Chunk M2 | Editing M2 to conform to *synX*I_3.37 |
| BB369 | CCTAAACACTAGTATTGACAAAAGAATAATATTATATGC | Chunk M2 | Editing M2 to conform to *synXI*_3.37 |
| BB370 | /5Phos/ATAACTTCGTATAATGTACATTATACGAAGTTATTTC CTCATCTATACCGGTCGAC | Chunk M2 | Editing M2 to conform to *synXI*_3.37 |
| BB371 | GAAGACAATAAAGAGGAATGGGGTG | Chunk M2 | Editing M2 to conform to *synXI*_3.37 |
| BB372 | AACTTAAACAGAGGAGACGATTTGAC | Chunk M2 | Editing M2 to conform to *synXI*_3.37 |
| BB373 | CCTAATACCTCAATGGTCCAATACTAAATAAGGTACTATTC ATTGTATTGATTGATTCTG | Chunk M2 | Editing M2 to conform to *synXI*_3.37 |
| BB374 | ACCTTATTTAGTATTGGACCATTGAGGTATTAGGTATTAGT AGAAATATCCTAAACACTA | Chunk M2 | Editing M2 to conform to *synXI*_3.37 |
| BB418 | GGAGTCACTGCCAGGTATCGTT | BY4741, *his3*Δ1 5' | CRISPR/Cas9 removal of *TRT2* from *HIS3* locus repair template |
| BB419 | GTTGTAGCCGCCGTTGTTGTTA | BY4741, | CRISPR/Cas9 removal of *TRT2* from |

| Name | Sequence (5’ - 3') | Target | Details |
|---|---|---|---|
| | | *his3*Δ1 3' | *HIS3* locus repair template |
| BB420 | TTTTTACTCCACGCGCCAGT | *HIS3* region deleted in BY4741 | *TRT* removal screening |
| BB421 | ATCAACAACAGAGGACATATGCCCTACC | pWS082 | Amplification of retargeted gRNA piece for CRISPR/Cas9 |
| BB422 | ATCCTGCACTCATCTACTACCCGCA | pWS082 | Amplification of retargeted gRNA piece for CRISPR/Cas9 |
| BB433 | TCAGGGAATTCGGTCTCTCCTCTTCTCCATTGCCACA | BY4741, *CEN11* upstream | Generating *CEN11** |
| BB434 | TACCGCTGATAGGCGATTCCCGAAGTTTAAATTACAGAAGT TAGGTGAGGTTAATTACCA | BY4741, *CEN11* upstream | Generating *CEN11** |
| BB435 | AAGAAATTGGTAATTAACCTCACCTAACTTCTGTAATTTAAA CTTCGGGAATCGCCTATC | pJJH1304, *klURA3* 5' | Generating *CEN11** |
| BB436 | GTTCCAAAGAGAAGGTTTTTTTAGGCTAAGATAATGGGGG GATCTGTTTAGCTTGCCTCG | pJJH1304, *klURA3* 3' | Generating *CEN11** |
| BB437 | CGGCGGGGACGAGGCAAGCTAAACAGATCCCCCCATTAT CTTAGCCTAAAAAAACCTTCT | BY4741, *PGAL1* 5' | Generating *CEN11** |
| BB438 | GCAAAATTGTAAATAGCGTTGACAAGTGAAGATCTTATAGT TTTTTCTCCTTGACGTTAA | BY4741, *PGAL1* 3' | Generating *CEN11** |
| BB439 | TTTAACGTCAAGGAGAAAAAACTATAAGATCTTCACTTGTC AACGCTATTTACAATTTTG | *CEN11* amplicon | Generating *CEN11** |
| BB440 | AGCAAGCTTGGTCTCTATCTTTTGTCCAATATGAAGAAGGT CAAC | *CEN11* amplicon | Generating *CEN11** |
| BB458 | GTTTGGTTGTCTTGCCAGATGAG | R5 *URA3* upstream | R5 *URA3* loss screening |
| BB459 | CATACATTATACGAAGTTATCGGTCCG | R5 *URA3* downstream | R5 *URA3* loss screening |
| BB468 | GCGCTTGCTCTAAACATGCATAGGG | *CEN11* | Generating *CEN11* variants |
| BB469 | CATCTATACCGGTCGACTCAAGCTTGG | *CEN11* | Generating *CEN11* variants |
| BB472 | CCTAATACCTCAATGGTCCAATACTAAATAAGGTACTATTC ATTGTATTGATTGATTCTG | *CEN11* | Generating *CEN11* variants |
| BB473 | /5Phos/ATAACTTCGTATAATGTACATTATACGAAGTTATTAT TAGTAGAAATATCCTAAACACTA | *CEN11* | Generating *CEN11* variants |
| BB475 | /5Phos/ATAACTTCGTATAATGTACATTATACGAAGTTATTGT AAATAGCGTTGACAAGTGATTAC | *CEN11* | Generating *CEN11* variants |
| BB476 | CTCTCCTTTCTATGTGTATTTGATTATTTTAATTACCAAG | *CEN11* | Generating *CEN11* variants |
| BB479 | /5Phos/TATTAGTAGAAATATCCTAAACACTAGTATTGACAAA AG | *CEN11* | Generating *CEN11* variants |
| BB481 | TCACTAAATAATATAGTTATACATCTAGCTAATCACTATTCA TTGTATTGATTGATTCTG | *CEN11* | Generating *CEN11* variants |
| BB570 | CGTATCTAGACTGTTTAGCTTGCCTCGTC | Y07039, *kanMX4* 5' | Generating pSXI_3_34_J4::*kanMX* |
| BB571 | CGTAGGCGCCGTTTTCGACACTGGATGG | Y07039, *kanMX4* 3' | Generating pSXI_3_34_J4::*kanMX* |
| BB582 | CTTTTGATGTCAGCTTCG | *GAP1* locus | Amplification of *GAP1* locus |

| Name | Sequence (5’ - 3') | Target | Details |
|---|---|---|---|
| BB583 | AAGAGTCTACTAAGCAAAACATGATG | *GAP1* locus | Screening of *GAP1* locus |
| BB584 | AGGAAATCCCCTCTTATGATCT | *GAP1* locus | Screening of *GAP1* locus |
| BB585 | TCAAGCGGTCTCTTTCGC | *GAP1* locus | Amplification of *GAP1* locus |
| BB683 | TGGTGTTATATACTGTTACTGGAGCTC | YKR087C | *OMA1* loxP screening |
| BB738 | TTCCGGTCATTTCCTCCGGT | *PRP16* 3' region | CRIPSR/Cas9 recoding of *PRP16* stop codon repair template |
| BB768 | GCAATAATCAGATTGTGAATTCC | *YKR084C* | CRISPR/Cas9 replacement of *HBS1-OMA1* locus repair template |
| BB775 | ATGTTACGCAACATCATCAGGT | *YKR087C* | CRISPR/Cas9 replacement of *HBS1-OMA1* locus repair template |
| BB788 | GAGTTATGCTCTTTTCTTTTGTACCC | *PRP16* 3' region | CRIPSR/Cas9 recoding of *PRP16* stop codon repair template |
| BB789 | CACGCTCTTCCCATAAAT | *PRP16* TAA stop codon | *PRP16* stop codon screening |
| BB790 | CATAACTCGTGGAGATAACTTCG | *YKR087C* (loxPsym present) | *OMA1* loxPsym screening |
| BB791 | CATAACTCGTGGAGTGCTTACAT | *YKR087C* (loxPsym absent) | *OMA1* loxP screening |
| BB877 | CGACATTAACGGATACGCAACTG | *CEN11* variants 5' | *CEN11** CRISPR/Cas9 replacement repair template |
| BB878 | CAGGAACTTTCTGGTGGTGCTAG | *CEN11* variants 3' | *CEN11** CRISPR/Cas9 replacement repair template |
| oLM394 | GTCAAGCCAATAATGGTTTAGGTAGTAGG | *YNCQ0002W* | Mitochondrial genome screening |
| oLM395 | CACGGGTATCGAATCCGTTTCGC | *YNCQ0002W* | Mitochondrial genome screening |
| oLM396 | CTAGCTTAAGCAGCTTTTCAATTCAATTCA | *URA3* | Amplification of *URA3* for marker swapper construct |
| oLM397 | CTAGCTTAAGATAGGGTAATAACTGATATAATTAAATTG | *URA3* | Amplification of *URA3* for marker swapper construct |
| oLM398 | CAGGATTCAGCAACACCAAATCAAGAA | *Q0250* | Mitochondrial genome screening |
| oLM399 | CCCACACAACTCAGAACATGCTC | *Q0250* | Mitochondrial genome screening |
| XL217 | GGATAGTGTTCAAGTTGACCAT | *YKRCδ11* upstream | Screening *GAP1* locus rearrangements |
| XL492 | CATCATGTTTTGCTTAGTAGACTC | Region 5' of *GAP yEGFP* insertion | Amplification of 5' homology arm for *yEGFP* insertion |
| XL493 | CGCCAAAAATCATGGTAAAACCTCCCACACAGAATCGAGG TATTTTTTGATTACTTGACC | Region 5' of *GAP yEGFP* insertion | Amplification of 5' homology arm for *yEGFP* insertion |
| XL494 | TACACCTCGCTCTGGGTCAAGTAATCAAAAAATACCTCGAT TCTGTGTGGGAGGTTTTAC | *PPFY1* | Amplifying *yEGFP* expression cassette |
| XL495 | GTCAAACATAAACCAAGCGACAGATTTGTCGAAGATATTCG ATCTTCGAGCGTCCCAAA | *TCYC1* | Amplifying *yEGFP* expression cassette |
| XL496 | GAAAACCTTGCTTGAGAAGGTTTTGGGACGCTCGAAGATC GAATATCTTCGACAAATCTG | Region 3' of *GAP yEGFP* insertion | Amplification of 3' homology arm for *yEGFP* insertion |

| Name | Sequence (5' - 3') | Target | Details |
|---|---|---|---|
| XL497 | GAGAAAGTTTCTTTTAACTTTTGGC | Region 3' of *GAP yEGFP* insertion | Amplification of 3' homology arm for *yEGFP* insertion |
| XL788 | CCTACTGGGGGTTATTTATGG | *GAP1* | Screening *GAP1* locus rearrangements |
| XL789 | TTCATATGATCTGGGTATCTAGC | *yEGFP* | Screening *GAP1* locus rearrangements |
| XL790 | CACTCCGATAACTAAACTTTGC | *ARS1116* | Screening *GAP1* locus rearrangements |
| XL808 | AACGTAGAATTGGGCAATG | *GAP1* | Screening *GAP1* locus rearrangements |
| XL809 | CCGCAGTGAAAGTTAATAACG | *ARS1116* downstream | Screening *GAP1* locus rearrangements |

***Table S2: List of primers used in this study. 5PHOS denotes that the primer is 5' phosphorylated. PCRTag primers are listed in Table S3 and CRISPR/Cas9 gRNA retargeting primers are listed in Table S4.***

| Amplicon | WT_F | WT_R | syn_F | syn_R | bp | chunk |
|---|---|---|---|---|---|---|
| *YKL222C 2* | TCGGACCGGAATTC CTGTAGAGAGTGAA | CCATCCAAAAGAAG TGCCAGAGGCTATT | TCTAACTGGGATAC CGGTGCTCAAGCTG | ACACCCTAAGGAAG TTCCTGAGGCCATC | 340 | A1 |
| *YKL222C 1* | AACACTTGGTGAGC CTGAGTGCGGT | CTGTGGAAGTTGCT CTTCTCGAAATCGA | GACTGAAGGGCTAC CGCTATGTGGA | TTGCGGTTCATGTA GCAGCAGAAACAGA | 421 | A1 |
| *YKL221W 1* | ATGCACATCACTTA GCACCATTGCGCTT | AGGTATCCACATGG CCCAACAAAGGATG | CTGTACCAGCTTGT CAACTATCGCTTTG | TGGGATCCACATAG CCCAGCACAAAATA | 373 | A1 |
| *YKL220C 1* | CACAGATAATGAGC CACTCAATTCAGCC | ATTGTTACTCACTGG AGGTACCGGTTTG | AACGCTCAAGCTAC CTGATAATTCGGCA | TTTATTGTTGACCGG TGGCACTGGCTTA | 403 | A2 |
| *YKL220C 2* | GTGTGCGAATGCGA GTACACCACTTCTG | CATGGCTTTTGCAG GCGTTCTTCATTGC | ATGAGCAAAAGCCA AAACGCCTGAACGA | TATGGCCTTCGCTG GTGTCTTGCACTGT | 331 | A2 |
| *YKL219W 1* | ATCATTTGGGGATT CTGTCCCCTACATC | AAAACCTCCCCCAC AACACAAGATACGA | GAGCTTCGGTGACA GCGTTCCATATATT | GAAGCCACCACCGC AGCATAAAATTCTG | 283 | A2/ B1 |
| *YKL218C 1* | TTTGCAGCCTGGAG AAAGGCTTCTAGCG | CAGGTATACTGAAG ATCGCGAGCAGATT | CTTACAACCAGGGC TCAATGAACGGGCA | TAGATACACCGAAG ACAGAGAGCAGATC | 229 | B1 |
| *YKL217W 2* | CCATAACCCATCAC TGCCTGCACAATCA | CAATGGAGCGACTG ATACTGAAACGCAA | TCACAATCCTAGCTT ACCAGCTCAAAGC | TAAAGGGGCAACGC TAACGCTGACACAG | 328 | B1/ B2 |
| *YKL217W 1* | GAAAGCACGTTCGT TCCTATCAGGTCTA | AACCACTACAATGAC AGTGACAGCATCC | TAAGGCTAGAAGCT TTTTGAGCGGCTTG | GACAACAACGATAA CGGTAACGGCGTCT | 454 | B2 |
| *YKL216W 1* | AGCTGGCGCATTCA TTACAAAGAGTGCT | GTCATAAGCAACTTG TGGTTTCCCAGGC | GGCCGGTGCTTTTA TCACCAAATCAGCC | ATCGTAGGCGACTT GAGGCTTACCTGGA | 319 | B2 |

| Amplicon | WT_F | WT_R | syn_F | syn_R | bp | chunk |
|---|---|---|---|---|---|---|
| *YKL215C 1* | GCTCAAGAGGGAAC CAGCAGGAATTTTT | CGATTTAAAAGCGC AAGTCGCAGCTAAC | TGATAACAAACTGCC GGCTGGGATCTTG | TGACTTGAAGGCTC AAGTTGCTGCCAAT | 430 | B2 |
| *YKL215C 2* | TCTTGAACCTGAGC AGCCTGGGTATTTG | AGGAAACTTAGTCG CTAACGCTCCTCAT | ACGGCTGCCGCTAC AACCAGGATACTTA | GGGTAATTTGGTTG CCAATGCCCCACAC | 406 | B2 |
| *YKL215C 4* | TCCGCTTTTAGATAA TGTCTTCGCAGCC | GAAAGCCGACGTGT CAAAGGATGCTAGT | ACCTGACTTGCTCAA GGTTTTAGCGGCT | AAAGGCTGATGTTA GCAAAGACGCCTCA | 244 | B2 |
| *YKL215C 3* | AGTAAGGTAGGCAT CTGCGACAGAACTG | TGGTGAACGTTGTG CCTTCATTACTACG | GGTCAAATAAGCGT CAGCAACGCTTGAA | CGGCGAAAGATGCG CTTTTATCACCACC | 481 | B2 |
| *YKL214C 1* | GTCAAGAGCTTCGA CAGAAGTAGGAGTG | ACCTAGACTGGGAT TTGCGCCTTCGGAT | ATCCAAGGCTTCAA CGCTGGTTGGGGTA | TCCACGTTTAGGTTT CGCTCCAAGCGAC | 478 | B3 |
| *YKL213C 1* | AGAGTTCCCTTTCG TGACGAGAGCTGAA | TCTGGCGCAAATTG GTGGCGCACTACAT | GCTATTACCCTTGGT AACCAAGGCGCTG | CTTAGCTCAAATCG GCGGTGCTTTGCAC | 412 | B3 |
| *YKL213C 2* | AACGGACCAAATAG AGATGGCAGGAAGT | CCAAGATGGTGTTG TGATAAGCGGTAGT | GACACTCCAGATGC TAATAGCTGGCAAG | TCAAGACGGCGTCG TTATCTCAGGCTCA | 481 | B3 |
| *YKL212W 2* | TCAAGGTGTTCGGG TCCTTGGAGCA | TCGTTCATCAGCGG TTTTCCAGGAGGCT | CCAAGGCGTCAGAG TTTTGGGTGCT | TCTTTCGTCGGCAG TCTTCCAACTAGCA | 358 | B3 |
| *YKL212W 1* | TAGACCACATACCG CTTCTATCAAGTCG | ATGCGTTTGATGAAC GACCAAGAACCCA | CCGTCCTCACACTG CCAGCATTAAAAGC | GTGGGTTTGGTGGA CAACTAAAAAACCG | 277 | B3 |
| *YKL211C 1 ** | GGCTCTTTTATGGG ATGATGACAACACC | TTCATTGGCCGGGA CAGAATCGTCCCTA | AGCACGCTTGTGAC TGCTGCTTAAAACG | CAGCTTAGCTGGTA CCGAAAGCAGTTTG | 382 | B3 |
| *YKL210W 2* | TAGATTCATTTCCTC GGAGACAAGAGGG | GACTGAACCAATTCT GAATGCAAAGGGC | CCGTTTTATCAGTAG CGAGACCCGTGGT | AACGCTGCCGATAC GAAAAGCGAATGGA | 283 | B4 |
| *YKL210W 1* | CTCTTCTTCTAGAGA CCCACCAGAAAAG | AAATGGTTCACCGTT GGAAGTCTTGGCA | TAGCAGCAGCCGTG ATCCTCCTGAAAAA | GAAAGGTTCGCCAT TACTGGTTTTAGCG | 364 | B4 |
| *YKL210W 3* | ATCCCTACCAGATC CTTCTACTTTGGCT | AGAAGTGGTTGTTG CGATGGCAGGAATA | TAGTTTGCCTGACC CAAGCACCTTAGCC | GCTGGTAGTGGTAG CAATAGCTGGGATG | 205 | B4 |
| *YKL209C 1* | GGAGCTGACAGAAT CCAAGGCTGAAGTA | TCCATCTGCACCTAC CGCCTTTGTTTAC | ACTTGAAACGCTGT CTAAAGCGCTGGTG | CCCTAGCGCTCCAA CTGCTTTCGTCTAT | 493 | B4 |
| *YKL209C 3* | AATGCTCAGAAGTT GTGGCCCTGTTGAA | TGTTGAAAGCGGCA CCCAGTCTGAACTT | GATTGATAACAATTG AGGACCGGTGCTG | CGTCGAATCAGGTA CTCAGAGCGAATTG | 358 | B4 |
| *YKL209C 4* | GGTGCTGCAATCGG CATTTCTTACCGAA | TTATCCAAGCAGAC CTTCGGAAGCAGTT | AGTTGAACAGTCAG CGTTACGAACGCTG | ATACCCTTCACGTCC AAGCGAAGCTGTC | 310 | B4 |
| *YKL209C 2* | TGCGGAGCTTGATC TTAGCTCTTCCACA | GGCAATTACTTCTAT CCTGACGGGCAGA | AGCACTTGAGCTAC GCAACTCTTCAACG | TGCTATCACCAGCAT TTTAACCGGTCGT | 325 | B5 |
| *YKL208W 1* | CGCTGGTAGTCTTG TCGATTCTTCCTAC | GAGAGATGGGATGT CCTGAACTGTCTGA | TGCCGGCTCATTGG TTGACAGCAGTTAT | CAAGCTAGGAATAT CCTGGACGGTCTGG | 235 | B5 |

| Amplicon | WT_F | WT_R | syn_F | syn_R | bp | chunk |
|---|---|---|---|---|---|---|
| *YKL207W 1* | CCTGTCTTCCGATG CATTTGCTGCGAAG | GCTAACCCATCTCA CATCCAGGTCCTGA | TTTAAGCAGTGACG CTTTCGCCGCTAAA | TGAGACCCAACGAA CGTCTAAATCCTGG | 331 | B5 |
| *YKL206C 1 ** | GAGGTTCGATAAAA GCTCAGCTTCATCG | ATTAGACAGACCAG AGGATGGTAGCGAC | CAAATTGCTCAACAA CTCGGCTTCGTCA | TTTGGATCGTCCTGA GGACGGCTCAGAT | 325 | B5 |
| *YKL205W 1* | GGATTCGGCTATTT CCAATACGAGCCCG | TCCGGGACATTGTG ACACAATTTGTTCG | TGACAGCGCCATCA GTAACACCTCACCA | ACCTGGGCATTGGC TAACGATTTGTTCA | 250 | B5 |
| *YKL205W 3* | AGTAACATCTATCAA CGCCCAAGGGACG | GAGACCAATGTGCA AACCTCTCGCTAAA | GGTTACCAGCATTAA TGCTCAAGGTACC | CAAGCCGATATGTA AGCCACGAGCCAAG | 295 | B5/ C1 |
| *YKL205W 2* | CCGCGATGCATCTA GGTTTACCTTTGCC | GGCTGTGACTGCAG TGTCTATAGGTTTG | TAGAGACGCTAGCA GATTCACTTTCGCT | AGCGGTAACAGCGG TATCGATTGGCTTA | 310 | C1 |
| *YKL204W 1* | CGGTAGCGTAGAAT CTTCAACTGATAGC | AGCCTTCCAACTGC CTCCACCATTG | TGGCTCAGTTGAAA GCAGCACCGACTCA | GGCTTTCCATGAAC CACCGCCGTTA | 367 | C1 |
| *YKL204W 2* | TGGACAGCCAAGCC TATCCAAAACCAGT | AATTAGTCCTGGGG GTGGAGGAGGA | CGGTCAGCCTTCAT TGAGTAAGACTTCA | GATCAAACCAGGTG GAGGTGGTGGT | 460 | C1 |
| *YKL203C 2* | ACCCAGTAACCCAG ACTTCGGAGATAAT | GACCCGTGCAAGTG GTGGAAAACCAATT | GCCTAACAAACCGC TTTTTGGGCTCAAA | TACTAGAGCTTCAG GCGGTAAGCCTATC | 292 | C1 |
| *YKL203C 1* | ATCATGCGCAGATA GTAGTTTTGGCGAC | CGAATCTCTCTCAC GACAGAAAGCAGCT | GTCGTGAGCGCTCA ACAACTTAGGGCTA | AGAAAGCTTGAGCA GACAGAAGGCTGCC | 460 | C1 |
| *YKL203C 3* | CGAGGAAGCATCAC TTCCTTCCTGTTTC | CAAAAGAGTCCCTC GTCACGTTGAAGAT | GCTACTGGCGTCTG AACCTTCCTGCTTT | TAAGCGTGTTCCAA GACATGTCGAAGAC | 262 | C1 |
| *YKL203C 7* | CCTACCCGATTTTCT GCATAAGTTGGCA | CTGGGGTTTGGAGC AATGGGATGAAATA | TCTGCCGCTCTTAC GACACAAATTAGCG | TTGGGGCTTAGAGC AATGGGACGAAATC | 445 | C1 |
| *YKL203C 4* | ATCTGGAGACTGCG ATTTCATGACGCTA | AGCATATAATGAGAA GGAGGCAGCAGGA | GTCAGGGCTCTGGC TCTTCATAACTGAG | TGCTTACAACGAGA AAGAGGCTGCTGGT | 244 | C1 |
| *YKL203C 5* | AGCAGCCGTGTGAT GGATTGACAACGAT | TGGCCCTTTACTAGA CTACCCAGAGTTA | GGCGGCGGTATGGT GAATGCTTAAGCTA | CGGTCCATTGTTGG ATTATCCTGAGTTG | 316 | C2 |
| *YKL203C 6* | CTTTAAAGCGGTGG ATGCTACGGCAGAT | CAACCCCGCTTATG TAGTTCCTTCTTTG | TTTCAAGGCAGTACT AGCAACAGCGCTA | TAATCCAGCCTACGT TGTCCCAAGCTTA | 226 | C2 |
| *YKL201C 3* | CTTTGCAGGGGAGC TTAACTTAGGAGCA | GCCAGCGTTGATTC CTCACAGACATACC | TTTAGCTGGACTTGA CAATTTTGGGGCG | TCCTGCTTTAATCCC ACATCGTCACACT | 499 | C2 |
| *YKL201C 1* | GTAATCCCTACTAG GGGCAGAAGTGCTA | TGCCGTGAAGCAAT ATAAAGGTATGGGG | ATAGTCTCTTGATGG AGCGCTGGTTGAG | CGCTGTTAAACAATA CAAGGGCATGGGT | 463 | C2 |
| *YKL201C 2* | GATGGAAATGGATA GTGGATGCGACTGT | TGAGACAGAGCTCG GCGAACCTTTGTTC | AATACTGATACTCAA AGGGTGGCTCTGG | CGAGACCGAGTTGG GTGAACCATTATTT | 289 | C3 |
| *YKL201C 4* | TAAAGACTCTTTGTC GAAAGCGGACTGG | AAATTCACCGCAGC GTAACGGACTTGGG | CAAGCTCTCCTTATC AAAGGCACTCTGA | TAACAGCCCACAGA GAAATGGTTTGGGT | 475 | C3 |

| Amplicon | WT_F | WT_R | syn_F | syn_R | bp | chunk |
|---|---|---|---|---|---|---|
| *YKL198C 2* | CGGAGGCGACGAC GCTAAAACC | CCCATGGTTCCAAG GAATTGAGACTTGT | TGGTGGGCTGCTAG CCAAGACG | TCCTTGGTTTCAAGG TATCGAGACCTGC | 238 | C3 |
| *YKL198C 1* | CGAAGTGGACGAAT GTCTGTTTCCTCTT | GTCTCTACCTCCAG ACTTCCACTCTACA | GCTGGTACTGCTGT GACGATTACCACGA | TAGCTTGCCACCTG ATTTTCATAGCACC | 214 | C3 |
| *YKL197C 2* | TCTTCCCGGTCTTA ACAATGCGCTATCA | TGAAAGGGCACAGT CCGTCAAACCCTGT | ACGACCTGGACGCA ATAAAGCTGAGTCG | CGAAAGAGCTCAGA GTGTTAAGCCATGC | 217 | C3 |
| *YKL197C 1 ** | CGCAGATGGTGTAA ACGCACTAAGCGAC | TGGAAAACCTCAAG CCAATGACGGAGAT | AGCGCTAGGGGTGA AAGCTGACAAGCTT | CGGTAAGCCACAAG CTAACGATGGTGAC | 454 | C3 |
| *YKL197C 3 ** | ATCTTTACTAGTGC GCGAAACTTCGCCA | TCCTACCAAGAGTG GGGATCAGTATTCT | GTCCTTTGAGGTTCT GCTGACTTCACCG | CCCAACTAAATCAG GTGACCAGTACAGC | 271 | C3 |
| *YKL196C 1* | GGAACTTGCCGTTA ATGACTCCGATTTG | CGCACATCCTAAGG AAGAGTGGGCAGAT | ACTTGAAGCGGTCA AGCTCTCGCTCTTA | TGCTCACCCAAAAG AAGAGTGGGCTGAC | 241 | C4 |
| *YKL195W 1* | TAAAACCGCAGAGG AAGAGCTTTCCAGT | CTTGTCATCAGATTG TCCCTGTTGCGAT | CAAGACTGCTGAGG AAGAGTTGAGTTCA | TTTATCGTCGCTTTG ACCCTGTTGGCTC | 442 | C4 |
| *YKL194C 1* | CCTTAGTCTAGTCA ATCCCGACAGTGCA | AGATCCACTATCTCC TGGATCCATGTTC | TCTCAAACGGGTTAA ACCGCTTAAAGCG | CGACCCTTTGAGCC CAGGTAGTATGTTT | 376 | C4 |
| *YKL191W 2* | CATTCTCGGTTGTA GCCAAAGCGGTATC | ATCAAACTCCGGTTC ATCGGAAGTTGTG | TATCTTGGGCTGCT CACAATCAGGCATT | GTCGAACTCTGGTT CGTCACTGGTGGTA | 223 | C5 |
| *YKL191W 1* | TAGCACATCAAGAC CACTGCGAGCGCTA | GGAAATACCTTCTTC GATATCCGCTCCA | CTCAACCAGCCGTC CTTTAAGAGCTTTG | ACTGATGCCTTCTTC AATGTCAGCACCG | 229 | C5 |
| *YKL189W 1* | ATTCAGCTCACCCT CGCTGACTCAGGAT | ACCCTTTTGTTGTAG AGCAACTTCCGCA | GTTTTCAAGCCCAA GCTTAACCCAGGAC | GCCTTTTTGTTGCAA GGCGACTTCAGCG | 352 | C5 |
| *YKL188C 1* | TAATGAAGATTTACC GCAACCGTTGGGC | GGAACCCGGGAATG ATATGACGGCT | CAAGCTGCTCTTGC CACAGCCATTTGGA | AGAACCAGGTAACG ACATGACCGCC | 355 | C5 |
| *YKL188C 2* | TGGAATCGAGCATA GTACCAAGCCAGCT | TATGGGTGAAGGAA CTTTGGCACTGGGC | AGGGATGCTACACA AAACTAAACCGGCA | CATGGGCGAAGGTA CCTTAGCTTTAGGT | 334 | C5 |
| *YKL188C 3* | AGATGCGCTTGCCA TAGTTAACCTAGTG | GTCCACACTAGTAA GAGCTCAATACGCA | GCTAGCTGAAGCCA TGGTCAATCTGGTA | TAGTACCTTGGTTCG TGCCCAATATGCT | 460 | C5 |
| *YKL187C 2* | GGGGTGACTTAGCT CTGTTCTATTACTC | CGTTTGCGTTGGTG TAGCCACTGTATTG | TGGATGTGACAACT CGGTACGGTTTGAG | TGTCTGTGTCGGCG TTGCTACCGTTTTA | 439 | C5/ D1 |
| *YKL187C 1* | CAGCGAAGCAGAAA CTAACCTTGAGAGG | TGCTGATACTCCCG CGCTGACG | TAAGCTGGCGCTGA CCAATCTGCTCAAA | CGCCGACACCCCAG CTTTAACC | 220 | D1 |
| *YKL186C 1* | GCATATCAACGTGC CGGATCCCGGAATA | ATTAGCGCATTTGGA TGATCCGGACTCC | ACAGATTAAGGTAC CACTACCTGGGATG | CTTGGCTCACTTAGA CGACCCAGATAGT | 211 | D1 |
| *YKL185W 1* | TTCGCTCAATGTGT CGCCAATTCTTTCC | ACCTAAGCCAGACC CATAGTTCAATGGA | CAGCTTGAACGTTA GCCCTATCTTGAGT | GCCCAAACCGCTAC CGTAATTTAAAGGG | 208 | D1 |

| Amplicon | WT_F | WT_R | syn_F | syn_R | bp | chunk |
|---|---|---|---|---|---|---|
| *YKL185W 2* | AAGATCATCCTCACCTGTGCGTCCCAAG | CGATGGCCTCCAACAGGGCGAA | TCGTAGCAGTAGCCCAGTTAGACCAAAA | GCTAGGTCTCCAGCATGGGCTG | 226 | D2 |
| *YKL184W 1* | CCACGTCGGTTCAGGCGCTTCT | AGATGCCAAAGTGAACGCTGTAGCTACA | TCATGTTGGCAGCGGTGCCAGC | GCTAGCTAAGGTAAAAGCGGTGGCAACG | 268 | D2 |
| *YKL183W 1* | TTTGTACCATGCATCAAGACGGCTACCA | TCCAAGCCCAGAATACGTGGCGTTGAGA | CTTATATCACGCTAGCCGTAGATTGCCT | ACCCAAACCGCTGTAGGTAGCATTCAAG | 421 | D2 |
| *YKL182W 3* | ATCCAACTCTGCTCTTTTTAGGGCCGTC | GGCAGTAACTACGTAGTGAGCCAATTGA | GAGTAATAGCGCCTTGTTCAGAGCTGTT | AGCGGTGACAACATAATGGGCTAATTGG | 340 | D2 |
| *YKL182W 4* | GGCCCCATCTGGACTGGATCAATCAAGA | TGAACCATCAAAAGTGTCGTAAACGGGG | AGCTCCTAGCGGTTTAGACCAAAGCCGT | GCTGCCGTCGAAGGTATCATAGACTGGA | 208 | D2 |
| *YKL182W 1* | CCTTTCAGGTTCCATTTCCGAGAGAATC | ATCATCTGGGTTAATGTCGAGAGTACCG | TTTGAGCGGCAGTATCAGTGAGCGTATT | GTCGTCAGGATTGATATCCAAGGTGCCA | 208 | D2 |
| *YKL182W 5* | GGCAACATCCTCTACTGATGAAGAAAGC | TGGCAAGCTGACGGGCTTACCATCCATA | AGCTACCAGTAGCACCGACGAAGAATCA | AGGTAATGAAACTGGTTTGCCGTCCATG | 328 | D2 |
| *YKL182W 2* | TTCTTCTGCTTCCGTCCGTGCTTTGATT | GGCTTCACCAGTCAAAACTACAACGTCA | CAGCAGCGCCAGTGTTAGAGCCTTAATC | AGCTTCGCCGGTTAAGACAACGACATCG | 214 | D3 |
| *YKL182W 6* | GAAATCTGCTGTCAAGCCTCGCCCACTT | GGAGCCAGTCAAATCATAAACGTCCTGG | AAAGAGCGCCGTTAAACCAAGACCTTTG | ACTACCGGTTAAGTCGTAGACATCCTGA | 259 | D3 |
| *YKL181W 1* | CCGTGGTCGCTCTGCCATCATTCTAGAT | AGCTATGCGTTCGCCAGAAATCGGATAT | TAGAGGCAGAAGCGCTATTATCTTGGAC | GGCGATTCTTTCACCGCTGATTGGGTAG | 223 | D3 |
| *YKL179C 1* | AGAACTCGTTGGGGACATTTTATGGGAC | AAGTACGGCTCAAGAGGAAGAACGAAAC | GCTTGAGGTAGGACTCATCTTGTGACTG | GTCAACCGCCCAAGAGGAAGAAAGAAAT | 220 | D3 |
| *YKL179C 2* | GTAAGATTCCGATTCCGCCACAACGCTA | TGAGGAACTAAGTGGTGCTCTTGCAAAG | ATAGCTTTCGCTTTCAGCAACGACTGAG | CGAGGAATTGTCAGGCGCCTTGGCTAAA | 208 | D3 |
| *YKL178C 1* | TGAAGCTGGAGAGTAGCAAAGACTTTCC | AGAGAGCCGTAACCCATTTTCTACAGAC | GCTGGCAGGGCTATAACACAATGATTCA | GGAGTCAAGAAATCCTTTCAGCACCGAT | 373 | D4 |
| *YKL176C 1* | GGGTGATCCAATGCTGTTAGAGCTCACA | TGGACCGAAACTAGGCTTCCTTCCTCTC | TGGGCTACCGATTGAATTGCTTGAAACG | CGGTCCAAAGTTGGGTTTTTTGCCATTG | 373 | D4 |
| *YKL176C 2* | CGTCAAATCGGAAGATCTTACGGGAGAA | CAAAGCGGCAAGCACAGGAACTACTAAG | GGTTAAGTCACTGCTACGAACTGGGCTG | TAAGGCTGCTTCAACCGGTACCACCAAA | 418 | D4 |
| *YKL175W 1* | CTATGGTCTTTCTCTAAGTGCCGGATCT | GTTTAGGTCACAACACTCGCCTTCTGAT | TTACGGCTTGAGCTTGTCAGCTGGTAGC | ATTCAAATCGCAGCACTCACCTTCGCTC | 457 | D4 |
| *YKL175W 2* | TTTTCTTTCGAGCCGCCATCATCATTCC | ACCGAGAACGGCAGTAATCAAAATGGCA | CTTCTTGAGCTCAAGACACCACCACAGT | GCCCAAGACAGCGGTGATTAAGATAGCG | 424 | D4 |
| *YKL174C 1* | CGCGCTTCGGATAGCTCTTCTTCTC | TCGTGACCAGGCTATACCATACTACGAT | AGCTGATCTAATGGCACGACGACGG | CAGAGATCAGGCCATCCCTTATTATGAC | 244 | D4 |

| Amplicon | WT_F | WT_R | syn_F | syn_R | bp | chunk |
|---|---|---|---|---|---|---|
| *YKL174C 2* | CGACGAAAACCATATACACACTCCACTG | TCAAGAGAACGAGGAGGCCGAGCATTTC | GCTGCTGAACCAGATGCAAACACCTGAA | CCAAGAGAATGAGGAGGCTGAGCACTTT | 211 | D5 |
| *YKL173W 1* | TGGACCTCTTCACTCAGGTAAGACCTCT | TGCAAGTGCGACTGCCGTTTCATCCATA | CGGTCCATTGCATAGCGGCAAAACTAGC | AGCCAAAGCAACAGCGGTTTCGTCCATG | 274 | D5 |
| *YKL173W 2* * | ACTTAATGGTTCCACTCTACTCTGCACC | ATATACGCTACCCCATAGCCTTGTGGTA | GTTGAACGGCAGTACCTTGTTGTGTACT | GTAAACTGAGCCCCACAATCTGGTAGTG | 484 | D5 |
| *YKL173W 3* | TTTACCAGGTCTGTCGATTAGTGTAGCC | AATTGGTTCTTCTGCCAAAGGCCCCTCT | CTTGCCTGGCTTAAGCATCTCAGTTGCT | GATAGGTTCTTCAGCTAATGGACCCTCC | 334 | D5 |
| *YKL172W 1* | GCAATCACTGGACGCTGTTTTAGTAGCA | TCCGTTAGGCCTGCCTCTATCGAACCTT | ACAAAGCTTAGATGCCGTCTTGGTTGCT | ACCATTTGGTCTACCACGGTCAAATCTC | 379 | D5 |
| *YKL171W 1* | TCCAGCATCGAATCGCAGCTATGGAAGA | TATTGCATCCGGTGAGCTCCAAAGTTCG | ACCTGCTAGCAACAGATCATACGGTCGT | GATAGCGTCTGGGCTTGACCACAATTCA | 448 | D5 |
| *YKL171W 2* | TTCGAACCCATTGAGTGGAAGCGCAGGT | GGTCAATCTGCATCTCTCTGCAAATGGT | CAGCAATCCTTTATCAGGTTCAGCTGGC | AGTTAAACGACAACGCTCAGCGAAAGGG | 397 | D5/ E1 |
| *YKL171W 3* | CGCTTCGCCAGAACTTCTAAGATACTCA | AGAGCGTAACCTAGGTTCGAACTCATGA | TGCCAGCCCTGAATTGTTGCGTTATAGC | GCTTCTCAATCTTGGTTCAAACTCGTGG | 334 | E1 |
| *YKL168C 2* | ATCACTGCCCAATGATGCGTCGGGATTC | TCGATTTGCAGGAGCAGATGTGCAATGT | GTCTGAACCTAAGCTAGCATCTGGGTTA | AAGATTCGCTGGTGCTGACGTTCAATGC | 379 | E1 |
| *YKL168C 1* | ACTTCTTGAACGTGGACGCCGTATGCCA | CTCTGAGCCACCAAGACGATCAACCGTA | TGAACGGCTTCTAGGTCTTCTGATACCG | TAGCGAGCCTCCTCGTAGAAGCACTGTT | 265 | E1 |
| *YKL166C 1* * | TAGGTGAACTCTCCCAAATGAGCCAGTT | GGTGCATGAGGAATGTTCTTCCAACACA | CAAATGGACACGACCGAAGCTACCGGTA | TGTTCACGAGGAATGCAGCAGTAATACC | 202 | E1/ E2 |
| *YKL165C 1* | CGACCCCTCCGTTCTTAAAAGGTAGAAA | CGCATTCTTTGGAACTGGTAACGTCGCT | GCTACCCTCGGTACGCAACAAATAAAAG | TGCTTTTTTCGGTACCGGCAATGTTGCC | 265 | E2 |
| *YKL165C 2* | TATATTCACGGATAGCTCTCTCACCCCA | CCAAAAAGCGTCACCATTAAGCCATGCA | GATGTTAACACTCAACTCACGAACACCG | TCAAAAGGCTAGCCCTTTGTCACACGCT | 346 | E2 |
| *YKL165C 3* | AAAAGCATCCAGCTCGATGGAAGATTGC | TACCAGAATGCCAACTGAATCCCGTCCT | GAAGGCGTCTAACTCAATACTGCTTTGG | CACTCGTATGCCTACCGAAAGTAGACCA | 271 | E2 |
| *YKL164C 1* | GGAGACTGGAGCAACAGTAGTATTGGTG | AATTGCCACTACTGCTTCCTCCAAGGCT | ACTAACAGGGGCGACGGTGGTGTTAGTA | TATCGCTACCACCGCCAGTAGTAAAGCC | 463 | E2 |
| *YKL163W 1* | TTCCTCCGTCGCCTCATCTAAAGCAAAG | GGCCTGAACTTGGCCGTCAGAAATCTGA | CAGTAGTGTTGCTAGCAGCAAGGCTAAA | AGCCTGGACTTGACCATCGCTGATCTGT | 361 | E3 |
| *YKL162C 1* | TGTGCTTCTCTCACTTGGGCTAATCCAT | TGACAAAGGGTGGCAACAGTGCTTTGTG | GGTTGAACGCTCTGAAGGTGAGATCCAC | CGATAAGGGTTGGCAACAGTGTTTCGTT | 487 | E3 |
| *YKL161C 1* | TAGTATGCTTTCAAACGATCTTCCCGGG | TATTTGGTCAACAGGCTGTATCTTGGCC | CAAGATTGATTCGAAGCTACGACCTGGA | CATCTGGAGCACCGGTTGCATTTTAGCT | 208 | E3 |

| Amplicon | WT_F | WT_R | syn_F | syn_R | bp | chunk |
|---|---|---|---|---|---|---|
| *YKL159C 1* | TCCATCGTTCGTGGGACATCTATCAATG | CTGCCCCAGTCATGATATATCTCAGCAT | ACCGTCATTGGTTGGGCAACGGTCGATA | TTGTCCATCACACGACATCAGCCAGCAC | 373 | E4 |
| *YKL155C 2* | TCTTCTTCCCGTTCCTTTTAGCCTCGAC | CAGGGCAAGGCAAATTACGCTTAGACCT | ACGACGACCGGTACCCTTCAATCTGCTA | TAGAGCTAGACAAATCACCTTGCGTCCA | 361 | E5 |
| *YKL155C 1* | AGATGCGGGAATAGAAGACCGCAAGTTA | CCCAGCGACTGGTATAGTTGCCTTGAAT | GCTAGCTGGGATGCTGCTTCTTAAATTG | TCCTGCTACCGGCATCGTCGCTTTAAAC | 289 | E5 |
| *YKL154W 1* | GCTGCTAACCACAGATTCAGTAAGACCA | CCCGTCGGTAGACTGTAAAACGTCTAGT | CTTATTGACTACCGACAGCGTTCGTCCT | ACCATCAGTGCTCTGCAAGACATCCAAG | 466 | E5 |
| *YKL151C 1* | AGAGTCTCCCTTCTTGCCTATGGCATCA | CGCTGGTACGGTCATTAAATCGTACACC | GCTATCACCTTTTTTACCGATAGCGTCG | TGCCGGCACCGTTATCAAGAGCTATACT | 388 | E5/F1 |
| *YKL150W 1* | GTTGGTCCTAGCATCTGCTCTGTTTGCT | GGTACCGGCACCTAACAAGGTGATTGAC | TTTAGTTTTGGCTAGCGCCTTATTCGCC | AGTGCCAGCGCCCAATAAAGTAATGCTT | 259 | F1 |
| *YKL149C 1* | TGACTTATTAGGCGGTTCGTGGCTTCTT | TATGCTGAGCCACGATTGGCCCAATGGA | GCTTTTGTTTGGTGGTTCATGTGAACGC | CATGTTATCACATGACTGGCCAAACGGT | 235 | F1 |
| *YKL148C 1* | ACGGGATTCCTTTCTATTAGCAGCGGAA | TGACACTTTACAGCCTGGGTTGCCACAC | TCTACTTTCTTTACGGTTGGCGGCACTG | CGATACCTTGCAGCCAGGTTTACCTCAT | 352 | F1 |
| *YKL148C 2* | ACATGTGTGAGCAGAGGTACAAGAGAAG | TGATTGGCTAGGTGACCAGGACTCCATC | GCAGGTATGGGCGCTAGTGCAGCTAAAA | AGACTGGTTGGGCGATCAGGATAGTATT | 427 | F1 |
| *YKL146W 2* | CCCAAGTCAACCTGCACTTTCCACCTCA | GAATCCACCAGGAGCCCTGATTTCTTCG | TCCTTCACAACCAGCTTTGAGTACTAGC | AAAACCGCCTGGGGCTCTAATTTCTTCA | 208 | F2 |
| *YKL146W 1* | ATTTAGCGCCCTTTGTCTACTCTCGTGT | ATCTGCTATTAACGCGGTCCCACTTAAC | TTTCTCAGCTTTGTGCTTGTTGAGCTGC | GTCAGCGATCAAAGCAGTACCTGACAAT | 358 | F2 |
| *YKL145W 1* | AGAAAGCGATACAGGGTTAGCACCCTCC | TGGAGAAACACGTTCACCTAATCCGACG | GGAATCAGACACCGGTTTGGCTCCAAGT | AGGGCTGACTCTTTCGCCCAAACCAACA | 286 | F2 |
| *YKL144C 1* | ACTAACGAGTCCCATGCCATCAGTTTGA | ATGGATTTGGCCCATGGATGAAGAGACG | TGAGACCAAACCCATACCGTCGGTTTGG | TTGGATCTGGCCAATGGACGAAGAGACC | 247 | F2 |
| *YKL143W 1* | TTCCATGAGTTCAAGTGCCATTGCGCGT | ATCTTTCTTAGCCAGTTTACGGCCACCA | CAGTATGTCAAGCTCAGCTATCGCTAGA | GTCCTTTTTGGCTAACTTTCTACCGCCG | 235 | F2 |
| *YKL142W 1* | TCCTGCTATTTCAGACGGAGTGTTCCCT | GGAGAGACCTAGATAGCGGGCAACATCA | CCCAGCCATCAGCGATGGTGTTTTTCCA | ACTCAAGCCCAAGTATCTAGCGACGTCG | 271 | F2 |
| *YKL140W 2* | TACCTTGGCCAAATCATCGCCCGGCTTT | GGTATCGGCATCTTTGCCCCAAATCAAA | CACTTTAGCTAAGAGCAGCCCAGGTTTC | AGTGTCAGCGTCCTTACCCCAGATTAAG | 496 | F3 |
| *YKL140W 1* | TCTCAGTTATAGAACTCACTCGGCAGAC | GGCACTTAACCGTCTTTGCTGCTCCTTA | CTTGTCATACCGTACCCATAGCGCTGAT | AGCTGACAATCTACGTTGCTGCTCTTTG | 202 | F3 |
| *YKL139W 1* | GCCACCAAAGAGAATACGGACTGATAGT | GCTCCTTTGCTGCGTTAGAACCGAAACT | TCCTCCTAAACGTATCAGAACCGACTCA | TGATCTTTGCTGGGTCAAGACGCTGACA | 430 | F3 |

| Amplicon | WT_F | WT_R | syn_F | syn_R | bp | chunk |
|---|---|---|---|---|---|---|
| *YKL139W 2* | TTCGCGAGCTGATT ACACTAACCGTGTC | CGGCATATCGTAAA GCGTTGGCCAGCTA | CAGCAGAGCCGACT ATACCAATAGAGTT | TGGCATGTCATACAA GGTAGGCCATGAG | 241 | F3 |
| *YKL135C 2* | TTTGCCTTTTCTCCT ATTTGAGTCGGGC | CTATGTCTGGTTGCT AGGACAGCATCCC | CTTACCCTTACGTCT GTTGCTATCTGGT | TTACGTTTGGTTATT GGGTCAGCACCCA | 412 | F4 |
| *YKL135C 1* | AGGCGGCGTAGACA TCAGTGATACAAAA | GGATTTATGCGTTGA GTTGGGTGTCGTC | TGGTGGGGTGCTCA TTAAGCTAACGAAG | AGACTTGTGTGTCG AGTTAGGCGTTGTT | 427 | F4 |
| *YKL134C 2* | ATTCAATCTTCTCAC GTCTGGGGACCAT | GCTCGTGAGACCAT GGGATAGGGATTAC | GTTTAAACGACGAA CATCAGGACTCCAG | ATTGGTTCGTCCTTG GGACAGAGACTAT | 217 | F4 |
| *YKL134C 1* | TGCGCTTAACTTCG AAGACACTTCTGGA | GTTTAGCCAGGTTTC TTTGCAGCAAGCA | AGCTGACAATTTGCT GCTAACTTCAGGG | ATTCTCACAGGTCA GCTTACAGCAAGCT | 304 | F4 |
| *YKL133C 1* | AATACCGTAGTGAG CAAGGGATAATGCC | TCTCAATCGGAGTAA TCTTTCTGGGGCC | GATGCCATAATGGG CCAAACTCAAAGCT | CTTGAACAGATCAAA CTTGAGCGGTGCT | 373 | F4 |
| *YKL132C 1* | ACGTATTAGCGGAT CAAGTAAGGGAGAC | CACAGACACATGGG GCACTCTTGAAGTT | TCTGATCAATGGGT CCAACAATGGGCTA | TACCGATACCTGGG GTACCTTGGAAGTC | 373 | F5 |
| *YKL130C 1* | TAAAATGGAGTGCC ACGCAGCTGACAAG | CGAATTAGACCCAG AGGCGGATTCGTTT | CAAGATACTATGCCA AGCGGCGCTTAAA | TGAATTGGATCCTGA GGCTGACAGCTTC | 409 | F5 |
| *YKL129C 4* | CAATGCAGAAGCTA AACCGTCGCTAAAC | ATTCGAAGCTGCATA CGACTTTCCAGGT | TAAAGCGCTGGCCA AGCCATCTGAGAAG | GTTTGAAGCCGCTT ATGATTTCCCTGGC | 367 | F5 |
| *YKL129C 2* | TAATGGCAACTCCG AAGATGATCCGGAA | TGCTGCAAAAGTAA GCTCGAAACACAGC | CAAAGGTAACTCGC TGCTGCTACCACTG | AGCCGCTAAGGTTT CAAGCAAGCATTCA | 292 | F5 |
| *YKL129C 1* | GGGTTTTGAGCTGT GTTTCGAGCTTACT | CCGCTCAAAGATAA GCGATTCAGCACCG | TGGCTTGCTTGAAT GCTTGCTTGAAACC | TAGAAGCAAAATCTC AGACAGCGCTCCA | 373 | F5 |
| *YKL129C 3* | AGCTGATCTGCCAA ATTTAGAGTGCAGG | TGATTACACTTGGGA TGGGGACACCCTT | GGCGCTACGACCGA ACTTGCTATGTAAA | CGACTATACCTGGG ACGGTGATACTTTG | 487 | F5 |
| *YKL128C 1* | AATGACACCTGAGT GACAAGTAAGCGAG | CATTGAGTCGTGGA CACCAGTGTTAGCG | GATAACGCCGCTAT GGCAGGTCAAGCTA | TATCGAGAGCTGGA CCCCTGTTTTGGCT | 355 | G1 |
| *YKL127W 1* | CTTGGCACGTTCAT TTCCTACATCCTCA | GATAACGGTTAAAG ACTCATCAGCTGGG | TTTAGCTAGAAGCTT CCCAACCAGTAGC | AATGACAGTCAAGC TCTCGTCGGCAGGA | 436 | G1 |
| *YKL127W 2* | TGTATCAAGGCCAA ACGTTTGTGGCTCC | GCGGACTGTTGGTT CGTCTGTTCCTAAA | CGTTAGCAGACCTA ATGTCTGCGGTAGT | TCTAACGGTAGGTT CATCGGTACCCAAG | 334 | G1 |
| *YKL126W 1* | ATACGATGCATCCT CATCGACCTCTACT | CAAATCTTCATCAGC TGGACCTTGGTTC | TTATGACGCTAGTAG CAGCACTAGCACC | TAAGTCTTCGTCGG CAGGGCCTTGATTT | 340 | G1 |
| *YKL126W 2* | TGGATTATTGAGCC GTGATCCGACAAGA | TGAGCTACCTAGCT GTTCATTTCCAACG | CGGTTTGTTATCAAG AGACCCAACCCGT | GCTTGAGCCCAACT GTTCGTTACCGACA | 286 | G2 |
| *YKL125W 2* | CGTGAAATCTGCCC TAGATGATCTGGAT | GTGTGGAGGCAATT CCCACCACTTCTCA | TGTTAAGAGCGCTTT GGACGACTTAGAC | ATGAGGTGGTAATT CCCACCATTTCTCG | 229 | G2 |

| Amplicon | WT_F | WT_R | syn_F | syn_R | bp | chunk |
|---|---|---|---|---|---|---|
| *YKL125W 1* | AGCAGCCGACGGTAGTCAAAGTGATAGT | CGACCTGGTATAGTACGTGGGTAGAACA | TGCTGCTGATGGCTCACAATCAGACTCA | GCTTCTAGTGTAATAGGTTGGCAAGACG | 262 | G2 |
| *YKL124W 1* | TAGCTTCCGTGAAGCTCTCTCTGAAGGC | AGTTTGCGCCGTTATCCTAGAGGATGAC | CTCATTTAGAGAAGCCTTGAGCGAAGGT | GGTTTGAGCGGTGATTCTGCTACTGCTA | 352 | G2 |
| *YKL122C 1* | CTTCGGTTCTGTAGCAGTTTTCTTAGCC | TGCATTAGGCCCACGAGGCGTATCT | TTTTGGTTCGGTGGCGGTCTTTTTGGCA | CGCTTTGGGTCCTAGAGGTGTTAGC | 208 | G2 |
| *YKL121W 3* | CAGGAGCCGACATTCCTCTATTGCAGAT | CTTACCATCATGACTGAACGTACAGCAG | TAGATCAAGACACAGTAGCATCGCTGAC | TTTGCCGTCGTGTGAAAAGGTGCAACAA | 430 | G2 |
| *YKL121W 1* * | AGTCAGTGCCTCTAGAGCAAACTCACTA | TTTCCTCTCAGGGTGCCATAGTTTTGCA | TGTTTCAGCTAGCCGTGCTAATAGCTTG | CTTTCTCTCTGGATGCCACAACTTAGCG | 280 | G3 |
| *YKL121W 2* | CCAAAGTGGTTCCTCACGTCATAGAGGC | GACAGGGGAGTTATGGGCATGAAATGAT | TCAATCAGGCAGTAGCAGACACCGTGGT | AACTGGACTATTGTGAGCGTGGAAGCTG | 367 | G3 |
| *YKL120W 1* | AGTTAACGTCTTTTCTGGTGCCGCATCT | GACATCCCATGGGTTCATAACGACGGCA | TGTCAATGTTTTCAGCGGCGCTGCTAGC | AACGTCCCAAGGATTCATGACAACAGCG | 370 | G3 |
| *YKL119C 1* | ATGGACCGGAAAGTTTGTTGAAGATCCG | GCGACACGCAATACCTATGCAGTCACTT | GTGAACTGGGAAATTGGTGCTGCTACCA | AAGACATGCTATCCCAATGCAGAGCTTG | 388 | G3 |
| *YKL117W 1* | TGCCCCTGAGTTAACCATTAAGCCATCA | GAAATCTCCCATGTCCAAATTACCACCG | CGCTCCAGAGTTGACTATCAAACCTAGC | AAAGTCACCCATATCTAAGTTGCCGCCA | 487 | G3 |
| *YKL116C 1* | TCTGCTACGCTGTCTTGCTGAAGCGTTT | AGGAGACTTATTAGCTGCCGTCATGGCC | ACGTGATCTCTGACGAGCGCTGGCATTA | TGGTGATTTGTTGGCCGCTGTTATGGCT | 439 | G3 |
| *YKL116C 2* | AAAATTTCCTGAACCAATGGGCCTGACC | CCAAAGGGTTGTTTCCTTACCAACAGTG | GAAGTTACCGCTGCCGATTGGTCTAACT | ACAAAGAGTCGTCAGTTTGCCTACCGTT | 211 | G4 |
| *YKL114C 1* | ACGCGATTTAGCACCTAATTTCTGCAGC | ATGCCATACATTTGCAGCAGGCTACGAT | TCTGCTCTTGGCGCCCAACTTCTGTAAG | CTGTCACACCTTCGCTGCTGGTTATGAC | 388 | G4 |
| *YKL113C 1* | ACCAACACCTCTGATGCTTTCACAGTAG | TCAATGTGCTGAGTTGGCAAAGAAGGGA | GCCGACGCCACGAATTGATTCGCAATAA | CCAATGCGCCGAGTTAGCTAAAAAAGGT | 244 | G4 |
| *YKL112W 2* | CAATGCTGCATCCTCAGAAACCTCATCT | GTGGTTACCGCTGTCATTACCATGAGTA | TAACGCCGCTAGTAGCGAAACTAGCAGC | ATGATTGCCTGAATCGTTGCCGTGGGTT | 412 | G4 |
| *YKL112W 1* | CTCACCATCCTCCATACGAGACACGTCT | TAAAAGACCGGACTCCTCCACGTAATGA | TAGCCCTAGTAGTATCAGAGATACCAGC | CAACAAGCCACTCTCCTCAACATAGTGT | 280 | G4 |
| *YKL110C 1* | ACTTACCCCGCCAATAGAAGTCAGGCTT | CCGATGGGACTCTCCACTCTTTGCTATT | TGAAACACCACCGATGCTGGTTAATGAC | TAGATGGGATAGCCCTTTGTTCGCCATC | 280 | G4/ G5 |
| *YKL109W 2* | TGTGGAAACCATTGCTGCTCACAACAGT | ATCAGGGAATGAAGCCTTATGAGGGATC | CGTTGAAACTATCGCCGCCCATAATTCA | GTCTGGAAAGCTGGCTTTGTGTGGAATT | 226 | G5 |
| *YKL109W 1* | AGATTTCTCTCCAGCAGATTCAGTCTCG | ATCTGGCTCAACCTTATCATCTTGCCCT | TGACTTTAGCCCTGCTGACAGCGTTAGC | GTCAGGCTCGACTTTGTCGTCTTGACCA | 340 | G5 |

| Amplicon | WT_F | WT_R | syn_F | syn_R | bp | chunk |
|---|---|---|---|---|---|---|
| *YKL108W 1* | TCTCCTAAAATCGTC GCCTGCAGATCGT | ATACCCCGATGGTTT GCCAGATTCTAGT | CTTGTTGAAGAGCA GCCCAGCTGACAGA | GTAACCGCTAGGCT TACCGCTTTCCAAA | 211 | G5 |
| *YKL107W 1* | AAGTAGAGAGCTGG CCCAAAGAAATGCC | GTCATCTGGGTCTC CCACCTGACCATTT | CTCACGTGAGTTAG CTCAACGTAACGCT | ATCGTCAGGATCAC CAACCTGGCCGTTA | 382 | G5/ H1 |
| *YKL106W 1* | TTCTCCACCAGGAT ACGGTTCTCGTGTG | GACTCCGCTAAGCG ACAATCTACCATCA | AAGCCCTCCTGGTT ATGGCAGCAGAGTT | AACACCTGACAAGC TTAAACGGCCGTCG | 277 | H1 |
| *YKL105C 1* | GGATGATGCAGTGT AAGGGGAAGAATCA | ATCGAGACGATTGC CTGATCATGTCCCT | ACTGCTAGCGGTAT ATGGACTGCTGTCG | CAGCCGTAGATTAC CAGACCACGTTCCA | 430 | H1 |
| *YKL105C 2* | TGAACTACCACGTA GAGTTCTTAGGCAC | TACCACAAGAGTTG CACCTGACCTTTTG | GCTTGAGCCTCTCA AGGTACGCAAACAA | CACTACCCGTGTCG CTCCAGATTTGTTA | 316 | H1 |
| *YKL105C 3* | GCTGCCAGTGAGAC TGTTTGAATCTGCA | CAAAGGACAGGAAG TAGCATCCGAGGTC | TGAACCGGTCAATG AATTGCTGTCAGCG | TAAGGGTCAGGAAG TTGCTAGTGAGGTT | 286 | H1 |
| *YKL104C 1* | TACACGGTCATCTG ACAGCGATAGAGCA | TGAAACTGCGGATA CCATGCTGGCTCTA | AACTCTATCGTCGCT TAAGCTCAAGGCG | CGAAACCGCTGACA CTATGTTAGCCTTG | 220 | H2 |
| *YKL104C 2* | AGATCCATCTTCTG ATAGGAAAGCCCTG | AGGGTCCCCTTTAC TGATTGGTGTCAAA | GCTACCGTCTTCGC TCAAAAAGGCTCTT | GGGTAGTCCATTGT TAATCGGCGTTAAG | 256 | H2 |
| *YKL103C 2* | ATCCTTGGACCCTG TAGCAGCTCTGATG | CGGATCGTTGACAA GACAAGGCGCAAAA | GTCTTTACTACCGGT GGCGGCACGAATA | TGGTAGCTTAACCC GTCAAGGTGCTAAG | 445 | H2 |
| *YKL103C 1* | TAGGGGTGTAGAAT CAACCAAGGCGCTT | TGGATCTCATGTGG ACGCTTTGACGGTC | CAATGGGGTGCTGT CGACTAAAGCTGAC | CGGTAGCCACGTTG ATGCCTTAACCGTT | 211 | H2 |
| *YKL101W 5* | CTCTCCACACTATG CGTCTCCTGAAATT | GGCAGTGATTAGTT CACGAGAAACTCCA | TAGCCCTCATTACG CTAGCCCAGAAATC | AGCGGTAATCAATTC TCTGCTGACACCG | 490 | H3 |
| *YKL101W 2* | AATCCACGCCTCCC CATCTACTAAATCA | CCCAGAGATCGTAA AAGTGTCTGCCGAA | TATTCATGCTAGTCC TAGCACCAAGAGC | ACCGCTAATGGTGA AGGTATCAGCGCTT | 400 | H3 |
| *YKL101W 3* | TAAACCGTCCTTATC TCTAGATCCCCGC | TGGCACACTTAGAT GCTTATCATCACCG | CAAGCCAAGTTTGA GCTTGGACCCAAGA | AGGAACTGACAAGT GTTTGTCGTCGCCA | 379 | H3 |
| *YKL101W 4* | CCAATCTAGAAGTG TAGCCATGTCACAC | CACACTTGCTCGTG AATCGCTTTCTGAA | TCAAAGCCGTTCAG TTGCTATGAGCCAT | AACTGAAGCTCTGC TGTCTGATTCGCTT | 469 | H3 |
| *YKL101W 1* | GAGGTTTTCTTCCA CGCGTGGTAGTCGT | ACTGGAAGGCGCTT CTTCAACTTCAACA | AAGATTCAGCAGTA CCAGAGGCTCAAGA | TGAACTTGGAGCTT CTTCGACTTCGACG | 487 | H3 |
| *YKL100C 1* | AAGCTCTTCCTCAG AGCTGTCACTGAGA | CGTTGCTTCTTTGGT ATCTGCCATGGTC | CAACTCTTCCTCGCT TGAATCTGACAAG | TGTCGCCAGCTTAG TTAGCGCTATGGTT | 355 | H3 |
| *YKL100C 2* | TAGAGCAATTAGGA TTAGGGCCCCTGAT | TTGGAATCCCCTGG TTGTTTTACCAAGG | CAAGGCGATCAAAA TCAAAGCACCGCTC | ATGGAACCCATTAGT CGTCTTGCCTAGA | 418 | H3 |
| *YKL098W 1* | CTCAACAACTGCAA GCTTGGACTCGTTA | TGGGGGTGCACATG CAAACCAGAATTTG | TAGCACCACCGCTT CATTAGATAGCTTG | AGGTGGAGCGCAAG CGAACCAAAACTTA | 331 | H4 |

| Amplicon | WT_F | WT_R | syn_F | syn_R | bp | chunk |
|---|---|---|---|---|---|---|
| *YKL096W 1* | TTTGAAACTTGGCA GCGGTAGTGGCTCA | CTTACTAGTTGGTCT AATCGCAACACCG | CTTAAAGTTGGGTTC AGGCTCAGGTAGC | TTTTGAGGTAGGAC GGATAGCGACGCCA | 277 | H4 |
| *YKL095W 1* | CATTTCATGTCCAC GTTGTGCCAATTCC | GTTGTCCAGATCATC AGTCGTTACTGCT | TATCAGCTGCCCTA GATGCGCTAACAGT | ATTATCTAAGTCGTC GGTGGTAACAGCC | 403 | H4 |
| *YKL094W 1* | AAGCGGATATATCG GGTCAGGCCCATTA | TAGTGAGAAAATCG AATGTCTTGCGCCC | CTCAGGTTACATTG GTAGCGGTCCTTTG | CAAGCTAAAGATGC TGTGACGAGCACCA | 433 | H4/ H5 |
| *YKL093W 1* | AAGGTCGTTTTCTCT ACCCAGGTCAAGA | CCTCCCAGCACCAA AGTTTGGTTTCGAA | TAGAAGCTTCAGCTT GCCAAGAAGCCGT | TCTACCGGCGCCGA AATTAGGCTTGCTT | 331 | H5 |
| *YKL092C 2* | GGAAGCACATTCAT AACCTGACAACCGT | CAATCACATGGATAA TGGTGGGAAGGGA | ACTGGCGCATTCGT AGCCGCTTAATCTC | TAACCATATGGACAA CGGCGGTAAAGGT | 391 | H5 |
| *YKL092C 1* | AATATCACTAGAGTT GGTGGCAGCACTG | AGGTTGGTTTTCCG CTATGGGGGTG | GATGTCTGAGCTATT AGTAGCGGCTGAA | TGGCTGGTTCAGTG CCATGGGTGTT | 256 | H5 |
| *YKL092C 3* | ACCATTTGTCTTCAT TGAGCTCCACCAG | TACCAGTACAGGTG CACTAACCCACTCA | GCCGTTGGTTTTCAT GCTTGACCACCAA | CACTTCAACCGGCG CTTTGACTCATAGC | 274 | H5 |
| *YKL091C 1* | AATAAGGTAGCCAG CTCTTCTCGAACAC | TGAACGATGGAGGG AAGAATACGGTGCC | GATCAAATAACCGG CACGACGGCTGCAA | CGAAAGATGGAGAG AAGAATATGGCGCT | 292 | H5/ I1 |
| *YKL090W 1* | ATTTTCAACACCGC AGATTTCCCGCAGA | TCGTTCTGCTACCTC TTTGGACCACCAC | GTTCAGCACCCCAC AGATCAGTAGACGT | TCTTTCAGCAACCTC CTTACTCCACCAT | 214 | I1 |
| *YKL089W 2* | TGTCCAAAGCCCAT CTGGGAAAGGCGAT | TCTATCAGGTGTCAA CCTTACAGAAGGG | CGTTCAATCACCTAG CGGTAAGGGTGAC | ACGGTCTGGGGTTA ATCTAACGCTTGGA | 259 | I1 |
| *YKL089W 1* | TTTGGATTCAGGCT CGGATTCAGCCAGT | TGGCCTTCCTCTTG GTCTCCCC | ATTAGACAGCGGTA GCGACAGCGCTTCA | AGGTCTACCACGAG GACGACCG | 400 | I1 |
| *YKL088W 1* | ATCGAGTGGTACGA GCAATAGCGAAGAT | GCCCGTCGCACCAA TGAGAATGTGGAAT | TAGCTCAGGCACCT CAAACTCAGAAGAC | ACCGGTAGCGCCGA TCAAGATATGAAAC | 274 | I1 |
| *YKL088W 2* | TTCTGTGATGAGAG ATTGGTCACCACTG | GTCATTGCTCTCATC GGATGCTGTTTCA | CAGCGTTATGCGTG ACTGGAGCCCTTTA | ATCGTTTGACTCGTC ACTAGCGGTTTCG | 400 | I1 |
| *YKL087C 1* | GGCCCTTGGTGTCA GTTTTTTGGAATCA | CACCATACCAAGGA CGAATTCTGACAGA | AGCTCTAGGGGTTA ACTTCTTACTGTCG | AACTATCCCTAGAAC CAACAGCGATCGT | 253 | I1 |
| *YKL085W 1* | CTCTATTAGAGCCG CCAGATTCATCTCA | AGCACCATTCTTTGC TTTGACGACTTCG | TAGCATCCGTGCTG CTCGTTTTATTAGC | GGCGCCGTTTTTAG CCTTAACAACTTCA | 211 | I2 |
| *YKL082C 1* | TCTAGAACCGGGTG CTTTCCTTTTACTC | CCCGGAGCAACGTG ATGATGAAACATCT | ACGGCTGCCTGGAG CCTTTCTCTTTGAT | TCCAGAGCAAAGAG ACGACGAAACCAGC | 427 | I2 |
| *YKL080W 1* | CGTTGTACCAGCAT CTGCCAGCGTGATT | ATCCTTCATGAAGG CATTGCCACCAAGA | TGTCGTTCCTGCTA GCGCTTCAGTTATC | GTCTTTCATAAAAGC GTTACCGCCCAAG | 430 | I2 |
| *YKL079W 2* | TGAGACAACCAAGG AGACCCATTCAACA | TCCGACATGTGCCG GACAAATATCTTTG | CGAGACCACTAAAG AGACTCACAGCACC | ACCAACGTGAGCTG GGCAGATGTCCTTA | 421 | I2 |

| Amplicon | WT_F | WT_R | syn_F | syn_R | bp | chunk |
|---|---|---|---|---|---|---|
| *YKL079W 1* * | CGACAAGACAGAGAGATCAAGATCACAC | TTTTGCGCTGTAGGCTGAATCACGCTCT | TGATAAAACCGAGCGTAGCCGTAGCCAT | CTTAGCTGAATAAGCGCTGTCTCTCTCC | 253 | I2/I3 |
| *YKL078W 2* | GTCGATTGCTGTAACTCAACCCCGTCGT | GACAGCGTCTACTATATCATCTGTTGGC | TAGCATCGCCGTTACCCAACCAAGAAGA | AACGGCATCAACGATGTCGTCGGTAGGA | 451 | I3 |
| *YKL078W 1* | AGCCAGAAGTGACGTCACATCTCCTGTG | GATAAGGTCACCATACCTTTTACCAGCG | CGCTCGTTCAGATGTTACCAGCCCAGTT | AATCAAATCGCCGTATCTCTTGCCGGCA | 382 | I3 |
| *YKL077W 1* | TACGGTCATTGCCGGTGTCACTTTTTCT | AACTTCATTAGGGCCCATATTGTGAGCC | AACCGTTATCGCTGGCGTTACCTTCAGC | GACTTCGTTTGGACCCATGTTATGGGCT | 232 | I3 |
| *YKL075C 1* | TCTTTCCAGTCTTTCTTGGCTAACGGCA | AGATGGTGAAAAGAACTCAGAGGCGTCT | ACGTTCTAAACGTTCTTGTGAGACAGCG | CGACGGCGAAAAAAATAGCGAGGCTAGC | 475 | I3/I4 |
| *YKL074C 2* | TGATCGACAGGCCAAAACCATAGTGCTA | TTCTCCGGCTTACTACAACATGGCCTCA | GCTTCTGCAAGCTAAGACCATGGTTGAG | CAGCCCAGCCTATTATAATATGGCTAGC | 262 | I4 |
| *YKL074C 1* | GCCCGAAATTACAAGCCTTGAATTTGCC | CGGTGAGCTTCCGAAAGCGCCAAAA | ACCGCTGATAACCAATCTGCTGTTAGCT | TGGCGAGTTGCCAAAGGCTCCTAAG | 490 | I4 |
| *YKL073W 3* | CTACGGTTCCGCTGTTGGCAGTTTAGCA | TGCAACAGACATCCCTTCACTAACCAGA | TTATGGCAGTGCCGTCGGTTCATTGGCT | AGCGACGCTCATACCTTCTGAGACTAAG | 412 | I4 |
| *YKL073W 2* | CATGGGGAGCGGTTCTATTAAGGCCTCA | AGAGCCTCCCGCCAAAATGACCCCATTT | TATGGGTTCAGGCAGCATCAAAGCTAGC | GCTACCACCAGCTAAGATAACACCGTTG | 469 | I4 |
| *YKL073W 1* | TTCCTCTGATAAGTGCTCGTCAGGAGTT | ATGCTCATGCAAACGCGAACGTTCTGAA | CAGTAGCGACAAATGTAGCAGCGGTGTC | GTGCTCGTGTAATCTGCTTCTTTCGCTT | 223 | I4 |
| *YKL072W 1* | GACTACGGTGGCATCACTACTTAGTCTT | GAGCTTACCAAGGGATCCACCGTGTATA | TACCACCGTTGCTAGCTTGTTGTCATTG | CAATTTGCCCAAACTACCGCCATGGATG | 370 | I5 |
| *YKL072W 2* | GAGAAGTGCTTCGAGAAAGGAAACATGC | CGAATTGCGCCTAAAATACTCCCCCCTA | TCGTTCAGCCAGCCGTAAAGAAACCTGT | GCTGTTTCTTCTGAAGTACTCACCTCTG | 328 | I5 |
| *YKL071W 1* | CTCTATTCGTGGATCACCTTCATTACCC | TAGTATCGGACCCAGAGCGTTTGTGCTA | TAGCATCAGAGGTAGCCCAAGCTTGCCA | CAAGATTGGGCCTAAGGCATTGGTTGAG | 271 | I5 |
| *YKL070W 1* | TCAGAGCGTCGATTCATACATTTCTCCG | ACTTCGTACATTTTCCCCCTCAGGTCTT | CCAGTCAGTTGACAGCTATATCAGCCCA | TGATCTAACGTTTTCACCCTCTGGACGC | 208 | I5 |
| *YKL069W 1* | AAACGCTTCGTCATTGATCTGGCATGCG | GACGCCCAATGTCTTACCATCATTGGAT | TAATGCCAGCAGCTTAATTTGGCACGCT | AACACCTAAGGTTTTGCCGTCGTTACTG | 301 | I5/J1 |
| *YKL068W 1* | CTCACCATTTGGGTCGTTAAACTCCTCA | ACTGCCCCCCAGCTGATTTTGATTACCA | TAGCCCTTTCGGTAGCTTGAATAGTAGC | TGAACCACCTAACTGGTTTTGGTTGCCG | 463 | J1 |
| *YKL068W 3* | AGTTGGTTCAGGGTCGCTGTTTGGC | GGACCCTTGCTGTTGATTATTTTGGCCA | TGTCGGCAGCGGTAGCTTATTCGGT | ACTACCTTGCTGTTGGTTGTTTTGACCG | 289 | J1 |
| *YKL068W 2* | CACCTTTTCAAATTCCGCATCAGGAGGT | GGAAGGTACTGTCGTAGAAGCTGTGTTG | TACTTTCAGCAACAGTGCTAGCGGTGGC | ACTTGGAACGGTGGTGCTGGCGGTATTA | 490 | J1 |

| Amplicon | WT_F | WT_R | syn_F | syn_R | bp | chunk |
|---|---|---|---|---|---|---|
| *YKL065C 1* | GTTTCCTTTCTTGGAAGCTTCAGCAGCT | CAACAGTGGCTCAATCGGGTCATCTGCT | ATTACCCTTTTTACTGGCTTCGGCGGCA | TAATTCAGGTAGCATTGGTAGCAGCGCC | 382 | J1/J2 |
| *YKL064W 3* | TGTTGGTCACTTGAAGTCAGACTCACGT | AGAGGGTCTTAACCCTGCGGGTTTC | CGTCGGCCATTTAAAAAGCGATAGCAGA | GCTTGGACGCAAACCAGCTGGCTTT | 310 | J2 |
| *YKL064W 2* | CCCTTCCACTTTATCGTCTATGTCTGTG | AGCACCATAGGCTTGTGATTTACGACGT | TCCAAGTACCTTGAGCAGCATGAGCGTT | GGCGCCGTAAGCTTGGCTCTTTCTTCTA | 319 | J2 |
| *YKL064W 1* | ACGCCGTCGGTCATCTGCAAATAGAACA | ATCGAACGGAAATGCATCCTCATCGCTA | TAGAAGAAGAAGCAGCGCTAACCGTACC | GTCAAATGGGAAAGCGTCCTCGTCTGAG | 472 | J2 |
| *YKL062W 1* | TTCTGCGAACAATAGCATATCTTCGCCG | GTTTGTCTTGGGAGAAGTTGCATGTTGG | CAGCGCTAATAACTCAATCAGCAGCCCA | ATTGGTTTTTGGGCTGGTAGCGTGTTGA | 424 | J2 |
| *YKL062W 2* | CGTTTCTCAAGCTCTGAGCGGTTATAGT | TGCCGACATTGTGGTTGCATCTAAGCTA | TGTCAGCCAAGCCTTATCAGGCTACTCA | AGCGCTCATGGTAGTAGCGTCCAATGAG | 286 | J2 |
| *YKL060C 1* | AGCAGCAGCAATGGAGAAGTTTGGAGAG | AGTTGTCTTACACTCTGACCACTGTGCC | GGCGGCGGCGATACTAAAATTAGGGCTA | TGTCGTTTTGCATAGCGATCATTGCGCT | 355 | J3 |
| *YKL059C 1* | ACCGCCGTTATTTACAGAAGTTGGAAGG | AGGTGGTCTTTTGAGGCAGCCGGTAAAG | GCCACCATTGTTAACGCTGGTAGGCAAA | CGGCGGCTTGTTAAGACAGCCAGTTAAA | 322 | J3 |
| *YKL057C 1* | TGTTCTTACTGTGTCGGAAAGACACTCG | TCCGGAGGCTATCAATGATAAATTGCCG | GGTACGAACGGTATCACTCAAGCACTCA | CCCAGAGGCCATTAACGACAAGTTACCA | 361 | J3 |
| *YKL057C 2* | TACGGTAGTCATAGAATCAGGCAGCTCT | CGTTACAAAGACTGGAGATGTTGAGAGG | AACAGTGGTCATGCTGTCTGGTAACTCA | TGTCACCAAAACCGGTGACGTCGAGAGA | 451 | J3 |
| *YKL057C 3* | TAGTAAAGGCTCGTAATGTACGCCGTCA | CATCTCTACACCAGTTCCTCAGCCTTAC | CAACAATGGCTCATAGTGAACACCATCG | AATTAGCACCCCTGTCCCACAGCCATAT | 466 | J3/J4 |
| *YKL056C 1* | AACAAATGGAGTGGTACCATCTTCACGG | TGTCGGAGGTGACAACATCGATATCGGT | GACGAAAGGGGTAGTGCCGTCTTCTCTA | CGTTGGTGGCGATAATATTGACATTGGC | 358 | J4 |
| *YKL055C 1* | TGTAAATCTGGACAGTGCGGCTTTAGAG | CTGGCTTGACTATGAGTCATATGACGGT | GGTGAAACGACTTAAAGCAGCCTTGCTA | TTGGTTGGATTACGAGAGCTACGATGGC | 412 | J4 |
| *YKL054C 2* | CTTCTGTTGAGAGTCTTCAGAATGCTGG | TCAACAACAACAGCCATACGGTGGCTCA | TTTCTGTTGGCTATCTTCGCTGTGCTGA | ACAACAACAACAGCCTTATGGCGGTAGC | 217 | J4 |
| *YKL054C 1* | AAGAGGAGACTCGGTCTTTTTAACAGCC | GGTACAAACACCACGAGCTCATACAACC | CAATGGGCTCTCAGTTTTCTTGACGGCT | AGTTCAAACCCCTAGAGCCCACACCACT | 397 | J4 |
| *YKL052C 1 ** | AGAGTCCGGAAGTTTCAATATCTGTCCC | GAATTTGCAGGCAAATGCAGCTGCACCA | GCTATCTGGCAACTTTAAGATCTGACCG | TAACTTACAGGCTAACGCTGCCGCTCCT | 256 | J4/J5 |
| *YKL051W 1* | TCTCGGCAGCATTGGTGAACTAGCCCTT | TTTATACCAAGGCAGCTCATCCACAGTG | CTTGGGTTCAATCGGCGAATTGGCTTTG | CTTGTACCATGGTAACTCGTCAACGGTA | 262 | J5 |
| *YKL050C 3* | GTTGGAATCGCGAACTCTTATCTCCGAT | GTTGGATGATGGGCCTAAAACACCAACC | ATTACTGTCTCTGACACGGATCTCGCTA | TTTAGACGACGGTCCAAAGACCCCTACT | 427 | J5 |

| Amplicon | WT_F | WT_R | syn_F | syn_R | bp | chunk |
|---|---|---|---|---|---|---|
| *YKL050C 1* | AGTGATTGACCTAG CACTGGCAGAACTA | GGCTAAGAGGTCTA GAAGGGGCACT | GGTAATGCTTCTGG CTGAAGCGCTTGAG | CGCCAAAAGAAGCC GTAGAGGTACC | 208 | J5 |
| *YKL050C 2* | CGCAAGATCAACTG ACTCACTCTTGGAG | GCTACATTCTTCTGA AATGCATCGCGCT | AGCCAAGTCGACGC TCTCTGATTTACTA | TTTGCACAGCAGCG AAATGCACAGAGCC | 226 | J5 |
| *YKL049C 1* | GGCATATGAATGAG TTCGCACTGGTGCA | CGATCGTGCGTTAT CGTTATTGCAGAGA | AGCGTAGCTGTGGG TTCTAACAGGAGCT | TGACAGAGCTTTGA GCTTGTTACAGCGT | 217 | J5 |
| *YKL048C 2* | TAGTGGGGAACTCG GTTTGTCCTGTGAG | TTCGCCAGACTCGA GCGATTATTGTTCA | CAAAGGACTTGATG GCTTATCCTGGCTA | AAGCCCTGATAGCT CAGACTACTGCAGC | 286 | J5 |
| *YKL048C 1* | CCCTAACGATGAAC AATAATCGCTCGAG | TTTGTCGCGTGACC AGCCCATAGATTCT | ACCCAAGCTGCTGC AGTAGTCTGAGCTA | CTTAAGCAGAGATC AGCCAATCGACAGC | 286 | J5 |
| *YKL047W 1* | TGCGTATGCTGCCA ACAGATACTCAAGC | AATTTCCTCGCTGGT CGTAGCCACATAA | CGCTTACGCCGCTA ATCGTTATAGCTCA | GATTTCCTCTGAAGT GGTGGCAACGTAG | 478 | K1 |
| *YKL047W 2* | CAAAGCAACTCCTC TTGACTTAGAGGTG | TGAATTGTTGGGTTC TGGGCCAGAAACG | TAAGGCTACCCCATT GGATTTGGAGGTT | GCTGTTATTTGGTTC AGGACCGCTGACA | 241 | K1 |
| *YKL046C 1* | TCTTGTGTGCCAGA GGTCTGATCCC | TCCCGAGAATGAGC CTCAATGGCTATAT | ACGGGTATGCCACA AATCGCTACCG | ACCAGAGAACGAGC CACAATGGTTGTAC | 415 | K1 |
| *YKL045W 1* | TCATCTTTCGTCGG GGTACACTATCGCG | AGATAGTCTCTCGT GACTCCAATCACGG | CCACTTGAGCAGCG GTTATACCATTGCT | GCTCAAACGCTCAT GTGACCAGTCTCTA | 460 | K1 |
| *YKL043W 1* | CAGTTCAACGTCAG TTCTCAAACCACGC | AGGAATCCAAACAC CCTTGAGGTGCATG | TTCAAGCACCAGCG TCTTGAAGCCTAGA | TGGGATCCAGACGC CTTTCAAATGCATT | 250 | K2 |
| *YKL042W 1* | TAGAGACTATAATG ACGTGGGGTCACGT | GAGGACCGAGAGG GTTCTTTCAATGGCT | CCGTGATTACAACG ATGTTGGTAGCAGA | CAAAACGCTCAAAG TACGTTCGATAGCC | 487 | K2 |
| *YKL041W 1* | CACTTCCAGAGCGC AGTTGGATTCTGTT | AACAGCTTCGTCCAT CTCTTCCCCTACG | TACCAGTCGTGCTC AGTTAGACAGCGTC | GACGGCTTCATCCA TCTCTTCACCAACA | 253 | K2 |
| *YKL040C 1* | GTCCTCCAAGATGG CTGGTCTTATTCTT | TTTGCAGTGTCCAG GAGTGGAGTCTTTA | ATCCTCTAAAATAGC AGGACGGATACGG | CTTACAGTGCCCTG GTGTTGAGAGCTTG | 298 | K2 |
| *YKL039W 2* | TGGTCTTTTGGCGG TCGCATACGTTGTG | TGGAGATGGTGCCT CCATGTCGGTCAAA | CGGCTTGTTAGCTG TTGCTTATGTCGTT | AGGGCTAGGAGCCT CCATATCAGTTAAG | 403 | K2 |
| *YKL039W 1* | TGCGTCCTTTTTGA GTGTACTAGCTGGT | ACGCTCACCAGTGA TGATACTTGCGTCA | CGCTAGTTTCTTATC AGTTTTGGCCGGC | TCTCTCGCCGGTAA TAATTGAAGCATCG | 307 | K2 |
| *YKL038W 1* | CAGTAACACAGTTC CAAGTACCCCTTCG | GGTGACACTTCCGT TTGCATTTCCAGAA | TTCAAATACCGTCCC TTCAACTCCAAGC | AGTAACTGAACCATT AGCGTTACCGCTG | 337 | K3 |
| *YKL038W 2* | CCCATCGATATCAT CTTTTGGCCAGTTC | CGTGGATAATGTTAC TGATACGGGGGCT | TCCTAGCATCAGCA GCTTCGGTCAGTTT | GGTACTCAAGGTAA CGCTAACTGGAGCA | 241 | K3 |
| *YKL038W 3* | TTTGGAAGCATCCA GCCCTGGATCAACG | ACCTGGCTTTTGTTT CAGAGCAACACGA | CTTAGAAGCTAGTTC ACCAGGTAGCACC | GCCAGGTTTTTGCTT TAAGGCGACTCTG | 379 | K3 |

| Amplicon | WT_F | WT_R | syn_F | syn_R | bp | chunk |
|---|---|---|---|---|---|---|
| *YKL038W 4* | TATTGTTGGCGTGG CTTTGTCGCCAAGT | AGTGCTGACCTCTT CATCAACTGGAGGC | CATCGTCGGTGTTG CCTTAAGCCCTTCA | GGTTGAAACCTCTTC GTCGACAGGTGGT | 415 | K3 |
| *YKL035W 1* | CTCTAGTAAACTTGA CGATGCTGCTCGC | AACAGATTTAGGGC CAACGCAGCCCATG | TAGCTCAAAGTTGG ATGACGCCGCCAGA | GACGCTCTTTGGAC CGACACAACCCATT | 268 | K3 |
| *YKL034W 1* | TGGATTACTGTATTC TCCAGACTGTGGC | GAAAAACCTACCATA TAACGAGCCACCG | AGGTTTGTTATACAG CCCTGATTGCGGT | AAAGAATCTGCCGT ACAAGCTACCGCCA | 499 | K3 |
| *YKL034W 2* | CCGTAACGCCGTTA AAGGTATTCCTTCA | AGGCGTTACCATGT AAGAGTGCTGATCC | TAGAAATGCTGTCAA GGGCATCCCAAGC | TGGGGTAACCATAT AGCTATGCTGGTCA | 463 | K3 |
| *YKL033W-A 1* | TACTTGCGAATTTCT TCCCGGTGCATTG | TGCTTCTGGATGGG GGACCCAAATAACG | CACCTGTGAATTCTT GCCAGGCGCTTTA | AGCTTCAGGGTGTG GAACCCAGATGACA | 355 | K3/ K4 |
| *YKL033W 1* | CGCTCTAGGACATT CTGTTACCATTCTG | TTTTGCCAGAACCTC TAGCGTGCGAACA | AGCCTTGGGTCACA GCGTCACTATCTTA | CTTAGCTAAGACCTC CAAGGTTCTGACG | 241 | K4 |
| *YKL033W 3* | TCTTCCGGGGAACG TTTCTGTTTTCGCT | CGGATCACGCTCCA AATGTACTAAAGTG | CTTGCCAGGTAATG TCAGCGTCTTTGCC | TGGGTCTCTCTCTAA GTGAACCAAGGTT | 466 | K4 |
| *YKL033W 2* | CGACTCCAAATTAA GAGAGCCAGATAGC | ACATGCGGGCTGAA CGATCGAATAATCA | TGATAGTAAGTTGC GTGAGCCTGACTCA | GCAAGCTGGCTGGA CAATGCTGTAGTCT | 331 | K4 |
| *YKL032C 2* | GAAATAAGCGGAAG AAGGTCTCTTTGGG | TGCAGGAAATGCTG CAGGAAATGCCAAC | AAAGTAGGCACTGC TTGGACGTTTAGGA | AGCTGGTAACGCCG CTGGTAACGCTAAT | 331 | K4 |
| *YKL032C 1* | ACTAGACGCGCCTG TACCCGTG | GCCTTATCAAGGTC ACTTCCAGCAGTCG | TGAGCTAGCACCGG TGCCGGTA | TCCATACCAAGGCC ATTTTCAGCAGAGC | 337 | K4 |
| *YKL029C 1* | AGACCCGGCACCGT AAATAAGCACTCTG | TGACCAAGGTATCG GTGGTGTACGTATT | GCTACCAGCGCCAT AGATCAAAACACGA | CGATCAAGGCATTG GCGGCGTTAGAATC | 430 | K4 |
| *YKL029C 2* | AATTCCCAGAATAC CTTCCGAATCCGAC | TGCCACTCGTTTGA CTGTAGAAGGTGCC | GATACCTAAGATGC CTTCGCTGTCGCTA | CGCTACCAGATTAA CCGTTGAAGGCGCT | 484 | K4 |
| *YKL028W 1* | CAGTAGTAGACGTG CCGGTGCTAACTCT | CTTTGCTTGCTTTTT AGCCAAAGCAGCG | TTCATCACGTAGAG CTGGCGCCAATAGC | TTTAGCTTGTTTCTT GGCTAAGGCGGCA | 370 | K5 |
| *YKL027W 1* | CAAAGTGATCTCTT CCATGGGTGCTTCC | GCTCGGTATACCGA CTCTGCTCATTTGA | TAAGGTTATTAGCAG TATGGGCGCCAGT | TGATGGGATGCCAA CACGTGACATTTGG | 442 | K5 |
| *YKL026C 1 ** | AAGCTCTAGTGGCC TAGTCATGCATGAA | GGCCTTTCCCTGTG GTCAATTCGGAAAT | CAACTCCAAAGGTC TGGTCATACAGCTG | TGCTTTCCCATGCG GCCAATTTGGTAAC | 274 | K5 |
| *YKL025C 2* | ACCTGAAGAGGGGA AAACCTGCAAAATC | TCCTATGCAACCGC CACCCATAGAGAGT | GCCGCTGCTTGGAA AGACCTGTAAGATA | ACCAATGCAACCAC CTCCAATCGAGTCA | 232 | L1 |
| *YKL025C 1* | AGTGTGAATTGAGG AGGTTGAAGACGGA | CGCAAGTTTCACTC CGATGACAGTCGGT | GGTATGGATGCTAC TAGTGCTGCTTGGG | AGCTTCATTTACCCC AATGACCGTTGGC | 286 | L1 |
| *YKL024C 1* | GGACTCAATGTTGT CATCGCTTCTACCA | CAGAGCAGGTTCCC AATATGGGGAATTG | ACTCTCGATATTATC GTCTGAACGGCCG | TCGTGCTGGCAGTC AATACGGTGAATTA | 298 | L1 |

| Amplicon | WT_F | WT_R | syn_F | syn_R | bp | chunk |
|---|---|---|---|---|---|---|
| *YKL023W 1* | GGCCCATGAGGAAA GCTCTTCACAAAGT | ATTTGGCGACGGAC TTGCTCCGTTGACA | CGCTCACGAGGAAT CAAGCAGCCAATCA | GTTAGGGCTTGGTG AAGCACCATTAACG | 292 | L1 |
| *YKL022C 1* | AAGTGGGTGGCTGG CATCCAATGATAAC | TCTCGGGATGCAGT TTATGGCGATGAAT | CAAAGGATGTGAAG CGTCTAAGCTCAAG | CTTGGGTATGCAGT TCATGGCTATGAAC | 487 | L1 |
| *YKL022C 2* | TCCAGGAAAGAATC TGGAGGCTGTAGAG | TGCGATAACATGGTT TAGCGTTGCGACC | ACCTGGGAAAAAAC GACTAGCGGTGCTA | AGCTATCACCTGGTT CTCAGTCGCTACT | 202 | L1 |
| *YKL022C 3* | GGACGGTATGGTAG AACCGAACATACTA | CAGAACCAATACGG CTACGAGTCCCTAC | ACTTGGGATAGTGC TGCCAAACATTGAG | TCGTACTAACACCG CCACCTCACCATAT | 205 | L1 |
| *YKL021C 1* | GCTATTCTTGCTCC CCTTTGAAACAGCA | TACCTCATCTCGAAA AGATCCTAAGGGC | TGAGTTTTTTGAACC TTTGCTGACGGCG | CACTAGCAGCAGAA AGGACCCAAAAGGT | 409 | L1 |
| *YKL020C 3* | AGAGCTATTCTGTG AGTGTAATCCTGCG | TATTCAGCGTGTCAT CCCCGCACAAGGT | GCTTGAGTTCTGGC TATGCAAACCAGCA | AATCCAGAGAGTTAT TCCAGCTCAAGGC | 478 | L1 |
| *YKL020C 2* | GTCCTCACTGGAGG AATTTGGTGATGGA | CAATGCCCAGGTTT CGCCAATGACAAAT | ATCCTCTGAACTACT GTTAGGGCTAGGG | TAACGCTCAGGTCA GCCCTATGACCAAC | 250 | L2 |
| *YKL020C 1* | GTCACTTTGCGGCG ATGAAAAGGATCTA | CAACAGGGAAAAAA GAAGAGCGTCGAGA | ATCTGATTGTGGGC TGCTGAAACTACGG | TAATAGAGAAAAGC GTCGTGCTAGCCGT | 397 | L2 |
| *YKL019W 1* | GCTACATCCGTCTC CTTCCTTCAAAAGA | GCCAATTGGAAGTG ACAATACGTCACCG | ATTGCACCCAAGCC CAAGTTTTAAGCGT | ACCGATAGGCAAGC TTAAAACATCGCCA | 418 | L2 |
| *YKL018W 1* | TGGCCAGTTTCTAC TGACCTCTTCTTCC | TGACGACGATAGAA ACGTATCGTTCACG | CGGTCAGTTCTTGTT AACTAGCAGCAGT | GCTGCTGCTCAAGA AGGTGTCATTAACT | 283 | L2 |
| *YKL017C 2* | ACGGCCCTGGAACC CATCTACAGTTGAT | TGGGAAACTATTGG CCGATGCAACGGTC | TCTACCCTGAAAACC GTCAACGGTGCTG | CGGTAAGTTGTTAG CTGACGCTACCGTT | 364 | L2 |
| *YKL017C 1* | CGATGATGAACCGT GTAAAGTGGTGACG | TTCTGTGGATACGAT TCTGGAGAGGCTA | GCTGCTGCTGCCAT GCAAGGTAGTAACA | CAGCGTTGACACCA TCTTAGAGAGATTG | 331 | L2 |
| *YKL016C 1* | TCTGTCCTTGTAAC CAGGTACGTCCCAT | TGAGGCACGTAGAC AATTACTAGAGCTG | ACGATCTTTATAGCC TGGAACATCCCAC | CGAGGCTAGACGTC AATTGTTGGAGTTA | 388 | L2 |
| *YKL015W 1* | AGCTTTGCTCTTGG AGAGACCAGTGAGT | ATGACCATTGGCAC CGTTGCCATTTTCG | GGCCTTATTGTTAGA GCGTCCTGTTTCA | GTGGCCGTTAGCGC CATTACCGTTTTCA | 310 | L2/ L3 |
| *YKL015W 3* | TGGTAGTAGTGGCC AGGAAGAGGTAATA | GGCTAAAGTGTAAT CAGCCACCTGTAAG | CGGCTCATCAGGTC AGGAAGAGGTTATC | AGCCAAGGTATAGT CGGCAACCTGCAAA | 475 | L3 |
| *YKL015W 2* | CAAAGATGGCGATC GCATTGACCCTTCA | ATTTCCCAGCCACAT TGCTTGATCAGTG | TAAGGACGGTGACA GAATCGATCCAAGC | GTTACCTAACCACAT AGCTTGGTCGGTA | 415 | L3 |
| *YKL014C 3* | CAAACCACCATCAT CTGCTACCCTTTGG | TGTTGATCTATTGGG AAGTCCAACTGCG | TAAGCCGCCGTCGT CAGCAACTCTTTGA | CGTCGACTTGTTAG GTTCACCTACCGCT | 370 | L3 |
| *YKL014C 2* | GGCAACGGTATGAC AATCGCTTAATGAG | CTACGATGATACCG AACGATCTGGTGTT | AGCGACAGTGTGGC AGTCTGACAAGCTA | TTATGACGACACTGA AAGAAGCGGCGTC | 202 | L3 |

| Amplicon | WT_F | WT_R | syn_F | syn_R | bp | chunk |
|---|---|---|---|---|---|---|
| *YKL014C 4* | ATGGCCTGTTTCAAAGCATACTGCACCG | TTGGTTCGAAGCAACTGCCTGTGAGTTA | GTGACCGGTTTCGAAACAAACAGCGCCA | CTGGTTTGAAGCTACCGCTTGCGAGTTG | 208 | L3 |
| *YKL014C 1* | TGCTACGCTAAAGTCACGGGAAGCTCCA | ACCAAGCAAGTCTGAGTTAGCCAATGGG | AGCAACTGAGAAATCTCTACTGGCACCG | TCCTTCAAAAAGCGAGTTGGCTAACGGT | 400 | L4 |
| *YKL012W 2* | AACAAGAGAAACCAGCTGGACTATTCCG | CCTTGGATCTCTGGTCCCCAGTTCTGAA | CACCCGTGAAACTTCATGGACCATCCCA | TCTAGGGTCACGAGTACCTAATTCGCTG | 319 | L4 |
| *YKL012W 1* | CAAGCCTAGTACGTGGGATTTAGCTTCC | CAATTCCACAGCCGGAGTTAAATGCCTC | AAAACCATCAACCTGGGACTTGGCCAGT | TAATTCAACGGCTGGGGTCAAGTGTCTT | 202 | L4 |
| *YKL011C 1* | CAAATCGTCATCTTTTCTCACCCCCGAA | CATTCCAAGAGAAGAGACACCGACCAGT | TAAGTCATCGTCCTTACGAACACCGCTG | TATCCCTCGTGAAGAGACCCCAACTTCA | 262 | L4 |
| *YKL010C 4* | AATGGCTTGACAAAGACGAGACCGCATA | TTTCGGACGAGTTGAGGAAGACTGGTCT | GATAGCTTGGCACAATCTGCTTCTCATG | CTTTGGTAGAGTCGAGGAAGATTGGAGC | 349 | L5 |
| *YKL010C 1* | TCTAGCGAGCTTAGCGCTTAGTTTTGAG | CCTTACATCTTCCAGTACCGAAGCTGAT | ACGGGCCAATTTGGCTGACAACTTGCTA | TTTGACCAGCAGTTCAACTGAAGCCGAC | 415 | L5 |
| *YKL010C 3 ** | AAGCAACGAGAGACCACCAACCAACAGT | TCCTCCCGAGGATGAAAGAATACTGTCA | CAATAAGCTCAAGCCGCCGACTAATAAG | CCCACCAGAGGACGAACGTATCTTAAGC | 343 | L5 |
| *YKL010C 2* | ACTGCTACAGGCGTTCGAAACAATTGCG | GAATGCCTCCGAGGATCCTTATATTGCA | TGATGAGCAAGCATTGCTGACGATAGCA | TAACGCTAGTGAGGACCCATACATCGCT | 457 | L5 |
| *YKL009W 1* | CGTTCGTTCAGACTATTCAAGACCGAAC | GACTTTGAATTCAGAAGCAGCGATACCG | TGTCAGAAGCGATTACAGCCGTCCAAAT | AACCTTAAATTCGCTGGCGGCAATGCCA | 289 | L5 |
| *YKL008C 1* | TATCCCGAAAGGACCGGAAACACCG | TTACCGTCATGCCTGGATAGCTCCCTTG | GATACCAAATGGGCCACTGACGCCA | CTATAGACACGCTTGGATCGCCCCATTA | 337 | L5 |
| *YKL007W 1* | TCTTCTTCAGAGCTTTCCAGGTGATGTC | ATAACTGCCAATCGCACTACCCCAGTTA | CTTGTTGCAGTCATTCCCTGGCGACGTT | GTATGAACCGATAGCTGAGCCCCAATTG | 412 | M1 |
| *YKL005C 1* | TGTACTTGGGCTGGGTTCTTTCGAAATG | CAAAGCTTGCCTTAACGTCGAGTTTGGT | GGTTGAAGGTGATGGTTCCTTGCTGATA | TAAGGCCTGTTTGAATGTTGAGTTCGGC | 427 | M1 |
| *YKL005C 2* | AGTATCAGACTTGGCGGCATCAATGCTT | TTTGGAAACGTCAAAGGAGGCAGAGGTT | GGTGTCGCTTTTAGCAGCGTCGATTGAC | CTTAGAAACCAGCAAAGAGGCTGAGGTC | 241 | M1 |
| *YKL004W 1* | CACTTCATCGTACTTTCCAGATGACCGC | AAACAGGGCTTCCATAGTAGCACACCCG | TACCAGCAGCTATTTCCCTGACGATAGA | GAATAAAGCTTCCATGGTGGCGCAACCT | 487 | M1 |
| *YKL001C 1* | GTCGGTTCTCAAATGTAGCTCTGGAGCT | GTCAGGTAAAAGTACAATCGCCTGTGCG | ATCAGTACGTAAGTGCAACTCAGGGGCC | TAGCGGCAAGTCAACCATTGCTTGCGCT | 430 | M2 |
| *YKR001C 2* | AACTGGCGCCTTACTAGACGATGGTTGG | TGGTGTAGACTCTTTGGATCCATTCGAC | GACAGGAGCTTTTGAGCTGCTAGGTTGA | CGGCGTTGATAGCTTAGACCCTTTTGAT | 490 | M2 |
| *YKR001C 1* | CAAAGATGGGGCAGAACCTGAACTATTG | CTTGGGCCCAGAAACTATGGATTCAGCT | TAAGCTAGGAGCGCTGCCGCTTGAGTTA | TTTAGGTCCTGAAACCATGGACAGCGCC | 259 | M2 |

| Amplicon | WT_F | WT_R | syn_F | syn_R | bp | chunk |
|---|---|---|---|---|---|---|
| *YKR002W 1* | TACGTATGGGTCCTATAGACTAGGAGTC | AGCCCTTCTTTGGGCCCATAGCTTAATA | CACCTACGGTAGTTACCGTTTGGGTGTT | GGCTCTACGTTGAGCCCACAATTTGATG | 412 | M2 |
| *YKR002W 2* | CAGGTCTCATAGAATGCCCGTCATTACA | GGCGTTCAGTGCTGTCTCAGTTTTATGA | TAGAAGCCACCGTATGCCAGTTATCACC | AGCATTTAAAGCGGTCTCGGTCTTGTGG | 433 | M2 |
| *YKR003W 1* | TGAATCTTCGATTAGAGTGGATGGGGTC | AGTCCCTTCGATTGCATCGTAAGTTCCA | AGAAAGCAGCATCCGTGTTGACGGTGTT | GGTACCTTCAATAGCGTCATAGGTACCG | 301 | M3 |
| *YKR006C 1* | GAGTTTAGGATCGTTGGCAGATGGCAAA | TATTGAAGAAGGCACTAACGAGGCTTCC | CAACTTTGGGTCATTAGCGCTAGGTAAG | CATCGAAGAAGGTACCAATGAGGCCAGT | 415 | M3 |
| *YKR007W 1* | TCAAGGGACAGACCTGCAAGAAGCTCTT | TTTCGCTGAATCCGGTGATGTAAGCAGA | CCAAGGTACCGATTTACAAGAAGCCTTG | CTTAGCGCTGTCTGGGCTGGTCAATAAG | 202 | M3 |
| *YKR008W 1* | TCTGCTGGGTGCGTCATCTCCTAAAAAC | CGCACCTGCTGCGGCTGGA | CTTATTAGGCGCTAGCAGCCCAAAGAAT | AGCGCCAGCAGCAGCAGGG | 415 | M3 |
| *YKR008W 2* | CGAAGGTCTCGGCAATGGTTATAATCGT | TGGCCTTGAACCCAGTAAAGGCTTAGTT | TGAAGGCTTGGGTAACGGCTACAACAGA | AGGTCTGCTGCCTAACAATGGTTTGGTA | 343 | M3 |
| *YKR009C 1* | TGCCATAGAAGACTCCTCTGTGGAGCTC | TGCATCTCAAGTCTCCCCACTTGTTGTT | AGCCATGCTGCTCTCCTCGGTACTTGAA | CGCTAGCCAAGTTAGTCCTTTGGTCGTC | 256 | M4 |
| *YKR009C 3* | GATTGCATGAGACTTCCCAAGACCACCT | CAATCCAGACCCCAAGACATATACTCCT | AATAGCGTGGCTTTTACCCAAGCCGCCA | TAACCCTGATCCAAAAACCTACACCCCA | 247 | M4 |
| *YKR009C 2* * | ACGCGATAGCTTATAGCCACCTGTCAAA | CGTCAATGATCTAGGTGGCACTTTGGGT | TCTGCTCAATTTGTAACCGCCGGTTAAG | TGTTAACGACTTGGGCGGTACCTTAGGC | 289 | M4 |
| *YKR010C 1* | AGGTGAAGAATGCGACCTGGTAACCTTG | GAAACCGCCAAATGCAGCTTCGACC | TGGGCTGCTGTGGCTTCTAGTGACTTTA | TAAGCCACCTAACGCTGCCAGCACT | 247 | M4 |
| *YKR010C 2* | TAAGGAACTCTTCCTGGGACTATGAGCA | TGATGGCGCTAAAAGGGCCGAATCAAAG | CAAACTTGATTTTCTTGGTGAGTGGGCG | CGACGGTGCCAAGAGAGCTGAAAGCAAA | 220 | M4 |
| *YKR011C 1* * | AGGTCTTTGTCTCGACCAGGCTCTCAAG | ATCGTGCTCAAACAATGAGCCTTCCGCA | TGGACGTTGACGGCTCCAAGCACGTAAA | TAGCTGTAGCAATAACGAGCCAAGTGCT | 274 | M4/M5 |
| *YKR013W 1* | TCAATCTACCGCCTCAAGCACTCAATCC | GAAGTGACCTGCGCTTTCACTAAAACCA | CCAAAGCACTGCTAGCTCAACCCAAAGT | AAAATGGCCAGCTGATTCTGAGAAGCCT | 310 | M5 |
| *YKR014C 1* | CTTAACCCCTTCACCTGTCTTAGCACTA | GCAGGAAGATTCACTACAAAAGGCAAGG | TTTGACACCTTCGCCGGTTTTGGCTGAG | CCAGGAAGACAGCTTGCAAAAAGCTAGA | 268 | M5 |
| *YKR015C 1* | AGTTGACCCAAGTACTTCACTCTCCCCA | TCAGGTCGAAGAACCTGGTAAGCACTCG | GGTGCTACCCAAAACTTCTGACTCACCT | CCAGGTTGAAGAACCAGGCAAACATAGC | 313 | M5 |
| *YKR015C 2* * | ATCACACGCTGCTGTAGCGTTGAATCTG | CTTAGTCAGGCGTAGCCCTGCTGTTTTA | GTCGCAAGCAGCGGTGGCATTAAAACGA | ATTGGTTAGAAGATCACCAGCCGTCTTG | 400 | M5 |
| *YKR016W 2* | TACCCTTTGGACTATTGCTTTGTCAGCG | GCGCGTTTGTGAACTACCAGTTAATCCA | CACTTTGTGGACCATCGCCTTAAGCGCT | TCTGGTTTGGCTTGAGCCGGTCAAACCT | 241 | N1 |

| Amplicon | WT_F | WT_R | syn_F | syn_R | bp | chunk |
|---|---|---|---|---|---|---|
| *YKR016W 1* | AAACCCTCCATCGC TATTAAGCGTAGCT | TTTATGAGGCCAAC CCTTTAGGGAGACA | CAATCCACCTAGCTT GTTGTCAGTTGCC | CTTGTGTGGCCAGC CTTTCAAACTAACG | 328 | N1 |
| *YKR017C 2* | ATGAGATGCATCCG AGTAATATGCCACG | CGAATTCTGTTGGAT TTGTGAGGGCCCC | GTGGCTAGCGTCGC TATAGTAAGCAACA | TGAATTTTGCTGGAT CTGCGAGGGTCCA | 352 | N1 |
| *YKR017C 1* * | CTTACTACTCGAAG GATGTCCCATCACT | GGAGCTTGGCTTAA GTACCACAGCAAAT | TTTTGATGAGCTTGG GTGACCCATAACC | TGAGTTGGGTTTGT CAACTACCGCTAAC | 286 | N1 |
| *YKR018C 1* | TAGCGCACTATTGG GAAATAGCGCTCTT | CAGGGGTTCCAGAG AAGAAGGTTTGAGA | CAAAGCTGAGTTTG GGAACAAAGCACGA | TAGAGGCAGTCGTG AAGAAGGCTTACGT | 277 | N1 |
| *YKR018C 2* | GGTGGACTCCTTAT CTTCACTGCCTGAC | CGAGGGTTTGGCTT TGCTGGATGAGAGT | AGTACTCTCTTTGTC TTCTGAACCGCTG | TGAGGGCTTAGCCT TATTAGACGAGTCA | 436 | N1 |
| *YKR019C 2* | TGAAGTGTCCGGAT TGAGACTATGTCTG | TGAATCACTGACGC CTGGCCAAAGAACT | GCTGGTATCTGGGT TCAATGAGTGACGT | CGAAAGCTTAACCC CAGGTCAACGTACC | 259 | N2 |
| *YKR019C 1* | TGTATCCAATGCCG AAGAATGGGAGGAG | TTCCAATGAGGTCA GCAGTCCATGCCCT | GGTGTCTAAAGCGC TGCTGTGACTACTA | CAGTAACGAGGTTT CATCACCTTGTCCA | 310 | N2 |
| *YKR021W 1* | TAATTCGTCCAATC GCCCATCTATGAGC | ATTTGCCGGTACCG ATCCTCTTCGACTA | CAACAGCAGTAACA GACCTAGCATGTCA | GTTAGCTGGAACGC TACCACGTCTTGAT | 265 | N2 |
| *YKR021W 3* | GCCAGTATCAAGTG ATGTTTCCCATGAC | TACGGATCCATCAG AAGGAAGCCTTACG | ACCTGTTAGCTCAG ACGTCAGTCACGAT | AACACTACCGTCGC TTGGCAATCTAACA | 433 | N2 |
| *YKR021W 2* | TCTTTCGTCTCCCG TTCTTTCACCTAAC | ACCATTGAATGAGTC CTCCTCAATGCTG | CTTGAGCAGCCCAG TCTTGAGCCCAAAT | GCCGTTAAAGCTAT CCTCCTCGATTGAA | 490 | N2 |
| *YKR022C 1* | ACTACTAATGTGCT CACCTTCGGAGCTA | GCCCCTAGTGTCGA ACGATTCTGAAGAT | TGATGAGATATGCTC GCCTTCACTTGAG | ACCATTGGTTAGCAA TGACAGCGAAGAC | 295 | N2 |
| *YKR023W 2* | AAGCTCGACAACGA GGCCTACCGTT | CGCGATGTCGAAAA CTGGATGCCTTGTA | GTCAAGCACCACCA GACCAACTGTC | AGCAATATCAAAGAC AGGGTGTCTGGTG | 208 | N2 |
| *YKR023W 1* | GTCCTTATCCAATG CGTCAAATGGAGCT | GTTTTCCTCAGCTTT ATGGTCATGCTCG | TAGTTTGAGTAACGC TAGCAACGGTGCC | ATTTTCCTCGGCCTT GTGATCGTGCTCA | 241 | N2 |
| *YKR024C 1* | AACGTGGGGAAGGT CTAGGCCTCTACTT | TGACGGGCACCATA AAAATCTAACGGGT | GACATGTGGCAAAT CCAAACCACGTGAA | CGATGGTCATCACA AGAACTTGACCGGC | 250 | N3 |
| *YKR024C 2* | AAACTCCACGCTGT CTGAACATGATACG | CATAATCGGTACAC CAGGTCGTGTTTTG | GAACTCAACTGAATC GCTGCAGCTAACA | TATCATTGGCACCC CTGGCAGAGTCTTA | 499 | N3 |
| *YKR025W 1* | CTCTGCAGCAAGAT ATAAACCCAAGTCG | TCTGGCGTCCTCTT GCTTTCTTGTTACA | TAGCGCTGCTCGTT ACAAGCCAAAAAGC | ACGAGCATCCTCTT GTTTACGGGTAACG | 277 | N3 |
| *YKR026C 1* | AACGGCACTTGGAG TGAGGACCCCTAAA | GACTCTTATTGTCGA TAGCGCGGTTGGA | GACAGCTGAAGGGG TCAAAACACCCAAG | TACCTTGATCGTTGA CTCAGCTGTCGGT | 358 | N3 |
| *YKR027W 1* | CGCCTTACTTCAGC TAAGATCTGACGTT | TCGTAGACACGCTA CTGCATCGGGTAAG | TGCTTTGTTGCAGTT GCGTAGCGATGTC | TCTCAAGCAAGCAA CAGCGTCTGGCAAA | 274 | N3 |

| Amplicon | WT_F | WT_R | syn_F | syn_R | bp | chunk |
|---|---|---|---|---|---|---|
| *YKR028W 1* | CTCTGGGGACGAAA GAAGTGTTGATTCT | GGGAGAGTCAGGTT TATCCGTCGAG | TAGCGGTGATGAAC GTTCAGTCGACAGC | TGGGCTATCTGGCT TGTCGGTGCTA | 388 | N4 |
| *YKR028W 2* | TTCCACTCAGTCTG CGGCAGGC | AGTGGGAGGGTGAC TTTCTAGGGTAGTG | CAGTACCCAGAGCG CTGCTGGT | GGTTGGTGGATGTG ATTCCAAAGTGGTA | 274 | N4 |
| *YKR028W 3* | ATCGGGTACTTCAT TGTTCCCTCCAGAC | TAATGAGTAGCCCT CCTCATTGCCCATA | CAGCGGCACCAGCT TATTTCCACCTGAT | CAAGCTATAACCCTC CTCGTTACCCATG | 385 | N4 |
| *YKR029C 2* | CCTAGTTGAATTGC CTGATAAGCGTGCA | AGACGCATATGGTG CAATATACTTGCCC | TCTGGTGCTGTTAC CGCTCAATCTAGCG | GGATGCTTACGGCG CTATCTATTTACCA | 394 | N4 |
| *YKR029C 1* | ATCGGAAGAATTGA CAGAAGCGCTTTCG | TGACCACTGCAACA GATGGCAGCATGCT | GTCACTGCTGTTAAC GCTGGCTGATTCA | CGATCATTGTAATCG TTGGCAGCACGCC | 358 | N4 |
| *YKR030W 1* | GCCAAAGAATTTAG ATCTCGAGACAGCC | GTCGAAACAATAAG CCCACTCTACCACA | CCCTAAAAACTTGGA CTTGGAGACCGCT | ATCAAAGCAGTAGG CCCACTCAACAACG | 391 | N4/ N5 |
| *YKR031C 3* | GGGCAGAGGCGTT CTCTTACGTGATGA T | TACCCTAAGTAAGG AGGGAGATAGTGGT | TGGTAATGGGGTAC GTTTTCTGCTGCTC | CACTTTGTCAAAAGA GGGTGACTCAGGC | 268 | N5 |
| *YKR031C 5* | AAGAGACCAGTTCC CAGCACTTCTAAGT | TGAGCCAGCAAGAG ACTTAGCCCGA | CAAGCTCCAATTACC GGCTGAACGCAAG | CGAGCCTGCTCGTG ATTTGGCTAGA | 208 | N5 |
| *YKR031C 1* | CGAAGAAAGATCCG ACACGTAAGTTAGG | AGCAGAAAATAAAC CATCCTCTGCAGCC | GCTGCTCAAGTCGC TAACATAGGTCAAA | GGCTGAAAACAAGC CTAGTAGCGCTGCT | 487 | N5 |
| *YKR031C 4* * | GGAGCTCCTTGGAA TAGAGGGTCTCCTA | CCCTCAAAGGCGTT CTTCTTCTGTAGCT | ACTTGATCTAGGGAT GCTTGGACGTCTG | TCCACAAAGAAGAA GCAGCAGCGTTGCC | 460 | N5 |
| *YKR031C 2* | GCTGGATTTCAGTC TACCAATAGCAGAG | TGTGTCACCGAACC ACTCATTAGACCAT | TGAACTCTTTAAACG GCCGATGGCGCTA | CGTTAGCCCAAATC ATAGCTTGGATCAC | 277 | N5 |
| *YKR034W 1* | GCAGCGGGGAAAAT TTTCTTTGGACCCC | TGGGTGAGATTGTA CTGAAGCTCTGGAG | CCAGAGAGGTAAGT TCAGCTTAGATCCA | AGGATGGCTTTGAA CGCTGGCACGACTA | 340 | O1 |
| *YKR036C 1* | TCTGCTCAGCTCAT CCCCATCTAGTGAA | ACCGGTGAGATTAC TAGAAGGCCATACC | ACGTGATAACTCGT CACCGTCCAAGCTG | GCCAGTTCGTTTGTT GGAAGGTCACACT | 292 | O1 |
| *YKR036C 2* | CCTAGATCCGCTAA CCAAAGCTTCAGAA | CCGAAATACCACGG CTTTCAGAATGACA | TCTGCTACCTGAGA CTAAGGCTTCGCTG | TAGAAACACTACCG CCTTTCGTATGACC | 418 | O1 |
| *YKR037C 1* | GTCGTGAGAAGAAG AAGGATGGTACTGT | TACAAGTGGCGGTG GCTTGAACGGTTTT | ATCATGGCTGCTGC TTGGGTGATACTGA | CACCTCAGGTGGCG GTTTAAATGGCTTC | 418 | O1 |
| *YKR038C 1* | TCTGGCGGCGATGA CAACTGAATGCAGA | TAAGCATCCGCTTCT GCCTAAACATGCC | ACGAGCAGCAATAA CGACGCTGTGTAAT | CAAACACCCATTGTT ACCAAAGCACGCT | 280 | O1 |
| *YKR039W 1* | CTCAAATGGCTCCG CTGTTTCTATTGAC | GAACCCACCCGAAA TAGGGAAGATAACA | AAGCAACGGTAGTG CCGTCAGCATCGAT | AAAACCGCCGCTGA TTGGAAAAATGACG | 361 | O1 |
| *YKR039W 2* | AAGTTTGATTGGTG CCTCCTCTGTGGAT | GGCGGCAACAAACG CAATAAGACCGAAT | GTCATTAATCGGCG CTAGTAGCGTTGAC | AGCAGCGACGAAAG CGATCAAGCCAAAA | 283 | O1 |

| Amplicon | WT_F | WT_R | syn_F | syn_R | bp | chunk |
|---|---|---|---|---|---|---|
| *YKR041W 1* | CTTTGACCAGGCAA AAGGTATCAGCTCT | ATGTTTGTCCGGGT GCCACCGGATTCTA | TTTCGATCAGGCTAA GGGCATTTCAAGC | GTGCTTATCTGGAT GCCATCTAATACGG | 262 | O2 |
| *YKR042W 1* | TACCTCTGCCGCCG CCTCTTCT | GCCGGAGACCTTGT TAACAGCTTGAGCA | CACTAGCGCTGCTG CTAGCAGC | ACCACTAACTTTATT GACGGCTTGGGCT | 400 | O2 |
| *YKR043C 1* | TCTGGCAATAGCGC GGGAAAGTCTCAAC | ACCATTAAGCGACG AGCAAAGAGCTAAG | ACGAGCGATGGCTC TACTCAAACGTAAA | GCCTTTGTCAGATG AGCAACGTGCCAAA | 226 | O2 |
| *YKR044W 1* | TACATCGCTGCGGC TTAATTCAAGAGGC | GCTACAGCCAATGT ATCTTTGCCCCGTG | CACCAGCTTAAGATT GAACAGCCGTGGT | TGAGCAACCGATAT AACGTTGACCGGTA | 226 | O2 |
| *YKR045C 1* | TGGTGGGATGGCCA GAACGCTAGAATTC | ACCAAGACGCGGCA CTAGGGCT | AGGAGGAATAGCTA AGACTGAGCTGTTG | GCCTCGTAGAGGTA CCAGAGCC | 370 | O2/ O3 |
| *YKR046C 1* | AGAGCCGATCTCCT TCGATGCGGTAGAA | TCGTCTGGTGAAGC AAGTTGTTGCTGCT | GCTACCAATCTCTTT GCTAGCAGTGCTG | AAGATTAGTTAAACA AGTCGTCGCCGCC | 256 | O3 |
| *YKR048C 1* | CAACAGCTTGAAAC CGGGTCTACCATCC | CATTTCAGGACAAG AGCAACCCAAGCCG | TAATAATTTAAAGCC TGGACGGCCGTCG | TATCAGCGGTCAAG AGCAACCAAAACCA | 283 | O3 |
| *YKR050W 3* | TTCCGGGTTCTTAA AATCATGCAGACCC | TGAAGCGGATTGTC GTGGGTCTTCTCTG | CAGTGGTTTTTTGAA GAGCTGTCGTCCA | GCTGGCACTTTGTC TAGGATCTTCACGT | 487 | O3 |
| *YKR050W 1* | CGCAAACTCCACGG ATTCTTTTTCGAGC | AACTACTTGGCCTG AGGTTTTAGGACTC | TGCTAATAGTACCGA CAGCTTCAGCTCA | GACAACTTGACCGC TAGTCTTTGGTGAA | 322 | O3 |
| *YKR050W 2* | CTTGGCCAAGTCGG CGCCTTCATCTTTT | GAAAATCAGCGGAT ACGGCGCAGTATCA | TTTAGCTAAAAGCGC TCCAAGCAGCTTC | AAAGATTAATGGGTA TGGAGCGGTGTCG | 490 | O3 |
| *YKR051W 1* | TTTAACACTACTACT AGGAGGCGAGAGG | ATATCCAGAGGTTC GATCTTGGTTTCCC | CTTGACCTTGTTGTT GGGTGGTGAGAGA | GTAACCGCTAGTTCT GTCTTGATTACCG | 457 | O3 |
| *YKR052C 1* | AGCAATGGTAGCAA TGGTACCACTCAGT | TCCAATAGATGCTCT GAAAACCCGAGTG | GGCGATAGTGGCGA TAGTGCCTGATAAA | CCCTATCGACGCCT TAAAGACTAGAGTT | 274 | O4 |
| *YKR053C 1* | CACCTTAGAACCAC ATGATGCGTACCTA | TATGCTGGATTTATT TAGCGGCGCCGCA | AACTTTGCTGCCGC AGCTAGCATATCTG | CATGTTAGACTTGTT CTCAGGTGCTGCT | 301 | O4 |
| *YKR054C 10* | AAGTTGTTCAGTTG AGCTAGCCACCTCT | TGAAGAGAAAGACC TTGAAGTGGTCGCT | CAATTGTTCGGTGCT TGAGGCAACCTCC | CGAAGAGAAGGATT TGGAAGTTGTTGCC | 268 | O4 |
| *YKR054C 8* | GGTACAATACACGC TCCATACTCCTGAA | TGATGAGTGGAATAT CGCGGATGTGGTA | AGTGCAGTAAACTG ACCAAACACCGCTG | CGACGAGTGGAACA TTGCTGACGTTGTT | 481 | O4 |
| *YKR054C 3* | TGGTACGCCATCGC TACTTTCAGAGAGA | TAGGGAAACAAGAG CAGCTAGGACACGG | AGGAACACCGTCTG ATGATTCGCTCAAG | AAGAGAAACCCGTG CTGCCAGAACCAGA | 220 | O4 |
| *YKR054C 4* | TGGAATATCTCCAC TGGGATCGCAAGAG | CGCTGTTCCTTTTCT TTTGGATCCAAGC | AGGGATGTCACCTG ATGGGTCACAGCTA | TGCCGTCCCATTCTT GTTAGACCCTTCA | 313 | O4 |
| *YKR054C 11* | CCCACACGCTTTGC TGGCCCTGTTAATT | GCAGGAGGCTACAG AGGAAATCAAAAAG | ACCGCAAGCCTTTG AAGCTCTATTGATG | ACAGGAGGCCACCG AGGAAATTAAGAAA | 415 | O4 |

| Amplicon | WT_F | WT_R | syn_F | syn_R | bp | chunk |
|---|---|---|---|---|---|---|
| *YKR054C 9* | CCCAGGTGACCTTGGATTTACCCCA | CTCAGCAATGATAAGTTCTCCAGCCCTT | ACCTGGGCTTCTAGGGTTAACACCG | AAGCGCTATGATCTCAAGCCCTGCTTTG | 274 | O4 |
| *YKR054C 1* | CAAGCCCGAAAAGAGAAGTGAAGTTGAG | CAATCCTCCAACCGACCCAGGTAGA | TAAACCGCTGAACAACAAGCTGGTGCTA | TAACCCACCTACTGATCCTGGCCGT | 493 | O4 |
| *YKR054C 2* | ATCATAGAGCGAAGAGTTCCGGAGTGCA | GAGATCCCTACTATATGCACTAGCTGGT | GTCGTACAAGCTGCTATTTCTCAAAGCG | ACGTAGTTTGTTGTACGCTTTGGCCGGC | 361 | O4 |
| *YKR054C 12* | CACATCCGTAGAAAACCACAGCAAACCA | TGATGGTCACGCTAACGTCGTCTATGTT | AACGTCGGTGCTGAACCATAATAAGCCG | CGACGGCCATGCCAATGTTGTTTACGTC | 376 | O5 |
| *YKR054C 5* | GTCACCGAGCGATGGTAAAATCACTCTT | TGAGAAAGTTTTGAGTGCGGTGTCTGCC | ATCGCCCAAGCTAGGCAAGATAACACGC | CGAGAAGGTCTTATCAGCTGTTAGCGCT | 463 | O5 |
| *YKR054C 6* | ACCTACTAGAAGCCTGGAAAGAACCTGA | GGAAGAAGCTCGACTACTATGGGCGAAA | GCCAACCAACAATCTACTCAAGACCTGG | AGAAGAAGCCAGATTGTTGTGGGCTAAG | 349 | O5 |
| *YKR054C 7* | AGTCAGGGTAGTGGAAAGAGGATGATCA | TTTTGACGGTACTGCTGATGATCTGCAC | GGTTAAAGTGGTACTCAATGGGTGGTCT | ATTCGATGGCACCGCCGACGACTTACAT | 352 | O5 |
| *YKR055W 1* | CGCTAGAAGCCTTTCTACCATTCCTAGT | GACTAGCATGATTGGAGTGGAAGGACAA | TGCCCGTTCATTGAGCACTATCCCATCA | AACCAACATAATAGGGGTACTTGGGCAG | 403 | P1 |
| *YKR056W 1* | GTCTCCTTTGACAGATTCTGGTAACCGC | ATTAACTACATCACCGGGCAGGCCAAAC | TAGCCCATTAACCGACAGCGGCAATAGA | GTTGACAACGTCGCCTGGTAAACCGAAA | 358 | P1 |
| *YKR056W 2* | TGGTGTAGAAATTTCCGCTGACAGTGTC | TATGCTTTCAATCTGGTGGGCGGAACCG | CGGCGTTGAAATCAGTGCCGATTCAGTT | GATTGATTCGATCTGATGAGCACTGCCA | 313 | P1 |
| *YKR058W 2* | CTCGGATCATAGCCCTGCTCCTAAC | ATTGTTATTGGAAGAAGCGGCCGTGCTT | TAGCGACCACTCACCAGCCCCAAAT | GTTATTGTTACTGCTGGCAGCGGTTGAG | 346 | P1 |
| *YKR058W 1* | CAGTGTCGAATCTAGCCCTTCAAATCCA | CGGTTGGTTGGTTTCCTCTGCTGTAAGT | TTCAGTTGAAAGCTCACCAAGCAACCCCT | TGGTTGATTAGTTTCCTCAGCGGTCAAC | 319 | P1 |
| *YKR060W 1* | CTTACTTCCGGAGGTTCTTGGTAGCAGA | TATTCCTCCTTTAATGACGCCGCCTTGC | TTTGTTGCCAGAGGTCTTGGGCTCACGT | GATACCACCCTTGATAACACCACCTTGA | 319 | P2 |
| *YKR061W 1* | ATATCAGAGACAGCTGAGCCTAGATTCG | GTCCTCCGCAGGGATTAAAGCGTATTGC | TTACCAGCGTCAGTTATCATTGGACAGC | ATCCTCAGCTGGAATCAAGGCATATTGG | 376 | P2 |
| *YKR062W 1* | CGACGAAGATTTTGGTAGCTCTCCTTCA | ATCGCATTGTGGCCAACCGTCTTTCAAG | TGATGAAGACTTCGGCTCAAGCCCAAGC | GTCACATTGAGGCCAGCCATCCTTTAAA | 385 | P2 |
| *YKR063C 2* | ATTGGTATCAGCCTTTAATGCCGGACTC | TGAGTGGGTAGCTAGGAATTACAGAACC | GTTAGTGTCGGCTTTCAAAGCTGGTGAA | CGAGTGGGTTGCCAGAAACTATCGTACT | 451 | P2 |
| *YKR063C 1* | GGACGGCAGTCCAATTTTAGCAGCTAAT | GGACTCTACAGCACAAATAACATGTGCG | ACTTGGTAAACCGATCTTGGCGGCCAAG | TGATAGCACCGCTCAAATCACCTGCGCT | 202 | P2 |
| *YKR064W 1* | CGATAGCTCTGGACAACCTGAAAGCAGT | GCCCTTACCAACAACATCATCGCTAATG | TGACTCAAGCGGTCAACCAGAATCATCA | ACCTTTGCCGACGACGTCGTCTGAGATA | 490 | P3 |

| Amplicon | WT_F | WT_R | syn_F | syn_R | bp | chunk |
|---|---|---|---|---|---|---|
| *YKR065C 1* | ACTTTGAGAAGAGG CATCGACCCTGTTA | CTTAAGATCATATTC TCAGCCCGCATCC | TGATTGGCTGCTAG CGTCAACTCTATTG | ATTGCGTAGCTACA GCCAGCCAGCTAGT | 382 | P3 |
| *YKR066C 1* | AGTGACACCCCCTA GACTGAACAGATCA | TGTCGCCTCTGTCG AAAAAGGGAGGTCA | GGTAACGCCACCCA ATGAAAATAAGTCG | CGTTGCTAGCGTTG AAAAGGGTAGAAGC | 325 | P3 |
| *YKR067W 2* * | GTCCCGATCCAGAA TGCCTTGTTTTGTT | AGTACCATTAGTCAA GAGCTCCACCACT | AAGTAGAAGTCGTAT GCCATGCTTCGTC | GGTGCCGTTGGTTA ACAACTCAACAACC | 376 | P3 |
| *YKR067W 1* | TTCTCATGACCGTC CTTCGTTACTACCC | TCTGGCAGCCTGAA TGACCATCAAAGTA | CAGCCACGATAGAC CAAGCTTGTTGCCA | ACGAGCGGCCTGGA TAACCATTAAGGTG | 319 | P4 |
| *YKR068C 1* | AGAGTCTCCTCTCA AGATATCCGACACA | TAGAACGGCATTGC CACGCTGTGAGAAT | GCTATCACCACGTA AAATGTCGCTAACG | CCGTACCGCTTTAC CTAGATGCGAGAAC | 298 | P4 |
| *YKR069W 2* | CATAGTAGTGAATC CCTCGTCACCAAGC | TTTTGGGTCGACCC ACGTAGGAATCATA | AATCGTTGTTAACCC AAGCAGCCCTTCA | CTTAGGATCAACCC AGGTTGGGATCATG | 295 | P4 |
| *YKR069W 1* | GGTTTTACCGGGCA TAAGCTCATCCCTA | ACCATCCCAGCCAT GCTTCAATAATCCC | TGTCTTGCCAGGTAT CTCAAGCAGTTTG | GCCGTCCCAACCGT GTTTTAACAAACCG | 226 | P4 |
| *YKR070W 1* | TAGGGACATTGCAC CATTTAGTGGGCTA | TGAAGGTTTCGTGC CTAAGAGTGGCAGT | CAGAGATATCGCTC CTTTCTCAGGTTTG | GCTTGGCTTGGTAC CCAACAAAGGTAAC | 481 | P4 |
| *YKR071C 1* | AGTTGGTAGCTTAC TACCACTCTGTAGC | TTTGACGCCAGAAG CCCAGACTGATATT | GGTAGGCAATTTTG AGCCTGACTGCAAT | CTTAACCCCTGAAG CTCAGACCGACATC | 268 | P4 |
| *YKR072C 2* | AGGTGCCAAAAGAA TGGGATAACTCGGA | TAAAATGTCGCAGTA CGAATCCACTCCG | TGGAGCTAACAAGA TTGGGTATGATGGG | CAAGATGAGCCAGT ATGAAAGTACCCCA | 307 | P4 |
| *YKR072C 1* | TCTACTTATTGGCGT TGAAGTGGCCATG | GTCTCCAAATAGTAA TCCTGCTCCTGTC | ACGTGAGATAGGGG TGCTGGTAGCCATA | AAGCCCTAACTCAAA CCCAGCCCCAGTT | 220 | P4/ P5 |
| *YKR075C 1* | AGAATGCTGGTCGA AAATGGAAGACGTG | AAGTTGCTCTTCAGA GGACGTGCTAGAT | GCTGTGCTGATCAA AGATACTGCTGGTA | TTCATGTAGCAGCG AGGATGTTTTGGAC | 412 | P5 |
| *YKR076W 1* | TACGAGGGCTTTGA AGGGATTAACCTCT | AGCAGGAACAAGGT CCGTTTTCTTGTGA | CACCAGAGCCTTAA AAGGTTTGACTAGC | GGCTGGGACCAAAT CGGTCTTTTTATGG | 445 | P5 |
| *YKR077W 1* | GTCATCGTCTTCAAT GAAGGTTCACGGC | CGGAGTTCTTGGGG AAGCTAACTTTTTG | TAGCAGCAGCAGCA TGAAAGTCCATGGT | TGGGGTACGAGGAC TGGCCAATTTCTTT | 361 | P5/ Q1 |
| *YKR078W 1* | AAGAAGACGTCGAA GACCAGAAAGCTGT | AATGTTGGACCTTGT AGGCGGAGGAATC | GCGTCGTAGAAGAC GTCCTGAATCATGC | GATATTACTTCTGGT TGGTGGTGGGATG | 280 | Q1 |
| *YKR078W 2* | GTCAACTGAGATAC AGGCATGCCATGAC | AATGCGGTATTCATC CTCTGCCGTGCGT | TAGCACCGAGATCC AGGCTTGTCACGAT | GATTCTATATTCGTC CTCAGCGGTTCTG | 469 | Q1 |
| *YKR079C 3* | CAGTGGAAAAATAC GCTGCTGTTCACCA | ACTAGAAAATCAGCT ACTGGAGGATGCC | TAAAGGGAAGATTCT CTGCTGTTCGCCG | CTTGGAAAACCAGTT GTTAGAGGACGCT | 244 | Q1 |
| *YKR079C 1* | ACCTCTGCTGAGTG GTGTATCGAATCTG | TGTTCCGAAGGGTC CCTTATTTGCAAAG | GCCACGTGACAAAG GGGTGTCAAAACGA | CGTCCCAAAAGGCC CATTGTTCGCTAAA | 466 | Q1 |

| Amplicon | WT_F | WT_R | syn_F | syn_R | bp | chunk |
|---|---|---|---|---|---|---|
| *YKR079C 2* | ATTCAAGTGCGGAT CACTAGAAGGATCG | TGGTGAAGGATCCC AAAGGAGTCTGACT | GTTTAAATGTGGGTC TGAGCTTGGGTCA | CGGCGAAGGTAGTC AAAGATCATTAACC | 439 | Q1 |
| *YKR080W 1* | ATCCATTCTACCTTG CACACCACTAGCT | TGCAAAATTGATGCA GACGGCACCTTCT | GAGTATCTTGCCAT GTACCCCTTTGGCC | AGCGAAGTTAATACA AACAGCGCCTTCC | 391 | Q1/ Q2 |
| *YKR081C 1* | AGAGGGCTCAATAT CAGTGGCACTGACG | CCAAGATGGTGAGC CATTGCCGAACGTT | GCTTGGCTCGATGT CGGTAGCTGAAACA | TCAAGACGGCGAGC CTTTACCAAATGTC | 445 | Q2 |
| *YKR082W 2* | CTCATTACGAAATG GACCTATCCTCGGT | CACCTGAGTAATGA CAACAGCATTGGGG | TAGCTTGAGAAACG GTCCAATTTTGGGC | AACCTGGGTGATAA CGACGGCGTTTGGA | 484 | Q2 |
| *YKR082W 3* | TACGTTGGGGCAAC ATCTCTCTGTCCGT | TGGTAGCTTGGATG TATCAACCTCCATC | CACCTTAGGTCAAC ACTTGAGCGTTAGA | AGGCAATTTACTGGT GTCGACCTCCATA | 469 | Q2 |
| *YKR082W 1* | ACCGCTAAGGCTTC TTTCTGTTGATGGA | GGCATTCTGATCAAT AGGAAACCTGCCA | CCCATTGAGATTGTT GAGCGTCGACGGT | AGCGTTCTGGTCGA TTGGGAATCTACCG | 274 | Q2 |
| *YKR084C 2* | CGCAATGGTTCTAC CGTCCTTTCTAAGA | AACGAATCACGAAG AGACAGATGTTGCC | AGCGATAGTACGGC CATCTTTACGCAAG | TACCAACCATGAAG AGACCGACGTCGCT | 430 | Q2 |
| *YKR084C 1* | TGCCCTATGGGTAG AGAAATGCGATGTA | GACTACGGTGCCAA CGAAACCAAAGAAA | AGCTCTGTGAGTGC TAAAGTGGCTGGTG | AACCACCGTTCCTA CCAAGCCTAAAAAG | 304 | Q2 |
| *YKR085C 1* | TTTTCTACGCTTACG GTCCTCTCTGGCT | TGAGACCATTCCAC GTAGCTTTATGGCA | CTTACGTCTTTTTCT ATCCTCACGAGCG | CGAGACTATCCCTA GATCATTCATGGCT | 364 | Q2 |
| *YKR086W 2* | TGAGACGTCCAGCC AAGTATCCGCTTTA | TGGCGTGAAAAGAG CTGTATCGTCGCAA | AGAGACCAGTTCAC AAGTTAGTGCCTTG | AGGGGTAAACAAGG CGGTGTCATCACAG | 259 | Q3 |
| *YKR086W 3* | GAGAAGAGTAGCAG CCATATCCGTTGCA | TACGGGGAAAGTTC TTCCAGGAATTGTG | TCGTCGTGTTGCTG CTATCAGTGTCGCT | AACTGGAAAGGTAC GACCTGGGATGGTA | 382 | Q3 |
| *YKR086W 1* | CCAAAGATCCGGAA GAGCGGGAAGA | CAATTGTCCACTAGT GTCTATAGCCCCT | TCAACGTAGTGGTC GTGCTGGTCGT | TAATTGACCTGAGGT ATCGATGGCACCG | 268 | Q3 |
| *YKR087C 1* | ACGAGTGCTTGCTG GATGTGTACTTAGA | CACAGCCGAAAATTT GTCGAAGGCTCCT | TCTGGTTGAAGCAG GGTGGGTTGACAAG | TACCGCTGAAAACTT AAGCAAAGCCCCA | 298 | Q3 |
| *YKR088C 1* | TCCTGTAGATTCAG ATTCTGCGAGACTC | CTATGGTGTCAGTTT TGAGGGATGGGTT | ACCGGTGCTTTCGC TTTCAGCCAATGAT | TTACGGCGTTTCATT CGAGGGTTGGGTC | 358 | Q3 |
| *YKR089C 1* | AGAAGAAGGCCTCG ATCGAAACGGTGAG | ACATTCTGCCAGCAT TGAAACACCTGCC | GCTGCTTGGTCTGC TTCTGAATGGGCTA | TCACAGCGCTTCAAT CGAAACCCCAGCT | 286 | Q3 |
| *YKR089C 2* | GAGAGAAGCGGAC GTAATGGAATACCG C | CTGTGGCCAGGAAT TTGCTCTGGATAAG | CAAGCTGGCACTGG TGATACTGTATCTA | TTGCGGTCAGGAAT TCGCCTTAGACAAA | 340 | Q3 |
| *YKR089C 3* | GCTCGAGGGGAAAA TTCCCGGTAGTGAA | CCACATCGGTGTCC TTGGTACTCTATTT | TGAGCTTGGAAAGA TACCTGGCAAGCTG | TCATATTGGCGTTTT GGGCACCTTGTTC | 457 | Q3 |
| *YKR090W 1* * | GAGTGCGTATACGG CGTCTTCAAAGAGT | TGTATGCCTAGAGG CAGCATATTGTTCC | ATCAGCTTACACCG CTAGCAGCAAATCA | GGTGTGTCTGCTAG CGGCGTATTGTTCA | 259 | Q3/ Q4 |

| Amplicon | WT_F | WT_R | syn_F | syn_R | bp | chunk |
|---|---|---|---|---|---|---|
| *YKR090W 2* | GTCAAGCGCATATT TATCTGGGTATCCC | AGGTCCCTCCCCAG GCGGATAT | AAGCTCAGCTTACTT GAGCGGTTACCCA | TGGACCCTCACCTG GTGGGTAC | 439 | Q4 |
| *YKR091W 1* | TTCCGTACGGGAGT TTTCACGCACGCTA | TGACCAACGTAACG GCGAAACGTTAACG | CAGTGTTAGAGAGT TCAGCAGAACCTTG | GCTCCATCTCAATG GGCTGACATTGACA | 304 | Q4 |
| *YKR092C 1* | GGCCTTGGATTCGT CTGCAGTTGCTTCT | GGCTAAAACTGAAC CAGAAAGCTCTTCC | AGCTTTACTTTCATC AGCGGTAGCTTCC | AGCCAAGACCGAAC CTGAATCAAGCAGT | 457 | Q4 |
| *YKR093W 1* | CACTTCGATTCCCT CCGTTGGTAACAGA | AACAGAAGGTTTAG CAGCGTTGAAGTCG | TACCAGCATCCCAA GTGTCGGCAATCGT | GACGCTTGGCTTGG CGGCATTAAAATCA | 466 | Q4 |
| *YKR093W 2* | TAAGAGAGCTTTGG CGGCTTGTAAAGTC | AGGTTCATTATACCA TGGACCAGCCTTG | CAAACGTGCCTTAG CTGCCTGCAAGGTT | TGGTTCGTTGTACCA AGGGCCGGCTTTA | 331 | Q4 |
| *YKR095W 1* | TTCTGTTTCTAATGA CTCGAAGGGACCC | GTATGACGCATCTTC GTTGGGAACAGAA | CAGCGTCAGCAACG ATAGCAAAGGTCCA | ATAGCTAGCGTCTTC ATTTGGGACGCTG | 391 | Q5 |
| *YKR095W 4* | GGATCTCTACGAGA CTACCTCTCAGTCT | AGCATTCTCCAGTTG ATCACGCGATAAG | AGACTTGTATGAGA CCACTAGCCAGAGC | GGCGTTCTCTAATTG GTCTCTGCTCAAA | 472 | Q5 |
| *YKR095W 6* | TGTTGCGCCTATCG AGTCCGAATTGACA | TCTCTCTTGCGCTTG TCTTCTCAGCCTG | CGTCGCTCCAATTG AGAGTGAATTAACC | ACGCTCTTGAGCTT GACGACGTAATCTA | 268 | Q5 |
| *YKR095W 5* | GGCTTCAAGAGAGC TTCAAGCCAAGTTA | TTTGAGGGCTTCCT CAGCTTCGCGAATC | AGCCAGCCGTGAGT TGCAAGCTAAATTG | CTTCAAAGCTTCCTC GGCTTCTCTGATT | 382 | Q5 |
| *YKR095W 3* | ACAAAGTGCCGAGA GTCCTCCGAAATCT | GGGTTCCAAATGAG TTGGTGGATCTGTC | GCAATCAGCTGAGT CACCACCAAAGAGC | TGGTTCTAAGTGGG TAGGAGGGTCGGTT | 262 | Q5 /R1 |
| *YKR095W 2* | CATTCCCGCCTCAA GGGGTCTAATATCT | CAGCTCTCCTACCT CATCAATAGGTCGC | TATCCCAGCTAGCA GAGGCTTGATCAGC | TAACTCACCAACCTC GTCGATTGGTCTT | 229 | Q5/ R1 |
| *YKR096W 3* | GCCAGCAAGTAATA AGAGATCGGGCATT | CGGTTGGGATGGTA GTAAAGCTGTAGTG | ACCTGCTTCAAACAA ACGTAGCGGTATC | TGGTTGACTAGGCA ACAAGGCGGTGGTA | 223 | R1 |
| *YKR096W 4* | CGAAAGAAGGCTAT GGGTTTACGGCACG | CGGAGTGAATGTTT CCTGACAAAAGACG | TGAACGTAGATTGT GGGTCTATGGTACC | TGGGGTAAAGGTTT CCTGGCAGAAAACA | 382 | R1 |
| *YKR096W 1* | TGACTCCTCCGTGA CGGAAAGTTCAACG | TGCGATACCTAAAG GCCCATGTAGAAAC | CGATAGTAGTGTTAC CGAATCAAGCACC | AGCAATGCCCAATG GACCGTGCAAGAAT | 268 | R1 |
| *YKR096W 2* | AACTTCTTGGCTAA GACATTCCGCACGC | TGTACACAGATGGC GCTGTTCGCCAAGT | TACCAGCTGGTTGC GTCACAGTGCTAGA | GGTGCATAAGTGTC TCTGTTCACCCAAC | 490 | R1 |
| *YKR097W 1* | CATTTCATCAAGCG GTGCATTGATCGCT | GTGTAAATTGGCTG GGAACTGACCAGCG | TATCAGCAGCTCAG GCGCTTTAATTGCC | ATGCAAGTTAGCAG GAAACTGGCCGGCA | 385 | R1 |
| *YKR097W 2* | CGGCTGGACTGGTT CTTCCTACGTATCT | GGCCAAGTTGGTAA CTGCACCTCTGTAT | TGGTTGGACCGGCA GCAGTTATGTTAGC | AGCTAAATTAGTGAC AGCGCCACGATAC | 238 | R1 |
| *YKR098C 2* * | AAGGTTACCGGAAT GATTCACCGTTCCA | TCACCATCATCACCA TAGCAGCGATGAT | CAAATTGCCACTGT GGTTAACGGTACCG | CCATCACCACCATC ACTCATCAGACGAC | 259 | R1 |

| Amplicon | WT_F | WT_R | syn_F | syn_R | bp | chunk |
|---|---|---|---|---|---|---|
| *YKR098C 1* | AGCAGTTGCTGGAC TAGACAGCGTTTTG | CTTACCGCTACGGC CTTTACCTGTTCAT | GGCGGTAGCAGGTG AGCTTAAGGTCTTA | ATTGCCATTGAGAC CATTGCCAGTCCAC | 286 | R2 |
| *YKR099W 2* | AATCTCAGATAGCC CACAGACAAGCCTT | TCCAGTATTGCTGCT GACAGGATGGTCT | TATTAGCGACTCACC TCAGACCTCATTG | ACCGGTGTTTGATG AAACTGGGTGATCG | 334 | R2 |
| *YKR099W 1* | AAGTTCGACGGCAT CATATACAACGAGC | CCCTTGATGTACCA CTTGAGGCTTGGCA | TTCAAGCACCGCTA GCTACACCACCTCA | ACCTTGGTGAACAA CTTGTGGTTTAGCG | 448 | R2 |
| *YKR100C 1* | TAAAAACGAGGATG AAGCTCTCCTTGGG | TAATGCCAGGCAGA AGTCCGTTGTGGAT | CAAGAAGCTACTGC TGGCACGTCTAGGA | CAACGCTAGACAGA AAAGTGTCGTTGAC | 319 | R2 |
| *YKR101W 1* | GAAGGATTCTAATT CGGTGCTCGTTGGA | ACATACCCCCGCTA AGGGCTTTTCTTTA | AAAAGACAGCAACA GCGTTTTGGTCGGT | GCAAACACCAGCCA ATGGTTTTTCCTTG | 250 | R2 |
| *YKR101W 2* | TTCGAAATTTCCTGT GTCCAGAGTCACC | TCCATCTGCTACTAA AAACCTCGGCGAA | CAGCAAGTTCCCAG TTAGTCGTGTTACT | ACCGTCAGCAACCA AGAATCTTGGGCTG | 352 | R2 |
| *YKR102W 1* | CAGCCTATCTTCAA CTGTAAGCGAGCAT | TGAGGGTAGAATGG TAACTGAGGCCGAG | TTCATTGAGCAGCA CCGTTTCAGAGCAC | GCTTGGCAAGATAG TGACGCTAGCGCTT | 238 | R3 |
| *YKR102W 2* | CGTAACATCTAGCT CTGTTGTCTCCACA | AACGGAGGTGGAGG CTGTAGAAGTTTCG | TGTTACCAGCTCAA GCGTCGTTAGTACC | GACACTAGTACTAG CGGTGCTGGTTTCA | 481 | R3 |
| *YKR102W 4* | ATCTTCTCAGTTTGT GACTCCATCCTCC | CCTATACACAGTGG TGCGATCACGAGAA | CAGCAGCCAGTTCG TTACCCCTAGTAGT | TCTGTAAACGGTAGT TCTGTCTCTGCTG | 226 | R3 |
| *YKR102W 3* | TAGCTGCTCAAACA CAGTCTCAAGTGCT | ACTTTTGCTACTGAC AGCCTCAGAAGCA | CTCATGTAGCAATAC CGTTAGCTCAGCC | TGACTTTGATGAAAC GGCCTCGCTGGCT | 487 | R3/ R4 |
| *YKR103W 4* | ATCTGCATGGGAAA GCCGTAGCTTTTGT | ACATGCTAAAACCCA CCGAAATGACTGG | GAGCGCTTGGGAAT CAAGATCATTCTGC | GCAAGCCAAGACCC ATCTGAAGCTCTGA | 259 | R4 |
| *YKR103W 3* | CTCTGGGATTAGTG AGGCTCTGAACTCA | AGCTTCCCATTCCAT GCTTGCATCTTCA | TAGCGGTATCTCAG AGGCCTTAAATAGC | GGCTTCCCATTCCAT TGAAGCGTCTTCG | 439 | R4 |
| *YKR103W 2* | GACTGGGTCAGGAA AGTCTTCGCTATTA | TGCAGTTTCAGGAT CTACTGCACTCAAG | TACCGGTAGCGGTA AAAGCAGCTTGTTG | AGCGGTTTCTGGGT CAACAGCTGATAAA | 421 | R4 |
| *YKR103W 1* | CACTGGATCTTCAG GGAGTAAGTCTACC | GTTCATTATTCTTCC AGTCGGCGTGACA | TACCGGTAGCAGCG GTTCAAAAAGCACT | ATTCATGATACGACC GGTTGGGGTAACG | 247 | R4 |
| *YKR104W 1* | TATAGCGGCTATTTA CCGGCTTTCAGAC | TGAGGCGGTTGCTT CGTCAATTAGCATT | CATCGCTGCCATCT ATAGATTGAGCGAT | GCTAGCAGTAGCTT CATCGATCAACATG | 427 | R4 |

***Table S3* - synXI *PCRTag primers. "Amplicon" given the chromosomal target locus. "WT_F" is the forward primer targeting wild type sequence. "WT_R" is the reverse primer targeting wild type sequence. "syn_F" is the forward primer targeting* synXI *sequence. "syn_R" is the reverse timer targeting synXI sequence. "bp" is the expected size, in bp, of the PCR amplicon. "chunk" is the* synXI *chunk or chunks that contain the corresponding target sequence. Asterisks denote that primer sequences deviate from target sequences.***

| Purpose | Target | Target sequence | Retargeting primer | Retargeting primer sequence | Repair template |
|---|---|---|---|---|---|
| M Debugging | M1 - *YKL005C*_2_WT_R | GGAAACGTCAAAGGAGGCAG[AGG] | BB392 | CTGCCTCCTTTGACGTTTCCaaagtcccattcgccacccg | M1 chunk |
| M Debugging | M2 - *YKR001C*_2_WT_F | ACTGGCGCCTTACTAGACGA[TGG] | BB394 | TCGTCTAGTAAGGCGCCAGTaaagtcccattcgccacccg | M2 chunk |
| M Debugging | M3 - *YKR006C*_1_WT_R | TATTGAAGAAGGCACTAACG[AGG] | BB396 | CGTTAGTGCCTTCTTCAATAaaagtcccattcgccacccg | M3 chunk |
| M Debugging | M4 - *YKR010C*_2_WT_F | CTAGTGGTGCTCATAGTCCC[AGG] | BB398 | GGGACTATGAGCACCACTAGaaagtcccattcgccacccg | M4 chunk |
| M Debugging | M5 - *YKR013W*_1_WT_F | TGAGGATTGAGTGCTTGAGG[CGG] | BB400 | CCTCAAGCACTCAATCCTCAaaagtcccattcgccacccg | M5 chunk |
| *CEN11** replacement | *KlURA3* 3' region | CTTCGGGAATCGCCTATCAG[CGG] | BB875 | CTGATAGGCGATTCCCGAAGaaagtcccattcgccacccg | *CEN11* variant BB877/BB878 PCR product |
| *CEN11** replacement | *YKR001C*_2_WT_F | ACTGGCGCCTTACTAGACGA[TGG] | BB394 | TCGTCTAGTAAGGCGCCAGTaaagtcccattcgccacccg | M2 chunk |
| *HIS3-TR2* removal | HIS3 | TTTTTACTCCACGCGCCAGT[AGG] | BB417 | ACTGGCGCGTGGAGTAAAAAaaagtcccattcgccacccg | *his3*Δ1 as BB418/BB419 PCR product |
| O integration patching | *YKR051W*_1_WT_R | ATTCTTGTACACATATCCAG[AGG] | BB390 | CTGGATATGTGTACAAGAATaaagtcccattcgccacccg | O3-O4 chunk ligation product |
| R5 *URA3* removal | *URA3* | GCAGACATTACGAATGCGCA[CGG] | BB486 | TGCGCATTCGTAATGTCTGCaaagtcccattcgccacccg | Neighbouring sequence |
| Mitotic crossover | O5 *LEU2* | GAAAAAAATAAAAATGGAGT[AGG] | BB457 | ACTCCATTTTTATTTTTTTCaaagtcccattcgccacccg | Sister chromatid |
| J repeat condensation | *LEU2* | CCAGCGCCTCATCTGGAAGT[GGG] | BB573 | ACTTCCAGATGAGGCGCTGGaaagtcccattcgccacccg | Megachunk J |
| J repeat condensation | *kanMX* | ATGAAGGAGAAAACTCACCG[AGG] | BB574 | CGGTGAGTTTTCTCCTTCATaaagtcccattcgccacccg | Megachunk J |
| J repeat condensation | J1 vector backbone sequence (5') | ACTGGAAAGCGGGCAGTGAG[CGG] | BB604 | CTCACTGCCCGCTTTCCAGTaaagtcccattcgccacccg | Neighbouring sequence |
| J repeat condensation | J1 vector backbone sequence (3') | CTATAGGGCGAATTGGCGGA[AGG] | BB603 | TCCGCCAATTCGCCCTATAGaaagtcccattcgccacccg | Neighbouring sequence |
| Q repeat condensation | *URA3* | GCAGACATTACGAATGCGCA[CGG] | BB486 | TGCGCATTCGTAATGTCTGCaaagtcccattcgccacccg | Megachunk Q |
| *TRK2* insertion removal | *TRK2* insertion - transposon sequence | GCTCAAAATTTATTCACACA[TGG] | BB736 | TGTGTGAATAAATTTTGAGCaaagtcccattcgccacccg | Neighbouring sequence |
| Q debugging | P5 *URA3* | GAAAAGCTTGCGGTAGTGAA[GGG] | BB783 | TTCACTACCGCAAGCTTTTCaaagtcccattcgccacccg | Megachunk Q |
| Q debugging | *YKR077W*_1_WT_R | GTTTGCATTCGGAGTTCTTG[GGG] | BB704 | CAAGAACTCCGAATGCAAACaaagtcccattcgccacccg | Megachunk Q |

| Purpose | Target | Target sequence | Retargeting primer | Retargeting primer sequence | Repair template |
|---|---|---|---|---|---|
| Q debugging | *YKR084C*_2_WT_F | AGAGTTGTTCTTAGAAAGGA[CGG] | BB705 | TCCTTTCTAAGAACAACTCTaaagtcccattcgccacccg | Megachunk Q |
| Glycerol locus replacement | *YKR084C*_1_syn_F | GCTAACGCCGATCAACGTAG[TGG] | BB757 | CTACGTTGATCGGCGTTAGCaaagtcccattcgccacccg | Debug colony 2 locus BB768/BB775 PCR product |
| Glycerol locus replacement | *YKR086W*_1_syn_R | CAATGGAGTTAATTGACCTG[AGG] | BB778 | CAGGTCAATTAACTCCATTGaaagtcccattcgccacccg | Debug colony 2 locus BB768/BB775 PCR product |
| *PRP16* stop swap | *PRP16* TAG | TCTTCCCATAAACTAAAAAA[AGG] | BB785 | TTTTTTTAGTTTATGGGAAGaaagtcccattcgccacccg | Q3 chunk BB738/BB788 PCR product |
| *GAP1*$_{syn}$ generation | *YKRCδ11* | TCACCCATTTGTCAAGATAA[TGG] | XL498 | TTTGGTCTCGCGCATCACCCATTTGTCAAGATAAGTTTTAGAGCTAGAAATAGCAAGTTA | synthetic *GAP1* locus amplicon |
| | | | XL499 | TTTGGTCTCGTTGATATAAGCCCTGCGCAAGCCCGGAATCGAAC | |
| *GAP1*$_{syn}$ generation | *YKRCδ12* | GGGCTTATATCAAGATCTGT[TGG] | XL500 | TTTGGTCTCCTCAAGATCTGTGTTTTAGAGCTAGAAATAGCAAGTTA | synthetic *GAP1* locus amplicon |
| | | | XL501 | TTTGGTCTCCGGATTGCGCAAGCCCGGAATCGAAC | |
| *yEGFP* insertion | sequence upstream of *GAP1* | TTTGTCGAAGATATTCGACG[AGG] | XL502 | AGATTTTGTCGAAGATATTCGACG | *PPFY1-yEGFP-TCYC1* |
| | | | XL503 | AAACCGTCGAATATCTTCGACAAA | |

***Table S4 - CRISPR/Cas9 target sequences and gRNA retargeting primers. Target sequence Protospacer Adjacent Motifs are shown in square brackets. Retargeting primer sequences are shown with the pWS082 binding sequence in lowercase and the retargeting sequence in uppercase.***

| Version | Strain(s) | Comment | Details |
|---|---|---|---|
| *synXI*_3.34 | N/A | | Final design by BioStudio |
| *synXI*_3.36 | N/A | | TAG stop codons in *YKL006C-A*, *YKR004C* and *YKR005C* recoded to TAA |
| *synXI*_3.37 | N/A | | *CEN11* right arm loxPsym site moved to 3 bp 3' of *YKR001C* |
| *synXI*_9.01 | ysXIb01 | Initial synXI assembly | Missing loxPsym sites: 526668-526701 (*YKR052C*). WT PCRTags: 466241-466268 (*YKR016W*_1_F), 493836-493863 (*YKR029C*_2_R),493911-493938 (*YKR029C*_1_F), 536959-536986 (*YKR054C*_6_R). Point mutations causing amino acid changes: 37065 T>C (*YKL210W*), 448094 G>C (*YKR008W*). Structural variations: 303238-328404 (repeated sequence), 523727-525128 (insertions/duplication at *YKR050W*), 578748-591038 (repeated sequence). |
| *synX*I_9.02 | ysXIb02 | Markers inserted flanking megachunk J repeated sequence | Missing loxPsym sites: 526668-526701 (*YKR052C*). WT PCRTags: 466241-466268 (*YKR016W*_1_F), 493836-493863 (*YKR029C*_2_R),493911-493938 (*YKR029C*_1_F), 536959-536986 (*YKR054C*_6_R). Point mutations causing amino acid changes: 37065 T>C (*YKL210W*), 448094 G>C (*YKR008W*). Structural variations: 303238-328404 (repeated sequence), 523727-525128 (insertions/duplication at *YKR050W*), 578748-591038 (repeated sequence). Marker gene insertions: 303694 (*YKL0069W*::*LEU2*), 334347 (*YKL053C*::*kanMX4*). |

| Version | Strain(s) | Comment | Details |
|---|---|---|---|
| *synXI*_9.03 | ysXIb03 | Megachunk J repeated sequence copy number reduced - respiratory growth defect present | Missing loxPsym sites: 526668-526701 (*YKR052C*). WT PCRTags: 466241-466268 (*YKR016W*_1_F), 493836-493863 (*YKR029C*_2_R),493911-493938 (*YKR029C*_1_F), 536959-536986 (*YKR054C*_6_R). Point mutations causing amino acid changes: 37065 T>C (*YKL210W*), 448094 G>C (*YKR008W*). Structural variations: 303238-311650 (repeated sequence with pMA-RQ insertion at 311649), 523727-525128 (insertions/duplication at *YKR050W*), 578748-591038 (repeated sequence). Marker gene insertions: 343787 (*YKL048C*::*URA3*). |
| *synXI*_9.04 | ysXIb04 | *URA3* marker removed - respiratory growth defect present | Missing loxPsym sites: 526668-526701 (*YKR052C*). WT PCRTags: 466241-466268 (*YKR016W*_1_F), 493836-493863 (*YKR029C*_2_R),493911-493938 (*YKR029C*_1_F), 536959-536986 (*YKR054C*_6_R). Point mutations causing amino acid changes: 37065 T>C (*YKL210W*), 448094 G>C (*YKR008W*). Structural variations: 303238-311650 (repeated sequence with pMA-RQ insertion at 311649), 523727-525128 (insertions/duplication at *YKR050W*), 578748-591038 (repeated sequence). |
| *synXI*_9.05 | ysXIb05 | Megachunk J repeated sequence removed - respiratory growth defect present | Missing loxPsym sites: 526668-526701 (*YKR052C*). WT PCRTags: 466241-466268 (*YKR016W*_1_F), 493836-493863 (*YKR029C*_2_R),493911-493938 (*YKR029C*_1_F), 536959-536986 (*YKR054C*_6_R). Point mutations causing amino acid changes: 37065 T>C (*YKL210W*), 448094 G>C (*YKR008W*). Structural variations: 523727-525128 (insertions/duplication at *YKR050W*), 578748-591038 (repeated sequence). |
| *synXI*_9.06 | ysXIb06 | *URA3* inserted upstream of megachunk Q repeated sequence - respiratory growth defect present | Missing loxPsym sites: 526668-526701 (*YKR052C*). WT PCRTags: 466241-466268 (*YKR016W*_1_F), 493836-493863 (*YKR029C*_2_R),493911-493938 (*YKR029C*_1_F), 536959-536986 (*YKR054C*_6_R). Point mutations causing amino acid changes: 37065 T>C (*YKL210W*), 448094 G>C (*YKR008W*). Structural variations: 523727-525128 (insertions/duplication at *YKR050W*), 578748-591038 (repeated sequence). Marker gene insertions: 579173 (*YKR077W*::*URA3*). |
| *synXI*_9.08 | ysXIb08 | Megachunk Q repeated sequence removed - respiratory growth defect present | Missing loxPsym sites: 526668-526701 (*YKR052C*). WT PCRTags: 466241-466268 (*YKR016W*_1_F), 493836-493863 (*YKR029C*_2_R),493911-493938 (*YKR029C*_1_F), 536959-536986 (*YKR054C*_6_R). Point mutations causing amino acid changes: 37065 T>C (*YKL210W*), 448094 G>C (*YKR008W*). Structural variations: 523727-525128 (insertions/duplication at *YKR050W*). |
| *synXI*_9.09 | ysXIb09 | *TRK2*/*YKR050W* insertions/duplication removed - respiratory growth defect present | Missing loxPsym sites: 526668-526701 (*YKR052C*). WT PCRTags: 466241-466268 (*YKR016W*_1_F), 493836-493863 (*YKR029C*_2_R),493911-493938 (*YKR029C*_1_F), 536959-536986 (*YKR054C*_6_R). Point mutations causing amino acid changes: 37065 T>C (*YKL210W*), 448094 G>C (*YKR008W*). |
| *synXI*_9.10 | ysXIb10, ysXIb11, ysXIb12 | Respiratory growth defect fixed | Remaining TAG stop codons: 597913 A>G (*YKR086W*). Missing loxPsym sites: 526668-526701 (*YKR052C*), 593898-593931 (*YKR085C*),598033-598066 (*YKR087C*). WT PCRTags: 466241-466268 (*YKR016W*_1_F), 493836-493863 (*YKR029C*_2_R),493911-493938 (*YKR029C*_1_F), 536959-536986 (*YKR054C*_6_R), 592963-592990 (*YKR084C*_1_F), 593239-593266 (*YKR084C*_1_R), 593956-593983 (*YKR085C*_1_F), 594292-594319 (*YKR085C*_1_R), 595453-595480 (*YKR086W*_2_F), 595684-595711 (*YKR086W*_2_R), 595912-595939 (*YKR086W*_3_F), 596266-596293 (*YKR086W*_3_R), 596749-596773 (*YKR086W*_1_F), 596989-597016 (*YKR086W*_1_R). Point mutations causing amino acid changes: 37065 T>C (*YKL210W*), 448094 G>C (*YKR008W*). |
| *synXI*_9.11 | ysXIb13, ysXIb14, ysXIb16, ysXIb17 | Final version | Missing loxPsym sites: 526668-526701 (*YKR052C*), 593898-593931 (*YKR085C*),598033-598066 (*YKR087C*). WT PCRTags: 466241-466268 (*YKR016W*_1_F), 493836-493863 (*YKR029C*_2_R),493911-493938 (*YKR029C*_1_F), 536959-536986 (*YKR054C*_6_R), 592963-592990 (*YKR084C*_1_F), 593239-593266 (*YKR084C*_1_R), 593956-593983 (*YKR085C*_1_F), 594292-594319 (*YKR085C*_1_R), 595453-595480 (*YKR086W*_2_F), 595684-595711 (*YKR086W*_2_R), 595912-595939 (*YKR086W*_3_F), 596266-596293 (*YKR086W*_3_R), 596749-596773 |

| Version | Strain(s) | Comment | Details |
|---|---|---|---|
| | | | (*YKR086W*_1_F), 596989-597016 (*YKR086W*_1_R). Point mutations causing amino acid changes: 37065 T>C (*YKL210W*), 448094 G>C (*YKR008W*). |

***Table S5 - synXI chromosome versions. For* in vivo *chromosome versions, variations from synXI_3.37 are listed.***

| Plasmid | Description | Use | Source |
|---|---|---|---|
| p*CEN11** | Contains *CEN11* region with upstream *KlURA3* and *GAL1* promoter | Inducible disruption of centromere | This study |
| p*CEN11*_3_37b | Modified M2 vector containing *CEN11*_3_37b | Centromere variant testing | This study |
| p*CEN11*_3_37c | Modified M2 vector containing *CEN11*_3_37c | Centromere variant testing | This study |
| p*CEN11*_3_37f | Modified M2 vector containing *CEN11*_3_37f | Centromere variant testing | This study |
| pRS413-*chrXI*_tRNA | Vector expressing tRNAs complementing those removed in *synXI* | tRNA complementation | Schindler, D. *et al*, unpublished |
| pHLUM | Vector expressing *HIS3, LEU2, URA3* and *MET15* | Complementation of auxotrophies | Mülleder *et al*., 2012 |
| pJCH021 | MoClo-YTK part plasmid: *PSCW11* | Generation of p*SCW11*-*cre*-EBD-*kanMX4* | Ellis lab |
| pJCH022 | MoClo-YTK part plasmid: *cre*-EBD | Generation of p*SCW11*-*cre*-EBD-*kanMX4* | Ellis lab |
| pJJH1304 | Source of *Kluyveromyces lactis URA3* expression cassette | Generation of p*CEN11** | Addgene, Bakota *et al*., 2012 |
| pRS403 | Yeast *HIS3* integration vector | Generation of pRS403::*TRT2* | Sikorski *et al*., 1989 |
| pRS403::*TRT2* | *HIS3* integrative vector expressing *TRT2* | *TRT2* complementation | This study |
| pRS405 | Yeast *LEU2* integration vector | Generation of pRS405-*LEU2*::*URA3* | Sikorski *et al*., 1989 |
| pRS405-*LEU2*::*URA3* | Integrative vector for insertion of *URA3* into *LEU2* | Auxotrophy switching | This study |
| pRS406 | Yeast *URA3* integration vector | Generation of pRS405-*LEU2*::*URA3* | Sikorski *et al*., 1989 |
| pRS415 | Yeast centromeric plasmid with *LEU2* | Generation of pZY412 | Sikorski *et al*., 1989 |
| p*SCW11*-*cre*-EBD | Cre expression vector with daughter cell-specific and β-estradiol induced expression | Generation of p*SCW11*-*cre*-EBD-*kanMX4* | Cai *et al*., 2015 |
| p*SCW11*-*cre*-EBD-*kanMX4* | Cre expression vector with daughter cell-specific and β-estradiol induced expression and a *kanMX4* marker gene | SPecc formation | This study |
| pSV-PFY1p | Vector expressing *yEGFP* under constitutive *PPFY1* promoter | chromosomal *yEGFP* expression cassette | Blount *et al*., 2021 |
| pSXI_3_34_A1 | Vector containing synthetic chunk A1 | Chunk propagation for *synXI* assembly | This study - GenScript |
| pSXI_3_34_A2 | Vector containing synthetic chunk A2 | Chunk propagation for *synXI* assembly | This study - GenScript |
| pSXI_3_34_B1 | Vector containing synthetic chunk B1 | Chunk propagation for *synXI* assembly | This study - GenScript |
| pSXI_3_34_B2 | Vector containing synthetic chunk B2 | Chunk propagation for *synXI* assembly | This study - GenScript |
| pSXI_3_34_B3 | Vector containing synthetic chunk B3 | Chunk propagation for *synXI* assembly | This study - GenScript |
| pSXI_3_34_B4 | Vector containing synthetic chunk B4 | Chunk propagation for *synXI* assembly | This study - GenScript |
| pSXI_3_34_B5 | Vector containing synthetic chunk B5 | Chunk propagation for *synXI* assembly | This study - GenScript |
| pSXI_3_34_C1 | Vector containing synthetic chunk C1 | Chunk propagation for *synXI* assembly | This study - GenScript |
| pSXI_3_34_C2 | Vector containing synthetic chunk C2 | Chunk propagation for *synXI* assembly | This study - GenScript |
| pSXI_3_34_C3 | Vector containing synthetic chunk C3 | Chunk propagation for *synXI* assembly | This study - GenScript |
| pSXI_3_34_C4 | Vector containing synthetic chunk C4 | Chunk propagation for *synXI* assembly | This study - GenScript |
| pSXI_3_34_C5 | Vector containing synthetic chunk C5 | Chunk propagation for *synXI* assembly | This study - GenScript |
| pSXI_3_34_D1 | Vector containing synthetic chunk D1 | Chunk propagation for *synXI* assembly | This study - GeneArt |

| Plasmid | Description | Use | Source |
|---|---|---|---|
| pSXI_3_34_D2 | Vector containing synthetic chunk D2 | Chunk propagation for *synXI* assembly | This study - GeneArt |
| pSXI_3_34_D3 | Vector containing synthetic chunk D3 | Chunk propagation for *synXI* assembly | This study - GeneArt |
| pSXI_3_34_D4 | Vector containing synthetic chunk D4 | Chunk propagation for *synXI* assembly | This study - GeneArt |
| pSXI_3_34_D5 | Vector containing synthetic chunk D5 | Chunk propagation for *synXI* assembly | This study - GeneArt |
| pSXI_3_34_E1 | Vector containing synthetic chunk E1 | Chunk propagation for *synXI* assembly | This study - GeneArt |
| pSXI_3_34_E2 | Vector containing synthetic chunk E2 | Chunk propagation for *synXI* assembly | This study - GeneArt |
| pSXI_3_34_E3 | Vector containing synthetic chunk E3 | Chunk propagation for *synXI* assembly | This study - GeneArt |
| pSXI_3_34_E4 | Vector containing synthetic chunk E4 | Chunk propagation for *synXI* assembly | This study - GeneArt |
| pSXI_3_34_E5 | Vector containing synthetic chunk E5 | Chunk propagation for *synXI* assembly | This study - GeneArt |
| pSXI_3_34_F1 | Vector containing synthetic chunk F1 | Chunk propagation for *synXI* assembly | This study - GeneArt |
| pSXI_3_34_F2 | Vector containing synthetic chunk F2 | Chunk propagation for *synXI* assembly | This study - GeneArt |
| pSXI_3_34_F3 | Vector containing synthetic chunk F3 | Chunk propagation for *synXI* assembly | This study - GeneArt |
| pSXI_3_34_F4 | Vector containing synthetic chunk F4 | Chunk propagation for *synXI* assembly | This study - GeneArt |
| pSXI_3_34_F5 | Vector containing synthetic chunk F5 | Chunk propagation for *synXI* assembly | This study - GeneArt |
| pSXI_3_34_G1 | Vector containing synthetic chunk G1 | Chunk propagation for *synXI* assembly | This study - GeneArt |
| pSXI_3_34_G2 | Vector containing synthetic chunk G2 | Chunk propagation for *synXI* assembly | This study - GeneArt |
| pSXI_3_34_G3 | Vector containing synthetic chunk G3 | Chunk propagation for *synXI* assembly | This study - GeneArt |
| pSXI_3_34_G4 | Vector containing synthetic chunk G4 | Chunk propagation for *synXI* assembly | This study - GeneArt |
| pSXI_3_34_G5 | Vector containing synthetic chunk G5 | Chunk propagation for *synXI* assembly | This study - GeneArt |
| pSXI_3_34_H1 | Vector containing synthetic chunk H1 | Chunk propagation for *synXI* assembly | This study - GeneArt |
| pSXI_3_34_H2 | Vector containing synthetic chunk H2 | Chunk propagation for *synXI* assembly | This study - GeneArt |
| pSXI_3_34_H3 | Vector containing synthetic chunk H3 | Chunk propagation for *synXI* assembly | This study - GeneArt |
| pSXI_3_34_H4 | Vector containing synthetic chunk H4 | Chunk propagation for *synXI* assembly | This study - GeneArt |
| pSXI_3_34_H5 | Vector containing synthetic chunk H5 | Chunk propagation for *synXI* assembly | This study - GeneArt |
| pSXI_3_34_I1 | Vector containing synthetic chunk I1 | Chunk propagation for *synXI* assembly | This study - GeneArt |
| pSXI_3_34_I2 | Vector containing synthetic chunk I2 | Chunk propagation for *synXI* assembly | This study - GeneArt |
| pSXI_3_34_I3 | Vector containing synthetic chunk I3 | Chunk propagation for *synXI* assembly | This study - GeneArt |
| pSXI_3_34_I4 | Vector containing synthetic chunk I4 | Chunk propagation for *synXI* assembly | This study - GeneArt |
| pSXI_3_34_I5 | Vector containing synthetic chunk I5 | Chunk propagation for *synXI* assembly | This study - GeneArt |
| pSXI_3_34_J1 | Vector containing synthetic chunk J1 | Chunk propagation for *synXI* assembly | This study - GeneArt |
| pSXI_3_34_J2 | Vector containing synthetic chunk J2 | Chunk propagation for *synXI* assembly | This study - GeneArt |
| pSXI_3_34_J3 | Vector containing synthetic chunk J3 | Chunk propagation for *synXI* assembly | This study - GeneArt |
| pSXI_3_34_J4 | Vector containing synthetic chunk J4 | Chunk propagation for *synXI* assembly | This study - GeneArt |
| pSXI_3_34_J4::*kanMX* | J4 chunk vector with *kanMX4* insertion in *YKL053C-A* | Marker insertion | This study |
| pSXI_3_34_J5 | Vector containing synthetic chunk J5 | Chunk propagation for *synXI* assembly | This study - GeneArt |
| pSXI_3_34_K1 | Vector containing synthetic chunk K1 | Chunk propagation for *synXI* assembly | This study - GeneArt |
| pSXI_3_34_K2 | Vector containing synthetic chunk K2 | Chunk propagation for *synXI* assembly | This study - GeneArt |

| Plasmid | Description | Use | Source |
|---|---|---|---|
| pSXI_3_34_K3 | Vector containing synthetic chunk K3 | Chunk propagation for *synXI* assembly | This study - GeneArt |
| pSXI_3_34_K4 | Vector containing synthetic chunk K4 | Chunk propagation for *synXI* assembly | This study - GeneArt |
| pSXI_3_34_K5 | Vector containing synthetic chunk K5 | Chunk propagation for *synXI* assembly | This study - GeneArt |
| pSXI_3_34_L1 | Vector containing synthetic chunk L1 | Chunk propagation for *synXI* assembly | This study - GeneArt |
| pSXI_3_34_L2 | Vector containing synthetic chunk L2 | Chunk propagation for *synXI* assembly | This study - GeneArt |
| pSXI_3_34_L3 | Vector containing synthetic chunk L3 | Chunk propagation for *synXI* assembly | This study - GeneArt |
| pSXI_3_34_L4 | Vector containing synthetic chunk L4 | Chunk propagation for *synXI* assembly | This study - GeneArt |
| pSXI_3_34_L5 | Vector containing synthetic chunk L5 | Chunk propagation for *synXI* assembly | This study - GeneArt |
| pSXI_3_34_M1 | Vector containing synthetic chunk M1 | Chunk propagation for *synXI* assembly | This study - GeneArt |
| pSXI_3_34_M2 | Vector containing synthetic chunk M2 | Chunk propagation for *synXI* assembly | This study - GeneArt |
| pSXI_3_34_M3 | Vector containing synthetic chunk M3 | Chunk propagation for *synXI* assembly | This study - GeneArt |
| pSXI_3_34_M4 | Vector containing synthetic chunk M4 | Chunk propagation for *synXI* assembly | This study - GeneArt |
| pSXI_3_34_M5 | Vector containing synthetic chunk M5 | Chunk propagation for *synXI* assembly | This study - GeneArt |
| pSXI_3_34_N1 | Vector containing synthetic chunk N1 | Chunk propagation for *synXI* assembly | This study - GeneArt |
| pSXI_3_34_N2 | Vector containing synthetic chunk N2 | Chunk propagation for *synXI* assembly | This study - GeneArt |
| pSXI_3_34_N3 | Vector containing synthetic chunk N3 | Chunk propagation for *synXI* assembly | This study - GeneArt |
| pSXI_3_34_N4 | Vector containing synthetic chunk N4 | Chunk propagation for *synXI* assembly | This study - GeneArt |
| pSXI_3_34_N5 | Vector containing synthetic chunk N5 | Chunk propagation for *synXI* assembly | This study - GeneArt |
| pSXI_3_34_O1 | Vector containing synthetic chunk O1 | Chunk propagation for *synXI* assembly | This study - GeneArt |
| pSXI_3_34_O2 | Vector containing synthetic chunk O2 | Chunk propagation for *synXI* assembly | This study - GeneArt |
| pSXI_3_34_O3 | Vector containing synthetic chunk O3 | Chunk propagation for *synXI* assembly | This study - GeneArt |
| pSXI_3_34_O4 | Vector containing synthetic chunk O4 | Chunk propagation for *synXI* assembly | This study - GeneArt |
| pSXI_3_34_O5 | Vector containing synthetic chunk O5 | Chunk propagation for *synXI* assembly | This study - GeneArt |
| pSXI_3_34_P1 | Vector containing synthetic chunk P1 | Chunk propagation for *synXI* assembly | This study - GeneArt |
| pSXI_3_34_P2 | Vector containing synthetic chunk P2 | Chunk propagation for *synXI* assembly | This study - GeneArt |
| pSXI_3_34_P3 | Vector containing synthetic chunk P3 | Chunk propagation for *synXI* assembly | This study - GeneArt |
| pSXI_3_34_P4 | Vector containing synthetic chunk P4 | Chunk propagation for *synXI* assembly | This study - GeneArt |
| pSXI_3_34_P5 | Vector containing synthetic chunk P5 | Chunk propagation for *synXI* assembly | This study - GeneArt |
| pSXI_3_34_Q1 | Vector containing synthetic chunk Q1 | Chunk propagation for *synXI* assembly | This study - GeneArt |
| pSXI_3_34_Q2 | Vector containing synthetic chunk Q2 | Chunk propagation for *synXI* assembly | This study - GeneArt |
| pSXI_3_34_Q3 | Vector containing synthetic chunk Q3 | Chunk propagation for *synXI* assembly | This study - GeneArt |
| pSXI_3_34_Q4 | Vector containing synthetic chunk Q4 | Chunk propagation for *synXI* assembly | This study - GeneArt |
| pSXI_3_34_Q5 | Vector containing synthetic chunk Q5 | Chunk propagation for *synXI* assembly | This study - GeneArt |
| pSXI_3_34_R1 | Vector containing synthetic chunk R1 | Chunk propagation for *synXI* assembly | This study - GeneArt |
| pSXI_3_34_R2 | Vector containing synthetic chunk R2 | Chunk propagation for *synXI* assembly | This study - GeneArt |
| pSXI_3_34_R3 | Vector containing synthetic chunk R3 | Chunk propagation for *synXI* assembly | This study - GeneArt |
| pSXI_3_34_R4 | Vector containing synthetic chunk R4 | Chunk propagation for *synXI* assembly | This study - GeneArt |

| Plasmid | Description | Use | Source |
|---|---|---|---|
| pSXI_3_34_R5 | Vector containing synthetic chunk R5 | Chunk propagation for *synXI* assembly | This study - GeneArt |
| pSXI_3_36_M3 | Modified M3 vector conforming to synXI_3.36 | Chunk propagation for *synXI* assembly | This study |
| pSXI_3_37_M2 | Modified M2 vector conforming to synXI_3.37. Contains CEN11_3_37 | Chunk propagation for *synXI* assembly | This study |
| pUC19 | *E. coli* cloning vector | Generation of *pCEN11** | Norrander *et al.*, 1983 |
| pWS082 | GAP-repair CRISPR/Cas9 gRNA vector | Retargeted to specify CRISPR/Cas9 target | Shaw *et al.*, 2018 |
| pWS158 | GAP-repair CRISPR/Cas9 cas9 vector with *URA3* marker | CRISPR/Cas9 genome editing | Shaw *et al.*, 2018 |
| pWS171 | GAP-repair CRISPR/Cas9 cas9 vector with *LEU2* marker | CRISPR/Cas9 genome editing | Shaw *et al.*, 2018 |
| pXL007 | *LEU2* integration cassette containing *PRAD27*-[TetA-Nuclear Localisation Signal-*GAL4* activation domain]-T*ADH1* Tet-On cassette, a tet$O_7$-*PPHO5-mScarlet-TTDH1* fluorescent reporter cassette and a *LEU2* marker cassette on a *ColE1-kanR* backbone | Tetracycline-inducible *mScarlet* reporter construct | This study |
| pXL008 | *LEU2* integration cassette containing *PRAD27*-[TetA-Nuclear Localisation Signal-*GAL4* activation domain]-T*ADH1* Tet-On cassette, a tet$O_7$-*PPHO5-BFP2-TTDH1* fluorescent reporter cassette and a *LEU2* marker cassette on a *ColE1-kanR* backbone | Tetracycline-inducible *BFP2* reporter construct | This study |
| pXZX353 | *MATα* in pTOPO vector | Generation of pZY412 | Xie *et al.*, 2018 |
| pYKT083 | MoClo-YTK part plasmid: *AmpR-ColE1* | Generation of p*SCW11-cre*-EBD-*kanMX4* | Lee *et al.*, 2015 |
| pYTK003 | MoClo-YTK part plasmid: ConL1 | Generation of p*SCW11-cre*-EBD-*kanMX4* | Lee *et al.*, 2015 |
| pYTK051 | MoClo-YTK part plasmid: T*ENO1* | Generation of p*SCW11-cre*-EBD-*kanMX4* | Lee *et al.*, 2015 |
| pYTK067 | MoClo-YTK part plasmid: ConR1 | Generation of p*SCW11-cre*-EBD-*kanMX4* | Lee *et al.*, 2015 |
| pYTK077 | MoClo-YTK part plasmid: *kanMX4* | Generation of p*SCW11-cre*-EBD-*kanMX4* | Lee *et al.*, 2015 |
| pYTK081 | MoClo-YTK part plasmid: *CEN6/ARS4* | Generation of p*SCW11-cre*-EBD-*kanMX4* | Lee *et al.*, 2015 |
| pZY412 | *MATα* expression vector | Sporulation of homozygous diploid | This study |

***Table S6 - Plasmids used and generated in this study.***

### References

Bakota, L., Brandt, R., and Heinisch, J.J. (2012). Triple mammalian/yeast/bacterial shuttle vectors for single and combined Lentivirus- and Sindbis virus-mediated infections of neurons. Mol. Genet. Genomics *287*, 313–324.

Blount, B.A., Weenink, T., Vasylechko, S., and Ellis, T. (2012). Rational Diversification of a Promoter Providing Fine-Tuned Expression and Orthogonal Regulation for Synthetic Biology. PLoS ONE *7*, e33279. https://doi.org/10.1371/journal.pone.0033279.

Brachmann, C. B., Davies, A., Cost, G.J., Caputo, E., Li, J., Hieter, P., Boeke, J. D. (1998). Designer deletion strains derived from Saccharomyces cerevisiae S288C: a useful set of strains and plasmids for PCR-mediated gene disruption and other applications. Yeast *14(2)*, 115–132.

Cai, Y., Agmon, N., Choi, W.J., Ubide, A., Stracquadanio, G., Caravelli, K., Hao, H., Bader, J.S., and Boeke, J.D. (2015). Intrinsic biocontainment: multiplex genome safeguards combine transcriptional and recombinational control of essential yeast genes. Proc. Natl. Acad. Sci. U. S. A. *112*, 1803–1808.

Huxley, C., Green, E.D., and Dunham, I. (1990). Rapid assessment of S. cerevisiae mating type by PCR. Trends Genet. *6*, 236.

Lee, M.E., DeLoache, W.C., Cervantes, B., and Dueber, J.E. (2015). A Highly Characterized Yeast Toolkit for Modular, Multipart Assembly. ACS Synth. Biol. *4*, 975–986.

Mülleder, M., Capuano, F., Pir, P., Christen, S., Sauer, U., Oliver, S.G., and Ralser, M. (2012). A prototrophic deletion mutant collection for yeast metabolomics and systems biology. Nat. Biotechnol. *30*, 1176–1178.

Norrander, J., Kempe, T. & Messing, J. (1983). Construction of improved M13 vectors using oligodeoxynucleotide-directed mutagenesis. *Gene 26*, 101–106.

Shaw, W.M., Yamauchi, H., Mead, J., Gowers, G.-O.F., Bell, D.J., Öling, D., Larsson, N., Wigglesworth, M., Ladds, G., and Ellis, T. (2019). Engineering a Model Cell for Rational Tuning of GPCR Signaling. Cell *177*, 782–796.e27.

Sikorski, R.S., and Hieter, P. (1989). A system of shuttle vectors and yeast host strains designed for efficient manipulation of DNA in Saccharomyces cerevisiae. Genetics *122*, 19–27.

Winzeler, E.A., Shoemaker, D.D., Astromoff, A., Liang, H., Anderson, K., Andre, B., Bangham, R., Benito, R., Boeke, J.D., Bussey, H., et al. (1999). Functional characterization of the S. cerevisiae genome by gene deletion and parallel analysis. Science *285*, 901–906.

Xie, Z.-X., Mitchell, L.A., Liu, H.-M., Li, B.-Z., Liu, D., Agmon, N., Wu, Y., Li, X., Zhou, X., Li, B., et al. (2018). Rapid and Efficient CRISPR/Cas9-Based Mating-Type Switching of. G3 *8*, 173–183.
